# Cornealia™: automated collective order analysis reveals density-dependent reorganization of donor corneal endothelium

**DOI:** 10.64898/2026.10.21.760745

**Authors:** Majid Rostami, Saad Asim, Mason Steele, Darren Sai Kyauk, Lisa Moul, Samuel Haarman, Addie Richard, Gary Peh, David Moore, Muhammad Rizwan, Jodhbir S. Mehta, Sangyoon J. Han

## Abstract

The corneal endothelium is a non-regenerative monolayer whose health determines transplant success, yet donor tissue is assessed only by single-cell metrics that ignore how the monolayer is organized collectively. A physics-based spring constant *K*, derived from the radial distribution function of cell positions, captures this collective order but has remained limited to semi-manual studies without physical interpretation. Here we present Cornealia™, an automated pipeline for collective-order and intercellular-force analysis. In cultured endothelial cells, collective order declines with passaging and is higher under growth-factor conditions that preserve N-cadherin junctions, differences invisible to cell density and hexagonality. Across 2,177 donor cornea images from two cohorts, collective order, distinct from classical metrics, responds most to mechanical perturbation: corneas with endothelial cell density above 2,400 cells/mm^2^ mostly gain order, whereas less dense ones lose it, on one curve shared by both cohorts. Reconstructed intercellular forces track these changes. This switch lies within the minimum donor densities used by eye banks, suggesting a mechanical basis for these clinical thresholds and establishing collective order as an automatable measure of endothelial quality.

---

The corneal endothelium is a specialized monolayer of cells that maintains corneal transparency by regulating stromal hydration through active fluid transport^1,2^. Unlike most tissues, corneal endothelial cells have very limited proliferative capacity in vivo: cell loss is compensated primarily by enlargement and migration of neighboring cells rather than by regeneration^3,4^. When endothelial cell density falls below a critical threshold corneal edema and vision loss can occur, leaving transplantation as the only effective treatment^5^. In 2024, U.S. eye banks alone supplied tissue for 85,926 corneal grafts worldwide, including 41,558 endothelial and 31,060 penetrating keratoplasties, both of which transplant the donor endothelium ^6,7^, and because donor endothelial cells also possess limited regenerative capacity, the quality of the donor tissue is a major determinant of long-term graft success^8^.

In clinical practice, donor tissue quality is primarily evaluated using specular microscopy-derived metrics such as cell density (CD), coefficient of variation of cell area (CV), and hexagonality (HEX)^9^. Cell density is the reciprocal of mean cell area, and hexagonality counts the fraction of six-sided cells: both are single-cell properties that fail to account for intercellular interactions and many-body spatial correlations within the cell population ^10^. Endothelial pump and barrier function depend on intact cell-cell junctions that physically connect adjacent cells across the monolayer^11,12^. So measuring individual cells without measuring how they are coupled to each other misses the physical basis of tissue function.

Yamamoto *et al*. addressed this gap by introducing a physics-based framework that treats the corneal endothelium as a two-dimensional cellular assembly and quantifies its collective organization through an effective intercellular interaction potential^10^. From the radial distribution function of cell centroids, they derived a spring constant *K* that reflects how tightly cells maintain their preferred spatial arrangement. Unlike CD, CV, or HEX, *K* captures long-range spatial correlations and many-body interactions within the monolayer. In 12 penetrating keratoplasty patients, *K* outperformed conventional morphological metrics in predicting the 24-month cell density trajectory (AUC = 0.85 vs 0.56 for cell density alone).

Despite this result, the concept has not yet seen widespread clinical adoption. The barrier was partly practical: semi-manual segmentation and expert-supervised curve fitting confined the analysis to fewer than forty images, and the spring constant was obtained by fitting a parabola to a narrow window around the potential minimum, an estimate that is sensitive to noise and to the choice of fitting boundaries. Moreover, as noted in the accompanying commentary^13^, the method needs to be validated across different culture conditions, serial passages, and transplantation techniques before it can enter clinical practice. Beyond these practical barriers, more conceptual exploration is also required. Whether *K* is genuinely distinct from cell density at population scale remained untested. Moreover, since cell density is already an established biomarker, how collective order and density change together under perturbation was never characterized. The spring constant itself stayed an abstract statistical quantity, leaving open what physical mechanism underlies it and whether it relates to the junctional machinery that physically couples neighbouring cells.

Here we present Cornealia, a fully automated, open-source, CPU-only pipeline for collective order and intercellular force analysis of corneal endothelial monolayers. Training-free segmentation (CellADE) extracts cell centroids and generalizes across specular microscopes without retraining, including instruments with non-square pixels. Rather than fitting a narrow parabolic window around the potential minimum, Cornealia fits the entire radial distribution function with a physically motivated model and derives a new spring constant, *K*_ds_, from the resulting smooth potential; the same fit acts as a quality gate that rejects images whose spatial statistics depart from the expected form. The fitted potential is then converted into a truss model that gives the force each cell exerts on its neighbours. The pipeline is integrated into CelLAB, a desktop platform that requires no programming experience.

In cultured endothelial cells, *K*_ds_ resolves growth-factor conditions that cell density and hexagonality cannot distinguish and declines with serial passaging in cells from two independent laboratories, establishing that it tracks known biology. Applying Cornealia to over 2,000 donor cornea images from two independent clinical cohorts, we show that *K*_ds_ is distinct from classical metrics at baseline and is the metric that responds most to mechanical perturbation. How order changes depends on baseline density: once regression to the mean is corrected, corneas with endothelial cell density above 2,400 cells/mm^2^, within the range of minimum donor densities used by eye banks, mostly gain order, whereas less dense ones increasingly lose it, along one density curve that a second cohort extends to lower densities. Reconstructed intercellular forces track collective order and its change; in culture, conditions that preserve the junctional proteins N-cadherin and ZO-1 produce stronger force networks and higher order than EGF alone. Together, these results es-tablish collective order as a distinct, physically interpretable biomarker of corneal tissue and identify active intercellular force generation as a candidate mechanism for how the monolayer reorganizes after perturbation.

## Results

### Full-curve fitting yields a noise-robust spring constant

Cornealia processes specular microscopy images (Fig. 1a) through four stages: 1) preprocessing to isolate the image area from device exports that include interface elements, text, and overlays, such as those of the CellChek D (Konan Medical), 2) cell segmentation and centroid extraction (Fig. 1b), 3) extraction of classical parameters such as cell density, CV and hexagonality, together with collective order, and 4) intercellular force analysis.

**Figure 1.**
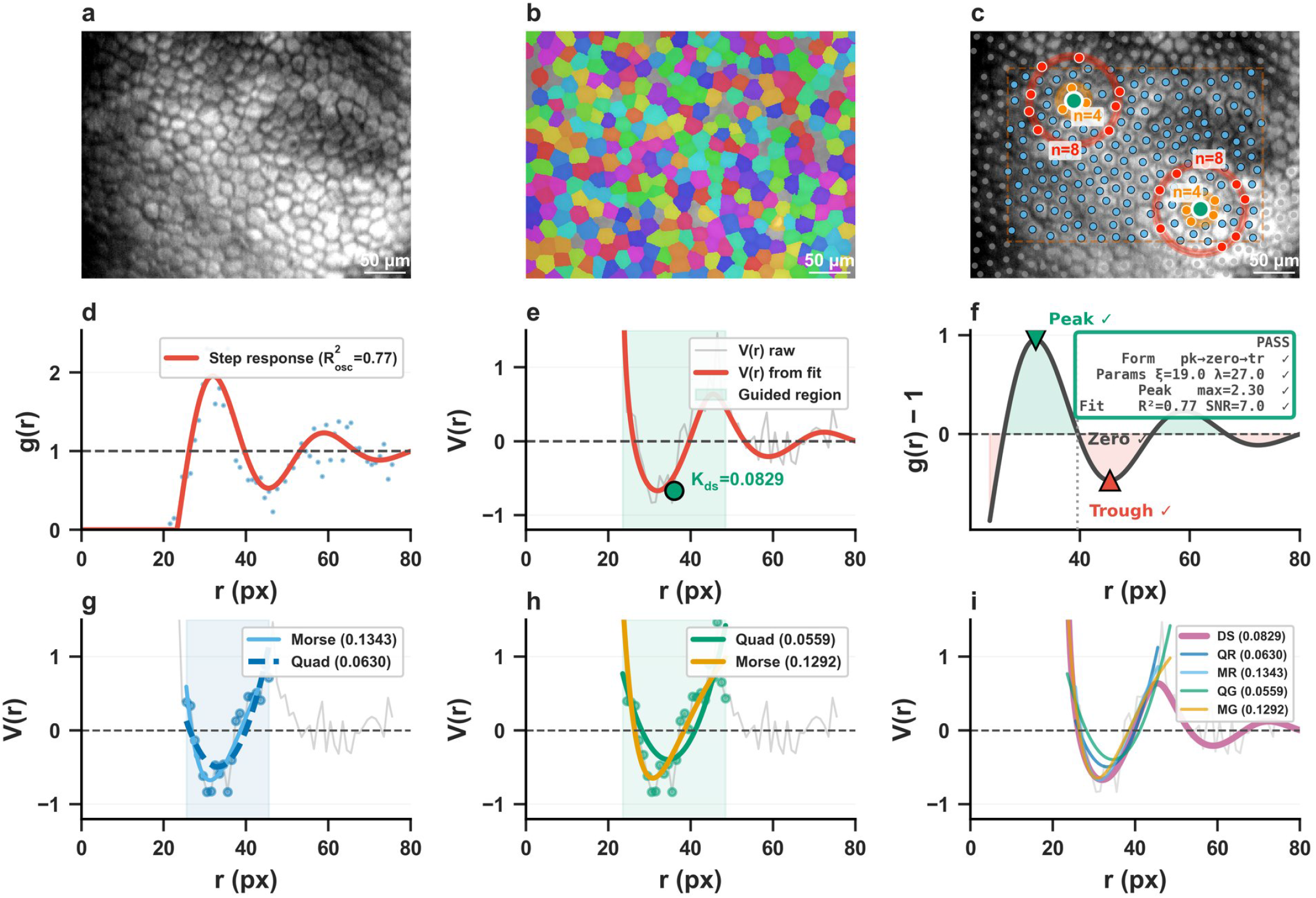
Cornealia pipeline overview. Row 1: (a) Raw specular microscopy image (contrast adjusted for print). (b) CellADE segmentation overlay. (c) Detected centroids with interior region (dashed box); for two example cells, the neighbours in the first and second shells are highlighted, as counted in *g*(*r*). Scale bars, 50 *µ*m. **Row 2, Quality gate:** (d) Step response fit to *g*(*r*). (e) Smooth *V* (*r*) with guided fitting region (green) and *K*_ds_ = *V* ^*′′*^(*r*_0_). (f) Oscillatory form check showing the three diagnostic features (peak, zero crossing, trough) with composite reliability assessment. **Row 3, Five extraction methods:** (g) Quadratic (dashed) and Morse (solid) fits on raw region (argmin ± 10 px). (h) Same models on analytically guided region.

Segmentation is performed by CellADE (see “Training-free segmentation generalizes to clinical images” below; Supplementary Section S7), from which cell centroids are extracted (Fig. 1c).Collective order follows the framework of Yamamoto *et al*. ^10^, which treats the corneal endothelium as a two-dimensional colloidal assembly. From the centroids, the radial distribution function *g*(*r*) gives the probability of finding a neighbour at distance *r* relative to a random arrangement (Fig. 1d); unlike cell density and its coefficient of variation, it takes all pair correlations into account and therefore captures long-range order. Its inverse Boltzmann transform, *V* (*r*) = − ln *g*(*r*), is the effective intercellular potential, termed the potential of mean force, a free energy that includes many-body interactions. The curvature of *V* at its first minimum, *K* = *V* ^′′^(*r*_0_), where *r*_0_ is the position of the first peak of *g*(*r*), reflects how tightly cells hold their preferred spacing (Fig. 1e; Methods).

Yamamoto *et al*. extract the spring constant by fitting a symmetric parabola to a narrow window around the minimum of *V* (*r*). This local approach has two limitations: the enforced symmetry misses any asymmetry around the minimum, and the narrow window discards the rest of the curve, leaving the fit with no global constraint against the noise that clinical *V* (*r*) inherits from *g*(*r*). To address both limitations, we replace the local approach with a physics-informed model that fits the entire *g*(*r*) curve. The radial distribution function behaves like the response of a damped oscillator: it is zero inside the exclusion zone, overshoots at the nearest-neighbour shell, and relaxes to unity through decaying oscillations, which an underdamped step response captures with five parameters (Fig. 1d, Eq. 1). *V* (*r*) is then obtained from the parametric *g*(*r*) to find the curvature:

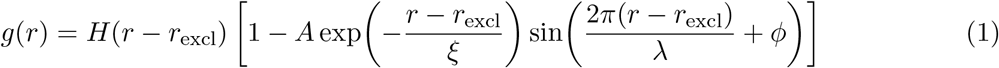

A single fit then serves three purposes. It gives a smooth analytical curve whose curvature at the first minimum yields the spring constant *K*_ds_ (Fig. 1e). It screens out unreliable images through four checks (oscillatory form, parameter plausibility, peak height, and fit quality), so that a curvature is reported only where a preferred spacing exists (Fig. 1f). And it locates the potential well analytically, supplying fitting boundaries for the alternative extraction methods compared below, which therefore no longer depend on finding a minimum in noisy data.

To evaluate the robustness and accuracy of *K*_ds_, we compare it with four parametric estimators on the same dataset. Two models are used: the quadratic model (the symmetric parabola of Yamamoto *et al*.) and the Morse potential, which captures possible asymmetry on the two sides of the minimum. Each is fitted either on the raw region, a fixed window of ±10 px around the lowest point of the raw *V* (*r*) as in the original method, or on the guided region derived from our step response fit, which prevents a single noisy bin from displacing the fitting window (Fig. 1g,h). This 2×2 design separates the effects of the fitting model and of boundary selection, and its four parametric estimators, together with *K*_ds_, make up the five methods shown in Fig. 1i and compared throughout.

### Training-free segmentation generalizes to clinical images

Instead of precise boundary segmentation, computing the collective order biomarker *K* requires reliable cell centroid positions. This motivates CellADE (Cell Adaptive Detection Engine), which, unlike deep-learning segmenters, runs on CPU alone and needs no training or re-training. Moreover, its underlying parameters have direct physical meaning, which makes it generalize more easily to new datasets (Supplementary Section S7; Supplementary Fig. S7). When automatic scale estimation underperforms on atypical images, adjusting three interpretable parameters, (i) the LoG scale range for cell size, (ii) the LoG detection threshold for faint cells and (iii) the FCM membership threshold for dim pixels, recovers accuracy, whereas a deep-learning segmenter would require collecting annotations and retraining model weights. These properties make CellADE directly embeddable in clinical and research pipelines such as Cornealia. Figure 2a compares CellADE and Cellpose-SAM detections on representative clinical images.

**Figure 2.**
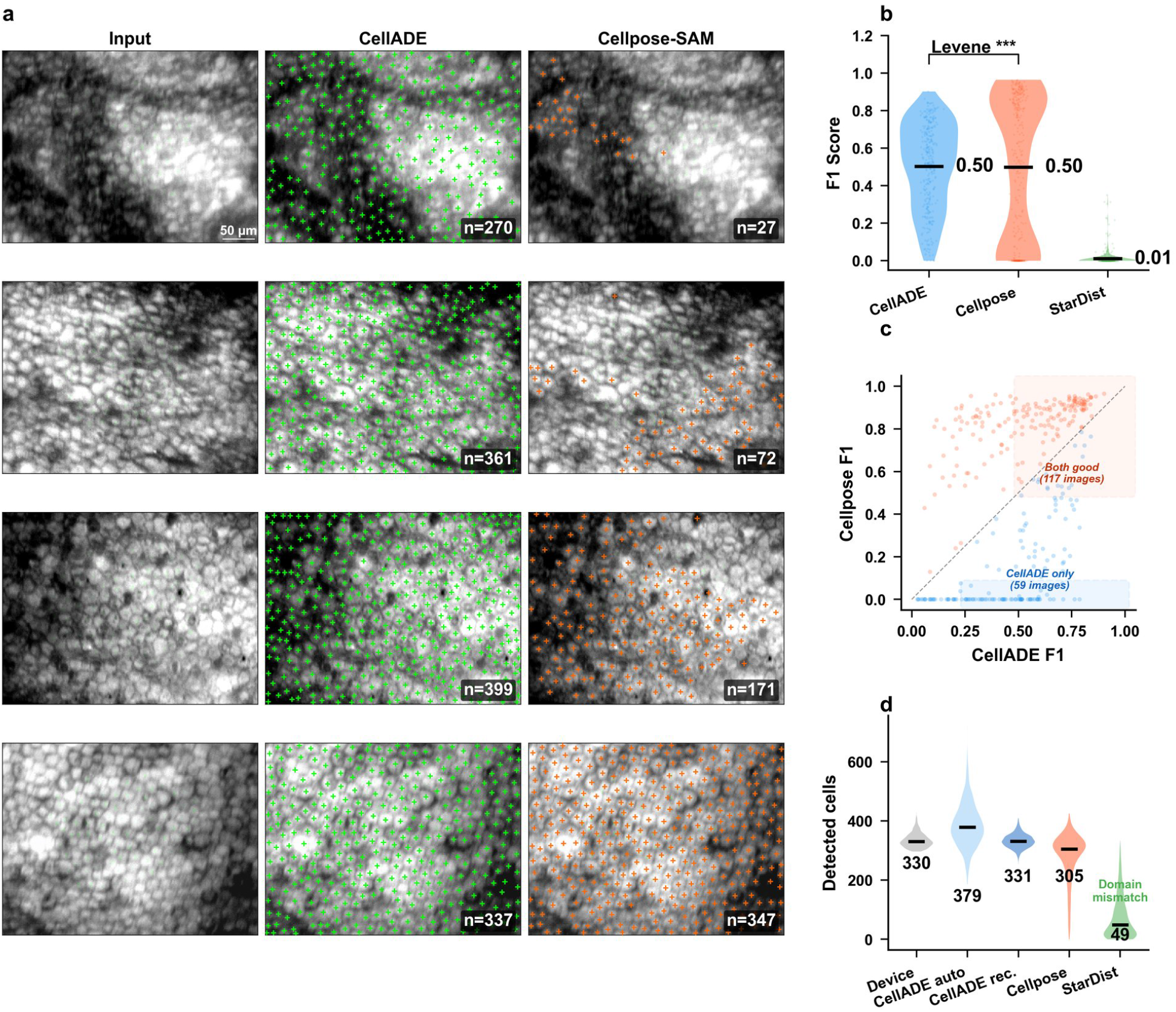
CellADE segmentation performance. **(a)** Representative clinical images with CellADE (green) and Cellpose-SAM (orange) centroid detections. Bottom row (blue tint): comparable performance. Scale bar: 50 µm; contrast adjusted for print. **(b)** F1 distribution on the Daniel *et al*. benchmark (324 annotated images, 15 px match radius); black line and label mark the mean per method. CellADE achieves comparable mean F1 with significantly lower variance (Levene’s test, *p <* 10^*−*31^); Cellpose-SAM returned zero detections on 25% of images.**(c)** Per-image F1 comparison. Blue: CellADE superior; orange: Cellpose-SAM superior. Shaded boxes mark images where both methods perform well (F1 *>* 0.5 for both, 117 images) and where only CellADE detects cells (CellADE F1 *>* 0.25, Cellpose F1 *<* 0.05, 59 images). **(d)** Cell counts on 1,606 clinical images for the device (density × imaged area, a clinical reference), CellADE with automatic scale estimation and after recalibration (rec.), Cellpose-SAM and StarDist; numbers give the median per method (Table 1).

On the Daniel *et al*. ^14^ corneal benchmark (324 annotated images), CellADE achieved a mean F1 of 0.50, comparable to Cellpose-SAM ^15^ at 0.50 (without corneal-specific fine-tuning), but with significantly lower variance (Levene’s test, *p <* 10^−31^), higher recall (0.58 vs 0.47), and substantially better cell density correlation (*R*^2^ = 0.71 vs 0.51), the metrics most relevant to downstream *g*(*r*) and centroids (Table 1; Fig. 2b). Cellpose-SAM returned zero detections on 80 images (25%); CellADE returned none. The per-image comparison shows that both methods perform well on 117 images, while on 59 images only CellADE detects cells (Fig. 2c). As a negative control, StarDist^16^, pretrained on fluorescence microscopy, detected virtually no cells (F1 = 0.011), confirming that its training domain does not transfer to specular microscopy. Notably, Daniel *et al*. ^14^ reported higher cell density correlation (*R*^2^ = 0.96) using a U-Net trained on 158 corneal images, but the trained model weights were not publicly released, precluding independent verification on our clinical dataset, testing of its generalizability to other imaging conditions, and integration into an automated pipeline.

**Table 1.** Cell detection on two datasets. Top: centroid detection on the Daniel *et al*. ^14^ dataset (324 annotated images, 15-pixel match radius). Bottom: cell counts on 1,606 clinical images against the count implied by the device-reported density (CD × imaged area), a clinical reference rather than ground truth. Deep learning methods applied without corneal-specific fine-tuning.

| *Daniel et al. benchmark (324 annotated images)* |  |  |  |  |  |  |
| --- | --- | --- | --- | --- | --- | --- |
| Method | F1 | Precision | Recall | CD $R^2$ | Time (s) | Requirements |
| **CellADE (ours)** | **0.504** | 0.471 | **0.579** | **0.708** | 3.0 | 1 CPU core |
| Cellpose-SAM | 0.500 | **0.649** | 0.469 | 0.507 | 8.0 | GPU (8 GB) |
| StarDist | 0.011 | 0.069 | 0.007 | 0.014 | — | GPU |
| U-Net<sup>14</sup> (trained) | — | — | — | 0.96 | — | GPU, 158 train. images |
| *Clinical images, agreement with the device (1,606 images)* |  |  |  |  |  |  |
| Method | Median cells | Median ratio | Bias (cells) | 95% LoA | MAPE (%) | $R^2$ |
| Device ($CD \times \text{area}$) | 330 | — | — | — | — | — |
| CellADE, automatic scale | 379 | 1.13 | +50.7 | −84 to 185 | 16.0 | 0.09 |
| **CellADE, recalibrated** | **331** | **1.00** | **−1.9** | **−44 to 40** | **3.9** | **0.34** |
| Cellpose-SAM | 305 | 0.93 | −52.2 | −202 to 98 | 7.5 | 0.05 |
| StarDist | 49 | 0.15 | −262.1 | −406 to −118 | 85.5 | 0.00 |

On the 1,606 donor cornea images from the Singapore National Eye Centre with a device-reported density, we compared each method’s cell count with the count implied by the density reported by the specular microscope (CD × imaged area), used as a clinical reference (Table 1; Fig. 2d). Because the CellChek analysis uses only a set of central cells, this count extrapolated to the full image is not exactly comparable, but it is accurate enough to estimate overall accuracy. With automatic scale estimation, CellADE overcounted (median 379 cells against 330 for the device; median absolute error 16%), Cellpose-SAM undercounted with a long tail toward zero (median 305; 7.5%), and StarDist failed (median 49). Recalibrating the LoG scale range (*σ* = 8–12 px) and lowering the LoG threshold recovered agreement: median 331 cells, bias −1.9 cells (95% limits of agreement −44 to 40), median absolute error 3.9% (Table 1; recalibration in Supplementary Section S3.2 and Supplementary Fig. S3b).

### *K*_ds_ is more accurate and stable than local-window fits

On 10,000 synthetic *g*(*r*) curves spanning 10 signal-to-noise ratios (SNR 1.5–50), the gate almost never admitted a curve without genuine oscillatory structure (specificity ≥ 98.5% across all non-oscillatory signal types), while the fraction of genuine curves it correctly admitted rose steadily with signal quality, from 59% at SNR = 5 to 84% at SNR = 50 (Fig. 3a). Comparing the accuracy of recovering the ground-truth *K* values, *K*_ds_ outperforms the others at most noise levels tested (Fig. 3b), reaching a median error of 2.6% at SNR = 50 and remaining below 14% at SNR = 10. The quadratic raw method of Yamamoto *et al*. has an approximately 28% error even on nearly clean data, because of its enforced symmetry. Widening the fitting region to the analytically guided boundaries improved the Morse fit, which can follow the asymmetry, but degraded the quadratic fit (Fig. 3c–e).

**Figure 3.**
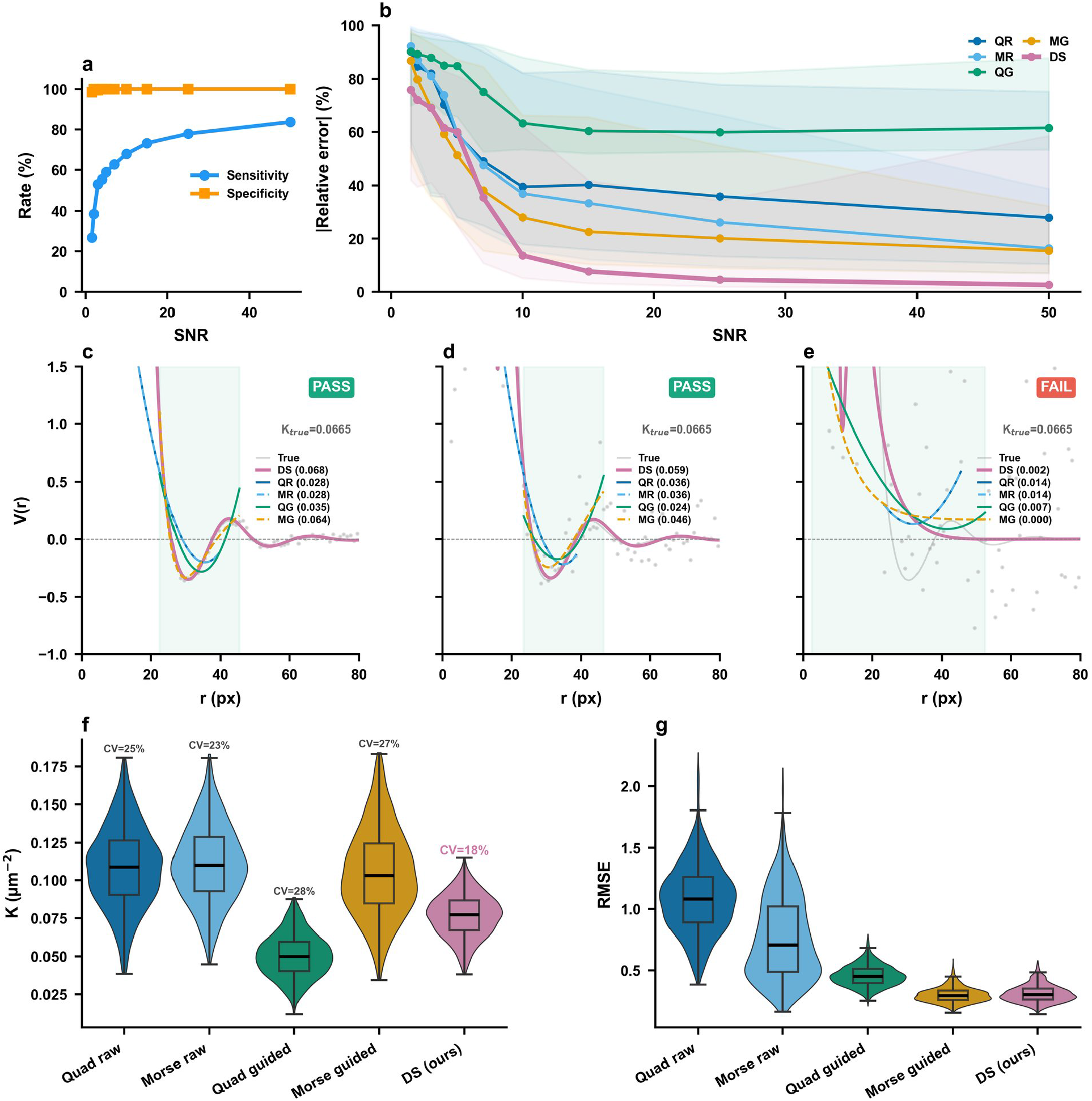
Method validation on synthetic and clinical data. **(a)** Quality gate sensitivity and specificity versus SNR on 10,000 synthetic *g*(*r*) curves and 2,000 non-oscillatory controls.**(b)** Median absolute relative error of *K* versus SNR per method with IQR bands; *K*_ds_ reaches 2.6% at high SNR. **(c–e)** Representative *V* (*r*) fits at SNR = 50, 7 and 2 with all five methods overlaid; ground truth in grey, guided region shaded green. PASS/FAIL badge indicates quality gate verdict; at SNR = 2 the curve is correctly rejected. **(f)** *K* distributions on 1,583 reliable clinical images, with coefficient of variation annotated per method. **(g)** RMSE between the raw *V* (*r*) points and each method’s fitted curve, evaluated for all five methods on the same guided well region. Method colours are consistent across all panels and with Fig. 1.

Testing 1,583 reliable clinical images, *K*_ds_ achieved the lowest coefficient of variation among all five methods (CV = 18%, vs. 23–28% for alternatives; Fig. 3f), indicating the most stable *K* estimates across the cohort. We assessed fit quality by the root-mean-square error (RMSE), the typical vertical distance between a fitted curve and the data points it describes, measured between the raw *V* (*r*) points and each method’s fitted curve on the same guided well region for fairness (Fig. 3g). The Morse guided fit attained the lowest median RMSE (0.293), as expected since it minimizes squared error on precisely that region by construction; *K*_ds_ matched it closely (0.302) despite being fitted to the entire *g*(*r*) curve, and both are well below the quadratic guided fit (0.452) and the raw-region methods (0.709 and 1.081).

### *K*_ds_ declines with passaging in cultured cells

Before applying *K*_ds_ to clinical tissue, we tested whether it tracks a known biological loss of organization. Primary rabbit corneal endothelial cells were cultured using the dual-media system of Peh *et al*. ^17^ and imaged at each passage: three culture lines, each established from one cornea and followed from P0 to P4 (137 reliable images, 10×; Fig. 4a). *K*_ds_ followed a clean exponential decay, *K*_ds_ = 0.057 *e*^−0.46*P*^ (*R*^2^ = 0.996; Fig. 4c), losing 37% of the remaining order at every passage and leaving one-sixth of the P0 value by P4. The decay was reproducible across the three culture lines (decay rate 0.41–0.49 per passage; mean 0.45 ± 0.04), with *K*_ds_ falling at every passage in every line. Cell density fell in parallel, from 2,686 to 563 cells/mm^2^ (Fig. 4e), while hexagonality declined far less, from 46.6% to 38.3%, a change that did not reach significance between consecutive passages at the level of culture lines (Fig. 4d). The population-averaged *g*(*r*) shows how order is lost: the first peak, the relative number of neighbours at the preferred spacing, falls while the preferred spacing itself moves outward, from 19.1 *µ*m at P0 to 21.2, 29.0 and 37.1 *µ*m at P1, P2 and P4 (Fig. 4b).

**Figure 4.**
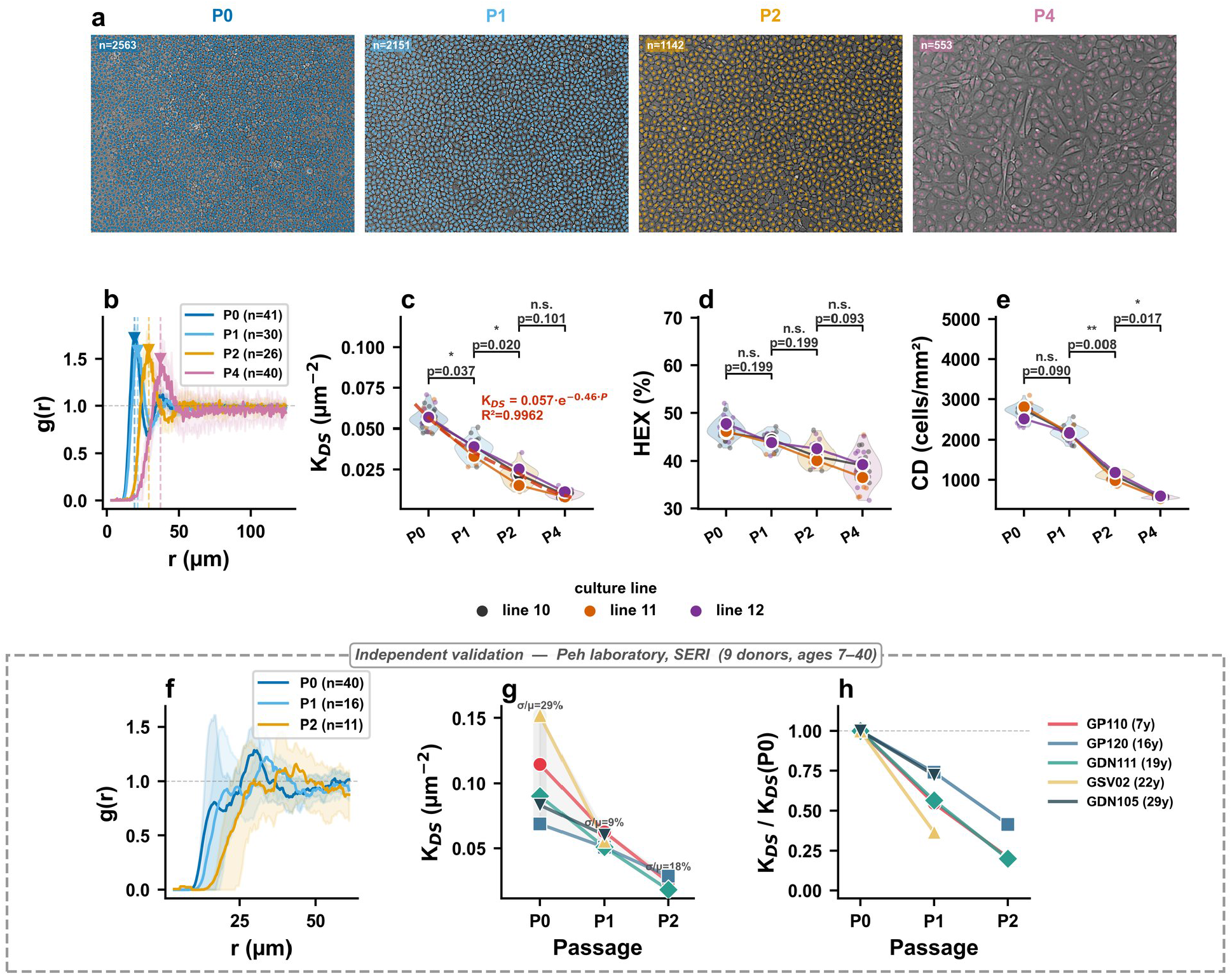
Collective order declines with passaging in cultured corneal endothelial cells. **(a)** Representative segmented phase-contrast images of primary rabbit cells at passages P0, P1, P2 and P4, with detected cell counts annotated; scale bar, 50 *µ*m. **(b)** Population-averaged *g*(*r*) per passage with IQR bands; triangles and dashed lines mark the first-peak position, which moves outward with passage number. **(c–e)** *K*_ds_, hexagonality and cell density per passage, shown as SuperPlots ^18^: small points are individual images coloured by culture line, large points are the mean of each line, joined across passages, and violins show the image-level distribution. Brackets give paired *t*-test *P* values on the line means (*n* = 3 culture lines), Bonferroni-corrected for the three adjacent comparisons. The dashed curve in (c) is the exponential fit *K*_ds_ = 0.057 *e*^*−*0.46*P*^ . **(f–h)** Independent validation in primary human cultures from a second laboratory (Peh, SERI): **(f)** population-averaged *g*(*r*) per passage; **(g)** *K*_ds_ per donor across passages, with the inter-donor coefficient of variation annotated at each passage; **(h)** the same trajectories normalized to each donor’s own P0 value.

To assess generalizability, we analyzed primary human cultures from a second laboratory (Peh, SERI), comprising nine donors aged 7–40 years, five of them followed across two to four passages. Population-averaged *g*(*r*) curves pooled by passage reproduced the rightward peak shift and amplitude reduction seen in the rabbit dataset (Fig. 4f). Every donor with multi-passage data declined monotonically in *K*_ds_, from a mean of 0.102 *µ*m^−2^ at P0 to 0.056 at P1 and 0.023 at P2 (Fig. 4g). Normalizing each trajectory to its own P0 baseline shows that the rate varies between donors, with 37–74% of the initial value preserved after one passage, and without dependence on donor age (Fig. 4h).

### *K*_ds_ is distinct from density and responds most to stripping

The primary clinical cohort comprises specular microscopy images of donor corneas from the Singapore National Eye Centre (CellChek D), each imaged before tissue preparation (Before Stripping, BS) and after the Descemet’s membrane–endothelium complex was stripped from the stroma for endothelial keratoplasty (After Stripping, AS), yielding 782 reliable BS/AS pairs (Methods). *K*_ds_ is only moderately correlated with cell density (*r* = 0.53, *P* = 1 × 10^−106^) and weakly with hexagonality (*r* = 0.34, *P* = 2 × 10^−40^) (Fig. 5a,b). The boxplots beneath the scatters confirm that tissues with the same cell density or hexagonality span a wide range of *K*_ds_ values, meaning that two tissues with identical numbers of cells per unit area, or the same fraction of six-sided cells, may have very different levels of structural order.

**Figure 5.**
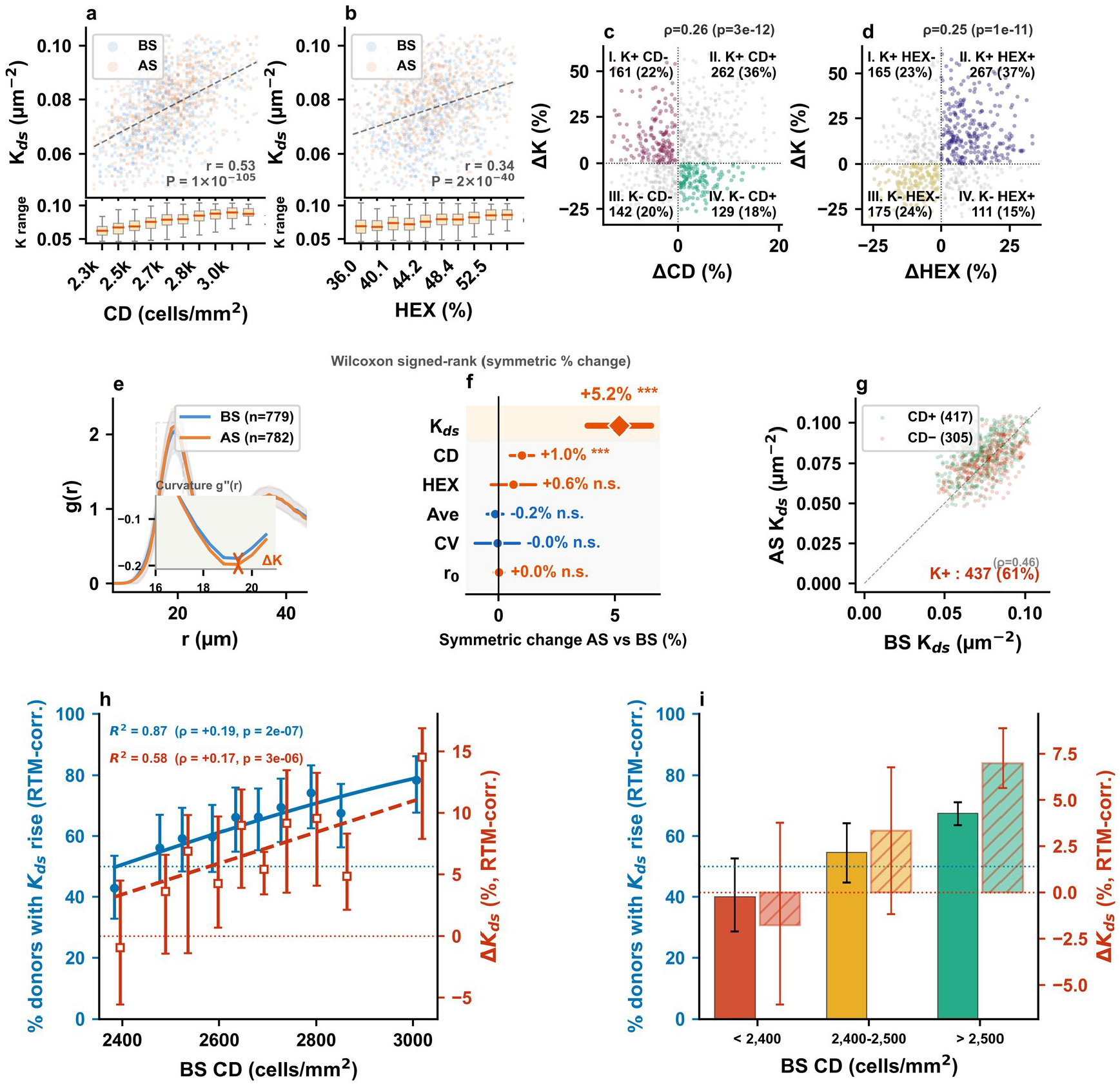
Collective order rises after tissue preparation, more often and more strongly in corneas with higher endothelial cell density. **(a**,**b)** *K*_ds_ versus cell density and hexagonality for reliable images, after a 2–98% clip on both axes (*n* = 1,458 and 1,443), with Pearson *r* and *P* annotated; boxplots below show the *K*_ds_ range within each bin of the *x* variable. **(c**,**d)** Paired percentage change in *K*_ds_ against change in cell density and in hexagonality, with counts and percentages per quadrant and Spearman *ρ*. **(e)** Population-averaged *g*(*r*) before (BS) and after (AS) stripping with IQR bands; inset shows the curvature *g*^*′′*^(*r*) at the first peak, sharper after stripping. **(f)** Mean symmetric percentage change after stripping, (AS − BS)*/*[(AS+BS)*/*2], with 95% confidence intervals for all metrics; *P* values from Wilcoxon signed-rank tests on the symmetric changes. **(g)** Paired *K*_ds_ before and after stripping, coloured by the direction of density change; the dashed line marks no change, K+ gives the number of donors whose *K*_ds_ increased (437 of 722, 61%), and *ρ* is the correlation between BS and AS values. **(h**,**i)** Change in *K*_ds_ after stripping, corrected for regression to the mean with *k* = 1, measured from split images (Supplementary Section S5). Left axis (blue): percentage of donors whose *K*_ds_ rose, with Wilson 95% confidence intervals. Right axis (red): median percentage change, computed on the log scale, with bootstrap 95% confidence intervals. **(h)** Per decile of baseline cell density, with a logistic fit (direction) and a linear fit (size); *ρ, P* and the *R*^2^ of each fit across deciles are annotated. **(i)** In the three eye-bank density bands: below 2,400 cells/mm^2^, 2,400–2,500 cells/mm^2^ (commonly required for endothelial grafts) and above 2,500 cells/mm^2^.

After stripping, collective order and cell density change only loosely together: 56% of donors moved both in the same direction and 40% in opposite directions (*ρ* = +0.26; Fig. 5c). Because *K*_ds_ and cell density are computed from the same centroids, we tested whether shared segmentation error could produce this correlation by adding missed cells or false detections to each image; in both cases the correlation weakened rather than strengthened, so it does not arise from segmentation error (Methods; Supplementary Section S6 and Supplementary Fig. S6). Hexagonality behaves similarly, moving with collective order in 61% of donors and against it in 38% (*ρ* = 0.25; Fig. 5d). Altogether, this shows that the three metrics report different aspects of tissue organization, and establishes *K*_ds_ as a distinct biomarker.

At the population level, we averaged all *g*(*r*) curves grouped by stripping status (BS or AS); AS cases show a slightly sharpened first peak (Fig. 5e and inset). Averaged across paired donors, *K*_ds_ changed the most of all metrics after stripping: the symmetric percentage change, taken relative to the mean of the BS and AS values, was +5.2% for *K*_ds_ (*P* = 4 × 10^−12^), against +1.0% for cell density (*P* = 2 × 10^−4^), with no significant change in hexagonality, cell-size variation, mean cell area or *r*_0_ (Fig. 5f). At the donor level, *K*_ds_ increased in 61% of donors (437 of 722; binomial test, *P* = 2 × 10^−8^), and a donor’s *K*_ds_ before stripping is moderately predictive of its value afterwards (*ρ* = 0.46; Fig. 5g). Collective order is therefore the metric most responsive to stripping; which tissues gain it, and by how much, is examined next.

### Baseline density sets how order changes after stripping

Having established that *K*_ds_ is distinct from cell density at baseline, we now ask how they change together after stripping. Each donor is measured once before and once after, with no control group, formally a single-arm pre–post design. In this design, regressing a variable’s change against its own baseline is contaminated by mathematical coupling and regression to the mean^19,20^, artifacts that produce a spurious negative slope even when the intervention has no differential effect: samples with higher initial values appear to decrease and those with lower initial values appear to increase, converging on the global mean from both sides (see Methods).

We therefore use a cross-variable design: we bin donors by baseline cell density and measure the change in *K*_ds_ (Fig. 5h), using the 750 reliable pairs that remain after trimming the 1st and 99th percentiles of *K*_ds_ (Table 2). Because the two share 28% of their baseline variance (*r* = 0.53), the artifact can still leak from one variable into the other, so each donor’s change is first corrected for regression to the mean (Methods; Supplementary Section S5 and Supplementary Fig. S5). The correction needs one constant, *k*, which we measured directly by comparing the two halves of each image as two simultaneous reads of the same tissue: *k* ≈ 1 for *K*_ds_, the classical correction^21,22^. After correction, the own-baseline artifact is removed (Supplementary Fig. S5d). Each analysis reports two measures: the direction of change (the percentage of donors whose value rose) and its size (the median percentage change, on the log scale).

**Table 2.** Sample sizes of the primary cohort. Rows above the line are successive filtering steps; rows below are analysis-specific subsets of the 782 reliable pairs.

| Step | $n$ | Definition | Used in |
| --- | --- | --- | --- |
| Images acquired | 1,607 | all specular images from SNEC | Fig. 2d |
| Images analysed | 1,585 | 22 clinician-flagged images excluded (792 BS, 793 AS) | — |
| Images passing the gate | 1,583 | step-response quality gate | Fig. 3f,g |
| BS/AS pairs | 784 | matched by image name (781 donors) | — |
| Reliable pairs | 782 | both images pass the gate (779 donors); SIMEX uses one pair per donor (779) | Fig. 5e,f; Fig. 6; Supp. Fig. S6; Supp. Fig. S4 |
| Paired scatter | 774 | top 1% of $K_{ds}$ (BS) removed | Fig. 6g |
| Density bins and bands | 750 | 1–99% trim on $K_{ds}$ (BS and AS); 753 in Supp. Fig. S5 | Fig. 5h,i; Supp. Fig. S5 |
| Change scatter | 718 | 2–98% clip on $\Delta K\%$ and $\Delta CD\%$ | Fig. 5c |
| Paired $K$ scatter | 722 | 2–98% clip on $K_{BS}$ and $K_{AS}$ | Fig. 5g |

Binning on baseline cell density and measuring the corrected change in collective order, both measures rise with density (Fig. 5h): the share of donors whose *K*_ds_ rose increased from 43% in the sparsest decile (2,072–2,430 cells/mm^2^) to 78% in the densest (2,913–3,303 cells/mm^2^), crossing 50% near 2,400 cells/mm^2^ (*ρ* = 0.19, *P* = 2 × 10^−7^; *R*^2^ = 0.87 across deciles), and the median change grew from −0.9% to +14.5% (*ρ* = 0.17, *P* = 3 × 10^−6^; *R*^2^ = 0.58). Baseline density thus describes the population trend well, although individual donors vary widely around it.

This crossing falls within the range of minimum donor densities used by eye banks. Minimum donor densities typically range from 2,000 to 2,500 cells/mm^2^ ^23^, with 2,400–2,500 cells/mm^2^ commonly required for endothelial grafts^24,25^. Grouping donors by these thresholds (Fig. 5i), corneas with endothelial cell density below 2,400 cells/mm^2^ gained order in only 40% of cases (median change −1.8%; *n* = 60), those at 2,400–2,500 in 55% (+3.3%; *n* = 97), and those above 2,500 in 67% (+7.0%; *n* = 593).

### Intercellular forces track collective order and its change

Collective order rises after stripping, more often and more strongly in corneas with higher endothelial cell density. Collective order, however, still remains an abstract concept. To gain insight into its underlying physics, we convert the centroid field into a field of effective intercellular forces using a Delaunay truss: each cell is linked to its immediate neighbours (Fig. 6a), and each link carries a scalar force *F* (*d*_*ij*_) = *g*^′^(*d*_*ij*_)*/g*(*d*_*ij*_), the negative gradient of the potential of mean force (in units of *k*_*B*_*T* per µm), evaluated at the neighbour distance, compressive when the pair sits closer than the equilibrium spacing *r*_0_ and tensile when it sits further apart. These forces are inferred from cell positions, not measured mechanically. Summing these contributions as vectors gives the residual force on each node, a measure of how far that cell is from local mechanical equilibrium. Looking at individual samples, an interesting relationship emerges: tissues with low collective order have low spatially averaged force (Fig. 6b), while tissues with high *K*_ds_ carry larger forces (Fig. 6c), and the per-node distributions separate cleanly (Fig. 6d).

**Figure 6.**
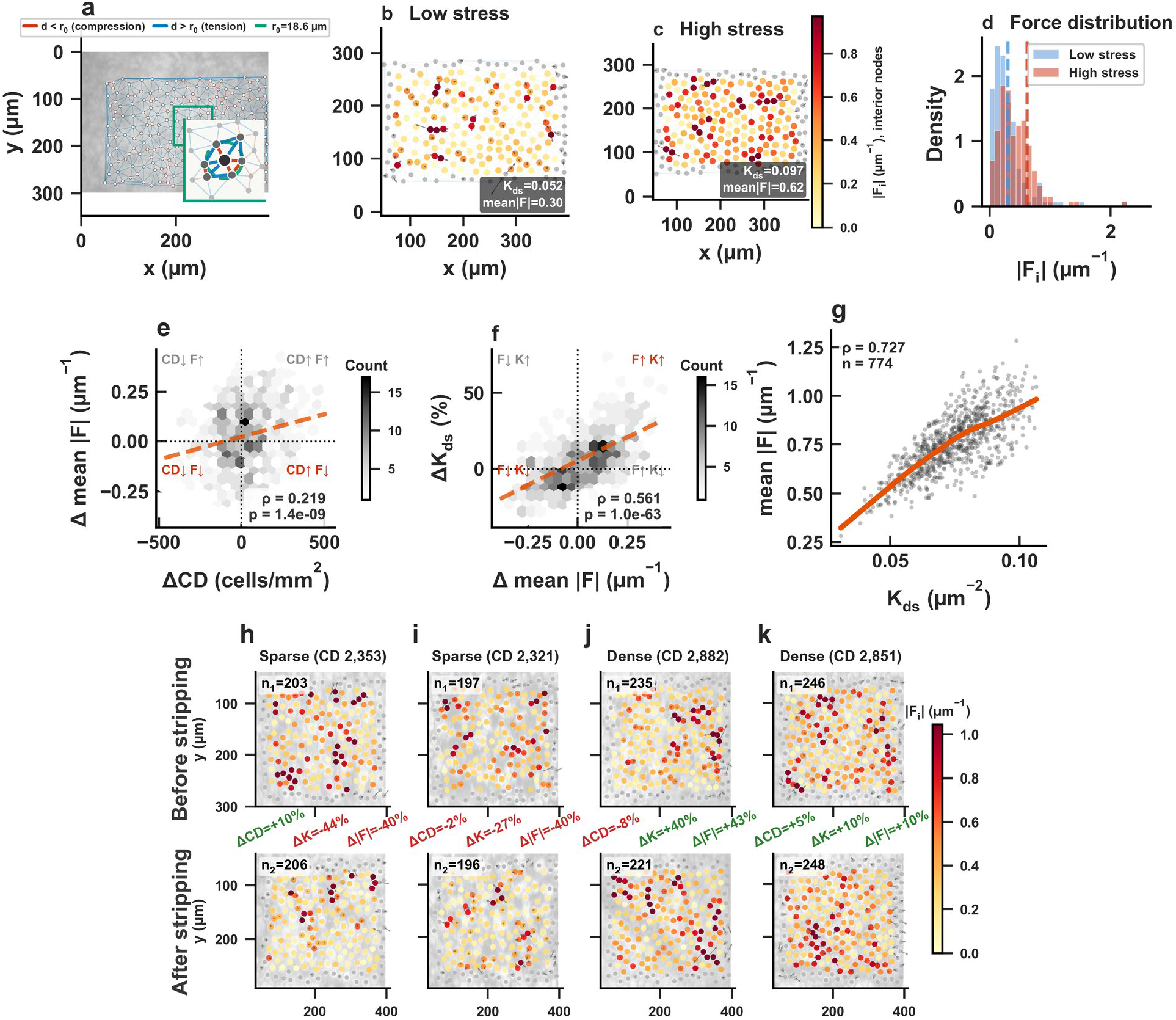
Intercellular forces track the change in collective order. **(a)** Delaunay truss built from cell centroids of one donor, with edges coloured by whether the neighbour distance falls below the equilibrium spacing *r*_0_ (red, compression) or above it (blue, tension); inset shows the force construction at a single node. **(b**,**c)** Residual force fields for a low-stress and a high-stress tissue; interior nodes are coloured by force magnitude |*F*_*i*_| on a shared scale capped at the 95th percentile, and nodes within one cell spacing of the image border, whose neighbourhoods are incomplete, are shown in grey; arrows are the residual force vectors, and *K*_ds_ and mean|*F* | are annotated. **(d)** Distributions of per-node force magnitude |*F*_*i*_| of interior nodes for the two tissues in (b,c); dashed lines mark the means. **(e)** Paired change in mean|*F* | against change in cell density and **(f)** paired percentage change in *K*_ds_ against change in mean|*F* |, as hexbin density with Spearman *ρ*. **(g)** Cross-sectional relationship between *K*_ds_ and mean|*F* | across paired donors (top 1% of baseline *K*_ds_ removed; *n* = 774), with LOWESS fit. **(h–k)** Two donors with lower endothelial cell density that lost order (h, i) and two with higher density that gained it (j, k), selected at the medians of baseline cell density and *K*_ds_, before (top) and after (bottom) stripping, coloured as in (b,c) on one scale shared by all eight fields; changes in cell density, *K*_ds_ and mean|*F* | are annotated between the rows.

Across paired donors, force change follows density change only weakly (*ρ* = +0.22; Fig. 6e), as the change in *K*_ds_ does (Fig. 5c), while changes in force and in collective order follow each other closely (*ρ* = 0.56; Fig. 6f): tissues gain or lose force and order together. Cross-sectionally, *K*_ds_ and mean|*F* | track each other closely (*ρ* = 0.73, *n* = 774; Fig. 6g). Both are computed from the same fitted *g*(*r*), but mean|*F* | additionally uses each cell’s actual neighbour distances and directions: the forces are summed as vectors at each node and averaged over the field, so it shares its basis with collective order while adding spatial and more directly mechanical information. Fig. 6h–k shows two donors with lower endothelial cell density that lost order and two with higher density that gained it, before and after stripping.

### A second cohort extends the density trend to lower densities

A second cohort of 570 specular microscopy images from a different eye bank (Lions VisionGift, Tampa, FL), acquired with a different instrument (HAI Labs EB-3000xyz) with non-square pixels, yielded 233 paired eyes before and after stripping (Methods; Supplementary Section S1). In this cohort, the same density dependence holds and extends to corneas with lower endothelial cell density. It lies mostly below the density range of the first (1,690–2,716 against 2,072–3,303 cells/mm^2^), largely below the minimum donor densities used by eye banks, and it responds to stripping as the first cohort predicts for such tissue: order fell in 71% of eyes (median change −19%), together with cell density (−3.6%) and mean force (−23%), whereas all three rose slightly in the denser primary cohort.

After correcting for regression to the mean, a single density curve describes both cohorts (Supplementary Fig. S1e): the share of eyes gaining order rises from about 20% in the sparsest eyes to 80% in the densest (*R*^2^ = 0.78 across deciles), and the median change from about −20% to +15% (*R*^2^ = 0.62), with the second cohort lying slightly below the curve. Across both cohorts, collective order therefore responds to stripping along one continuum of baseline density: corneas with higher endothelial cell density gain order, and those with lower density lose it.

### *K*_ds_ and force separate media effects at matched density

Culture also allows media conditions to be varied directly. We analyzed primary rabbit corneal endothelial cells isolated from three independent paired corneas and expanded on tissue culture plastic under three proliferation media conditions: FGF2-containing medium from the established dual-media culture system of Peh *et al*. ^17^ as the reference condition, epidermal growth factor alone (EGF), and EGF supplemented with TGF-*β* receptor inhibitor SB431542 and p38 MAP kinase inhibitor SB203580 to inhibit endothelial-to-mesenchymal transition-associated signalling (EGF+Dual Inh.). Cells from each cornea were split across the three media, so each cornea contributes to every condition; 14–17 reliable images per condition were analyzed at 20× (45 images from three corneas), and representative images from two corneas, with per-image *K*_ds_, are shown in Fig. 7a. Conditions were compared with a mixed-effects model on all images, with cornea as a random effect (Methods).

**Figure 7.**
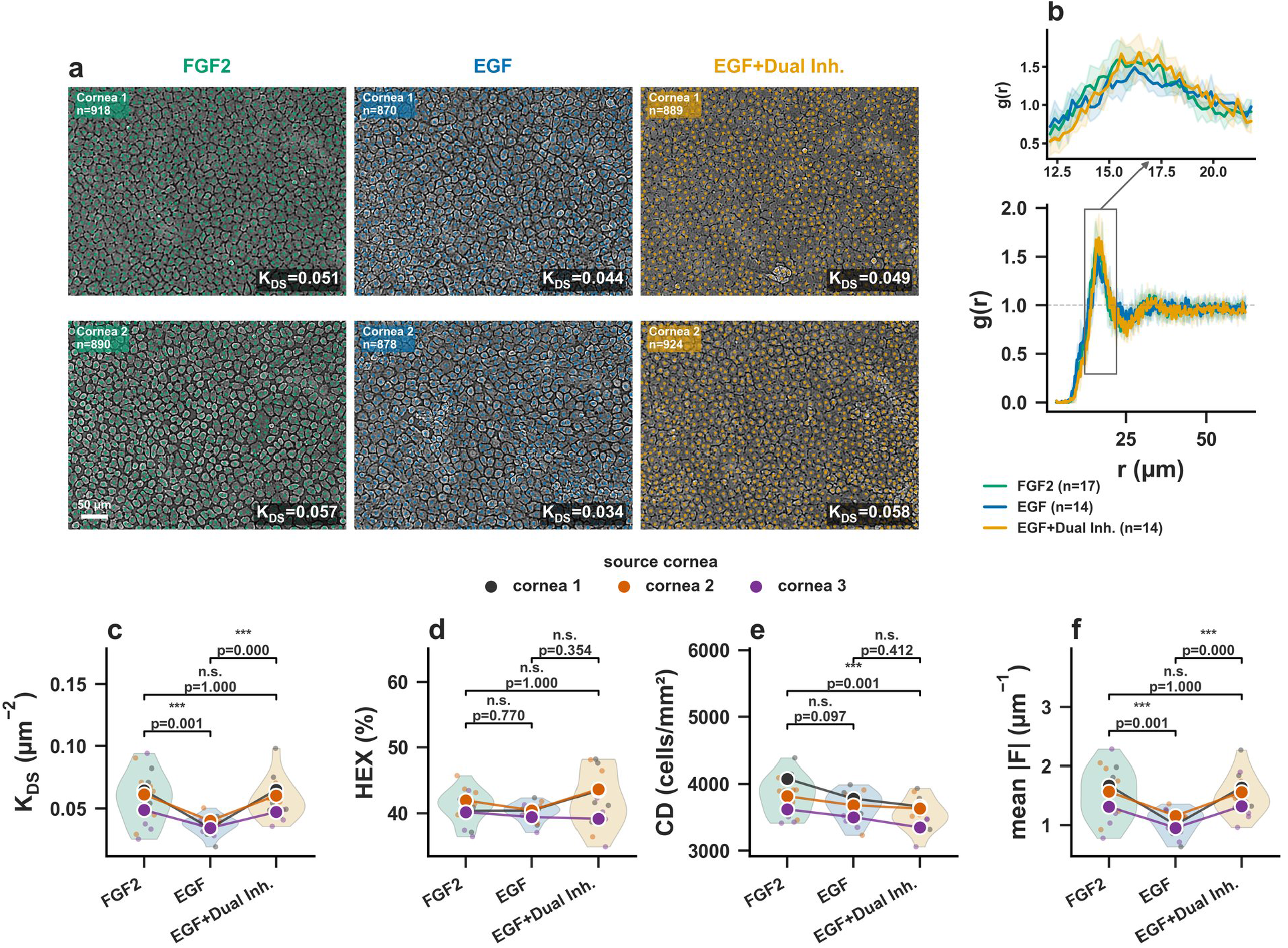
Collective order and intercellular force separate media conditions at matched density. **(a)** Representative segmented phase-contrast images of primary rabbit cells under each medium, from two of the three corneas, with per-image *K*_ds_ annotated. Scale as marked; contrast adjusted for print. **(b)** Population-averaged *g*(*r*) per medium with IQR bands; the view above the panel magnifies the first peak (12–22 *µ*m). **(c–f)** *K*_ds_, hexagonality, cell density and mean|*F* | per medium, shown as SuperPlots ^18^: small points are individual images coloured by source cornea, large points are the mean of each cornea, joined across media, and violins show the image-level distribution. Brackets give *P* values from a mixed-effects model on log-transformed values with cornea as a random effect (*n* = 3 corneas, 45 images), Bonferroni-corrected for the three comparisons.

All three conditions fall into a narrow density range of 3547–3837 cells/mm^2^, and density did not separate EGF from either other condition (Fig. 7e). Hexagonality did not differ between any of them (40.1–42.1%, all *p* ≥ 0.35; Fig. 7d). Yet collective order singled out EGF: *K*_ds_ was 37% lower than with FGF2 (*P* = 7 × 10^−4^) and 37% lower than with EGF+Dual Inh. (*P <* 10^−4^), lowest in each of the three corneas (Fig. 7c), while FGF2 and EGF+Dual Inh. were indistinguishable (*P* = 1.00). The population-level *g*(*r*) curves show nearly the same equilibrium spacing in all three (first peak at 15.6–16.5 *µ*m) but different first-peak heights (Fig. 7b): more neighbours sit at the preferred spacing, giving tighter spatial order at the same density.

Intercellular force follows the same ranking (Fig. 7f): mean|*F* | under EGF was 31% lower than with FGF2 (*P* = 6 × 10^−4^) and 31% lower than with EGF+Dual Inh. (*P <* 10^−4^), again in each of the three corneas, and the latter two were indistinguishable (*P* = 1.00). SB431542 blocks endothelial-to-mesenchymal transition and SB203580 suppresses culture-induced senescence, and both preserve N-cadherin and ZO-1 at cell junctions ^26–28^. These are the structures through which neighbouring cells transmit force to one another, so cells in conditions that keep them intact transmit more force locally and produce a better global arrangement.

## Discussion

This study shows that corneal endothelial tissue has a field-spanning collective order, quantified by the spring constant *K*_ds_, which is distinct from classical metrics at baseline and responds most to mechanical perturbation. Once the endothelium and Descemet’s membrane are released from the stromal support, how order changes depends on baseline density: after correction for regression to the mean, corneas with higher endothelial cell density mostly gain order and those with lower density lose it, along one density curve that spans two independent cohorts and crosses 50% near 2,400 cells/mm^2^, within the range of minimum donor densities used by eye banks. Effective intercellular forces computed from the same potential track these changes.

An independent clinical cohort from a different continent, eye bank and imaging device confirms and extends this picture. Its eyes lie mostly below the density range of the primary cohort, and there stripping reduced all three quantities: cell density, collective order and mean force. Once corrected for regression to the mean, these eyes fall on the same density curve as the primary cohort, with a modest downward shift that may reflect differences in instrument, storage and preparation. At the lower densities of this cohort, stripping therefore tends to degrade rather than improve the collective organization of the graft.

Extension to cultured cells reinforces the force–order link from a different angle. Media conditions that preserve N-cadherin and ZO-1 at cell junctions produce higher intercellular force and higher collective order than conditions allowing junctional degradation, while hexagonality does not differ and cell density does not follow the same pattern. This suggests a relationship between the junctional structures that transmit intercellular force and the field-spanning order they sustain. Serial passaging, in turn, drives collective order down in an exponential decay, and primary human cultures from a second laboratory reproduce the same decline.

The clinical need for a tissue-quality metric beyond cell density is well established. Filev *et al*. ^29^ assessed 1,031 organ-cultured corneas and found that endothelial cell density was a weak predictor of transplant suitability (AUC = 0.655) while morphological parameters added no predictive value (AUC = 0.597). Wakefield *et al*. ^30^ analysed 9,415 penetrating keratoplasties from the UK national registry and found no significant effect of preoperative donor ECD on 5-year graft survival above 2,200 cells/mm^2^. Wilhelm *et al*. ^31^ followed 2,956 keratoplasties for 10 years and found that grafts prepared with different surgical techniques lost cells at very different rates despite starting with similar cell densities. Only 3% of DMEK grafts retained ECD above 1,000 cells/mm^2^ at 10 years. Across these studies, the same conclusion emerges: cell density and morphology do not predict which grafts will survive and which will fail. Mehta *et al*. ^13^ identified this gap explicitly, noting that current quality control methods for cultured endothelial cells are destructive and that the field requires a non-invasive biomarker capturing a dimension of tissue quality inaccessible to per-cell counting. Collective order offers such a dimension: it is measured from the same specular images already used for cell counts, and its response to stripping changes across the range of density thresholds that eye banks already apply.

Yamamoto *et al*. ^10^ proposed exactly such a biomarker: a spring constant *K* derived from the radial distribution function of cell positions, which outperformed cell density in predicting graft trajectory in a small cohort (AUC 0.85 vs 0.56) by capturing field-spanning properties of the tissue. Despite this result, the concept has not yet seen widespread clinical adoption due to practical and conceptual limitations. It relied on semi-manual segmentation and expert-supervised curve fitting with no automated software, making it difficult to implement at scale.

Being fully automated, Cornealia removes the practical barrier, processing a specular microscopy image in under two seconds with no specialized hardware, scaling the analysis to over 2,000 images across two clinical cohorts, two culture systems, and two imaging devices. Having been applied to serial passaging, different culture conditions, and two independent clinical cohorts across continents, this study addresses what the original work and the accompanying commentary^13^ identified as the missing validation. On the conceptual side, the results ground the abstract spring constant in intercellular mechanics: *K*_ds_ changes go together with changes in local forces computable from the same *g*(*r*) curve, with the suggestion that the molecular junctions transmitting those forces determine the level of collective order a tissue can sustain.

Treating the corneal endothelium as a physical system is not new. Brookes^32^ showed that it behaves as a glassy viscous foam near the jamming boundary, with its phase state independent of cell density. After individual cell death, the monolayer transiently fluidizes, rearranges, and re-jams into structural equilibrium with minimum energy expenditure. Bi *et al*. ^33^ demonstrated theoretically that confluent tissues can undergo jamming transitions at constant density, controlled by cortical tension and cell-cell adhesion. These studies ask whether the tissue is solid or fluid. *K*_ds_ asks a different question: once cells have settled into their positions, how tightly does the monolayer hold them at their preferred spacing? The density-dependent reorganization we observe after stripping is consistent with Brookes’s observation that the endothelium actively rearranges after perturbation. What we add is that the direction of this rearrangement depends on where the tissue starts: corneas with higher endothelial cell density reorganize toward higher order, and those with lower density away from it.

The perturbation that revealed this dependence is itself a change in boundary conditions: stripping the Descemet’s membrane–endothelium complex from the stroma. In vertex models of confined epithelia, the strength of confinement alone shifts cell packing and disorder ^34^, so releasing the monolayer from the stromal support may be expected to change how it organizes. Lie *et al*. ^35^ showed that stripping leaves cell density essentially unchanged (2701 vs 2719 cells/mm^2^) while producing linear cellular disruptions that regenerate completely within weeks, implying an intrinsic capacity for spatial self-organization once the tissue is released from the stroma. Our data extend this observation to a much larger cohort and show that the reorganization is not uniform: its direction and magnitude depend on the tissue’s baseline density rather than following a single recovery trajectory.

The residual force analysis connects this density dependence to intercellular mechanics. In confluent tissues, intercellular forces are inherently nonlocal and nonadditive: the force on each cell emerges from the full configuration of its neighbors and cannot be decomposed into independent pairwise interactions^33^. The Delaunay truss model captures this by computing each node’s residual force as the vector sum of all neighbor contributions. The spatial mean of this unbalanced force tracks collective order across donors (*ρ* = 0.73), and its change tracks the change in order (*ρ* = 0.56), linking the interaction of local forces at global scale to a collective property of the tissue. Tissues that gained order mostly also gained force, and tissues that lost order lost force. Brookes ^32^ observed that endothelial cell migration follows the local orientation of maximum principal stress through a cooperative mechanism spanning many cell-to-cell junctions, termed plithotaxis. Our data are consistent with this picture, in which reorganization of the monolayer follows its field of intercellular forces.

To confirm that collective order is a field-spanning property, we asked whether relaxing only the most stressed nodes could account for the observed reorganization. It could not. When the three dominant stress concentrators in each image were released computationally, the change in *K*_ds_ agreed in direction with real stripping in 60% of donors, barely above the 54% expected by chance. The forces show the failure more clearly: after real stripping, mean force and order moved together in 69% of donors, whereas local release lowered the force (median −23%) while raising order, so the two moved in opposite directions in two-thirds of donors (Supplementary Section S4 and Supplementary Fig. S4). Collective order cannot be adjusted by patching local imbalances. It is a global property of the monolayer, and changing it requires the kind of whole-field rearrangement that stripping from the stroma triggers.

The culture experiments shed more light at the molecular level. *K*_ds_ distinguished EGF from EGF+Dual Inh. (37% lower under EGF, *P <* 10^−4^) at indistinguishable cell densities (*P* = 0.41),a difference in spatial organization that the conventional metrics do not see. The two inhibitors act on the structures through which neighboring cells transmit force: SB431542 prevents TGF-*β*-driven endothelial-to-mesenchymal transition, preserving ZO-1 and Na^+^/K^+^-ATPase at the plasma membrane^26^, while SB203580 counteracts p38 MAPK-mediated senescence and maintains N-cadherin, ZO-1 and Na^+^/K^+^-ATPase at cell junctions^27,28^. With these junctional proteins preserved, mean residual force and collective order were as high as under FGF2 (⟨|*F* |⟩ = 1.51 *µ*m^−1^, *K*_ds_ = 0.058), whereas EGF alone, which permits junctional degradation, produced the lowest values (⟨|*F* |⟩ =1.04 *µ*m^−1^, *K*_ds_ = 0.036). Conditions that keep adhesion proteins at the membrane thus produce stronger force networks and higher collective order, consistent with N-cadherin and ZO-1 carrying the forces that the Delaunay truss captures.

These observations suggest a three-level mechanistic chain: junctional adhesion proteins provide the physical coupling between adjacent cells, this coupling generates the intercellular forces measured by the Delaunay truss, and the collective balance of those forces determines *K*_ds_ across the monolayer. This chain is correlative, not causal: the cultured cell experiment was not designed to isolate force generation from other phenotypic effects of the inhibitors. A direct test would combine spatial force analysis with immunofluorescence for N-cadherin and ZO-1 on the same samples, mapping whether the stress concentrators identified by the residual force field correspond to sites of junctional disruption.

This study has several limitations. The design is single-arm, with one image before and one after stripping per donor; the correction for regression to the mean therefore relies on a constant measured from split images rather than on a control arm. The intercellular forces are effective forces inferred from cell positions, not measured directly. Segmentation of the second cohort was tuned by visual inspection, because no instrument-reported densities were available. The culture experiments rest on three corneas per comparison. Finally, this study includes no transplant outcome data; whether pre-transplant *K*_ds_ predicts graft outcome is the subject of ongoing work.

The findings presented here open several directions that this study was not designed to address. The most immediate clinical question is whether pre-transplant *K*_ds_ predicts post-transplant graft survival. *K*_ds_ is distinct from cell density and responds to perturbation in a density-dependent way, but whether it predicts graft outcome requires outcome data, which we are now collecting. Cornealia makes such a study feasible: it processes a specular microscopy image in under two seconds with no specialized hardware. Including donor age, medical history and surgical parameters would show whether the link between baseline density and the response to stripping varies across populations, and whether pre-transplant *K*_ds_ adds to cell density in predicting graft longevity.

On the mechanistic side, the three-level chain from junctional proteins to forces to collective order remains correlative. Direct molecular studies combining force field mapping with immunofluorescence for N-cadherin and ZO-1 on the same samples would test whether the spatial distribution of force magnitude across the monolayer corresponds to the spatial pattern of junctional protein localization at the membrane. Traction force microscopy would provide an independent measurement of cell-substrate forces to validate the Delaunay truss predictions against physical force data rather than spatial statistics alone. Live cell imaging during and after perturbation would capture how cells rearrange and forces redistribute over time, testing whether the density-dependent response seen here unfolds as the force field predicts.

## Methods

### Clinical datasets

#### Primary cohort

The original dataset comprised 1,607 donor cornea specular microscopy images acquired using a CellChek D specular microscope (Konan Medical, Hyogo, Japan), provided by the Singapore National Eye Centre. Each donor cornea was imaged before tissue preparation (Before Stripping, BS) and after the Descemet’s membrane–endothelium complex was stripped from the stroma for endothelial keratoplasty (After Stripping, AS). BS and AS images were acquired separately from the central endothelium as part of routine eye-bank evaluation; they sample the same central region but not the same field or cells. Twenty-two images with non-standard filenames (clinician-flagged as unusable due to patient unsuitability, procedural damage, or failed pre-strip assessment) were excluded, leaving 1,585 images (792 BS, 793 AS) for analysis. All 1,585 images were successfully processed (zero failures). Pair matching by full image name (minus the AS/BS suffix) yielded 784 BS/AS pairs. After reliability filtering (both images in a pair must pass the signal-processing quality gate for collective order analysis), 782 pairs from 779 donors (three donors contributed two pairs) remained for paired analysis. The pixel calibration was *p* = 0.58 *µ*m/px based on the instrument’s field of view (400 × 300 *µ*m at 684 × 514 px). Table 2 lists each sample size used in the paper and the step that defines it.

### Second cohort

An independent cohort of 570 specular microscopy images was obtained from a second eye bank (Lions VisionGift, Tampa, FL), captured on a HAI Labs EB-3000xyz with a field of view of 500 × 450 µm (width × height) on 640 × 480 px images. The pixels are therefore not square (0.781 × 0.938 µm), and all distances were computed with per-axis pixel sizes (Supplementary Section S8). Images were processed with the same pipeline and the same quality gate as the primary cohort. Because image quality was lower in this cohort, segmentation was tuned on these images by inspecting overlays of the detected cells: the automatic scale estimate was bypassed, the Laplacian-of-Gaussian range was set to 6–12 px with a detection threshold of 0.0005, and the overlap allowed between neighbouring detections was raised from 0.5 to 0.8, so that detections of adjacent cells were not discarded as overlapping. Of the 570 images, 530 (93%) passed the quality gate.

Each image carries a donor identifier, an eye designation and a processing phase. Images labelled *quarantine* were taken before stripping and those labelled *post cut* after, so a pair is defined by donor and eye across the two phases. When an eye had several images in the same phase, their values were averaged. Pairs missing *K*_ds_ or cell density in either state were dropped, and the same 1st–99th percentile trim on *K*_ds_ before and after was applied as in the primary cohort, leaving 233 paired eyes.

### Culture of primary corneal endothelial cells

#### Culture reagents

Human Endothelial-Serum Free Medium (SFM), Ham’s F12, Medium 199, Dulbecco’s phosphate-buffered saline (dPBS), TrypLE™ Select (TS), amphotericin B, penicillin and streptomycin were obtained from Life Technologies (Carlsbad, CA, USA). Collagen IV from human placenta, and trypan blue (0.4%) were from Sigma (St. Louis, MO, USA). Insulin/Transferrin/Selenium (ITS) was from Corning (Corning, NY, USA), and ascorbic acid was from Spectrum Chemical Mfg. Corp. (New Brunswick, NJ, USA). Human recombinant basic fibroblast growth factor (bFGF), and rho-associated, coiled-coil protein kinase (ROCK) inhibitor Y-27632 were from Miltenyi Biotec (Bergisch Gladbach, Germany). Liberase TH was purchased from Roche (Mannheim, Germany). Fetal bovine serum used to supplement the culture medium was purchased from Atlas Biologicals (Fort Collins, CO, USA). Finally, the staining solutions (Solution 18) used with the NucleoCounter NC-250 system for cell counting was obtained from Chemometec (Allerød, Denmark).

#### Research-grade human corneo-scleral tissues

A total of 9 donor cornea pairs deemed unsuitable for corneal transplantations were procured for this study from either the Lions Eye Institute for Transplant and Research (Tampa, FL, USA) or the Saving Sight (Kansas City, MO, USA), with informed consent obtained from the next of kin. All donor corneoscleral rims were preserved and transported to Singapore Eye Research Institute (SERI) in Optisol-GS (Bausch & Lomb, Rochester, NY, USA) at 4 ^°^C until they were processed.

#### Primary cell isolation and cell culture

Primary human CEnCs were isolated and propagated using a dual-media culture system as described^17^ and expanded to the second passage. Briefly, CEnCs were isolated by enzymatic digestion of the peeled corneal endothelium/Descemet’s membrane for approximately 4 hours. The dislodged clusters of CEnCs were treated with TS for 5 minutes to dissociate them into smaller clumps and single cells, then washed twice and seeded onto Collagen IV-coated vessels in corneal endothelial maintenance/stabilization medium (M5-Endo; Human Endothelial-SFM supplemented with 5% serum and 10 *µ*M ROCK-I) overnight. The CEnCs were subsequently cultured in proliferative medium (M4-F99; Ham’s F12/M199, 5% serum, 20 *µ*g/ml ascorbic acid, 1× ITS, 10 ng/ml bFGF, and 10 *µ*M ROCK-I) to promote proliferation. Once cultures reached 80% to 90% confluence, they were switched back to M5-Endo and maintained for at least one day before sub-culturing via single-cell dissociation using TS. Bright-field images of CEnCs for analysis were captured immediately after TS exposure to visualize their cell borders. Dissociated CEnCs were then plated at a density of at least 1 × 10^4^ cells/cm^2^ (split ratio of approximately 1:3 to 1:4, depending on final cell counts before passaging^36^) onto Collagen IV-coated surfaces, stabilized in M5-Endo, and expanded in M4-F99. All cultures were incubated at 37 ^°^C in a humidified atmosphere with 5% CO_2_.

#### Preprocessing

Cornealia accepts two input types. For CellChek D composite output (798 × 1078 px), the pipeline automatically extracts the cell region (684 × 514 px) and the metrics panel (193 × 156 px) using fixed coordinates determined from the instrument’s output layout. Green alignment dots overlaid by the instrument for focus confirmation are detected via HSV colour-space masking (hue 35–85, saturation *>* 40, value *>* 40) and removed using Telea inpainting^37^ with a 3-pixel radius. Device-reported metrics (CD, CV, HEX, SD, Ave) are extracted from the metrics panel using EasyOCR for validation purposes. For standalone specular microscopy images from any instrument, the image is passed directly to segmentation, bypassing all CellChek-specific steps. For this study, the cell regions and device metrics of the primary cohort were extracted once with this step, and the cropped images were then processed as standalone images, so that the slow OCR step was not repeated in every analysis run.

### Segmentation and classical metrics

#### CellADE segmentation

CellADE (Cell Adaptive Detection Engine) is a training-free, CPU-only segmentation algorithm for the low-contrast, uneven illumination typical of specular microscopy. Its guiding idea is that cells in a specular image form a quasi-periodic texture, so their size can be read from the image’s own power spectrum rather than learned from annotations.

After contrast-limited adaptive histogram equalization (clip limit 0.03), a central crop is mirror-tiled into a 4 × 4 mosaic and its radial power spectrum computed. The dominant frequency gives a characteristic cell scale 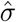, which sets the search range for multi-scale Laplacian-of-Gaussian blob detection,

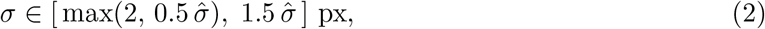

whose maxima are the cell seeds (a default range of 6–30 px is used when the spectral estimate is skipped). Each seed then anchors a local two-class fuzzy C-means clustering ^38^ of a circular patch scaled to its own *σ*; pixels with membership *u >* 0.4 in the brighter cluster are cell interior. Clustering locally rather than thresholding globally is what makes the method tolerant of illumination gradients. The union of these per-seed masks forms the foreground, on which a marker-controlled watershed^39^ seeded by the blob centres assigns each cell a label, and a second pass expands labels into the thin dark intercellular boundaries.

Three settings carry the adaptation. The LoG scale range encodes the expected cell size in pixels. When the spectral estimate fails, for example on images with atypical texture, a plausible range is found with a few trials on a handful of images and then fixed for the whole dataset. The LoG detection threshold sets how faint a blob may be and still become a seed; lowering it recovers cells in dim regions and in low-quality images. The FCM membership threshold plays the same role at the pixel level, setting how dim a pixel may be and still count as cell. None of these requires retraining. The settings used for each clinical cohort are given in Supplementary Section S3.2 and in the description of the second cohort above. Full details of the strip-based region selection, the fallback range for outlying 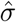, guttae protection during gap filling, pseudocode, and the complete parameter table are in Supplementary Section S7.

### Morphological metrics

From the segmented label image, five classical parameters are computed using only interior cells—those whose centroids lie at least MAX R = ⌊0.15 × min(*H, W* )⌋ pixels from any image border—to avoid bias from truncated boundary cells. Let *N*_int_ denote the number of interior cells, *a*_*i*_ the area of interior cell *i* in pixels, and *p* the pixel calibration (*p* = 0.58 *µ*m/px for the primary cohort). For non-square pixels, *p*^2^ is replaced by the pixel area *s*_*x*_*s*_*y*_, the margin is set from the physical size of the image, and hexagonality uses distances weighted per axis (Supplementary Section S8).

Cell density is computed as the number of interior cells divided by the interior image area:

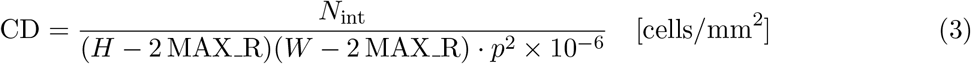

Average cell area and standard deviation:

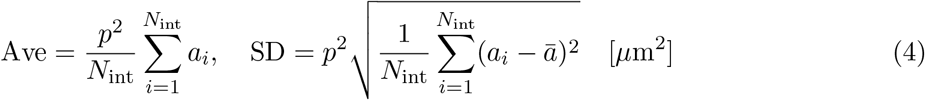

Coefficient of variation:

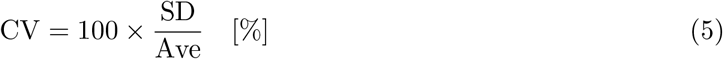

Hexagonality is determined via Delaunay triangulation of interior cell centroids. For each interior cell *i*, the neighbour set *N*_*i*_ consists of all interior cells sharing a Delaunay edge:

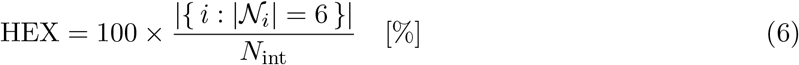

### Collective order analysis

Collective order quantifies the spatial organization of the endothelial monolayer through the curvature of an effective interaction potential derived from cell positions. In Cornealia, the analysis proceeds in four stages: (i) the radial distribution function *g*(*r*) is computed from the cell centroids returned by segmentation; (ii) a physics-informed step response model is fitted to the entire *g*(*r*) curve rather than to a narrow window around its peak, so that the whole curve constrains the fit against the noise clinical data carry; (iii) the fit is screened by four reliability checks, and the smooth fitted curve is converted to the effective interaction potential *V* (*r*) = − ln[*g*(*r*)], whose curvature at the first minimum gives the spring constant *K*_ds_ = *V* ^′′^(*r*_0_), the collective order parameter;(iv) four alternative estimators are computed alongside it for comparison, pairing Yamamoto’soriginal quadratic with the asymmetric Morse potential, each fitted both on the raw region located by argmin and on the boundaries the step response fit provides.

### Radial distribution function

Cell centroids are extracted as centre-of-mass coordinates from the segmented label image. To avoid boundary artifacts, only interior cells are used as reference cells, defined as those located at least MAX_ R = ⌊0.15 ×min(*H, W* )⌋ pixels from any image border. The radial distribution function is:

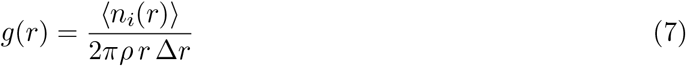

where *n*_*i*_(*r*) is the number of cell centroids in a shell of thickness Δ*r* = 1 px at distance *r* from interior cell *i*, evaluated at shell centres over 0 ≤ *r <* MAX R so that every shell lies within the image, *ρ* = *N*_cells_*/*(*H* × *W* ) is the global cell density in px^−2^, and ⟨·⟩ denotes averaging over all interior cells. For non-square pixels, distances are weighted per axis and expressed in units of 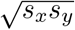(Supplementary Section S8). Each radius therefore yields a single value: the count of neighbours at that distance, averaged over all interior reference cells and expressed relative to the count random spacing would produce. Savitzky–Golay smoothing (window 11, order 3) is applied for display purposes only; all fitting operates on raw *g*(*r*).

### Effective interaction potential

The effective interaction potential is derived via the inverse Boltzmann relation from statistical mechanics. Setting *k*_*B*_*T* = 1 (dimensionless units):

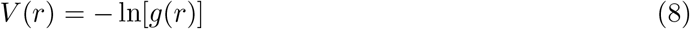

where *g*(*r*) values ≤ 0.05 are excluded to avoid numerical instability. While the corneal endothelium is not a thermodynamic system in the classical sense, Yamamoto *et al*. ^10^ showed that this formalism yields a meaningful effective potential whose curvature quantifies the sharpness of spatial confinement. The spring constant *K* = *V* ^′′^(*r*_0_) at the first minimum, which coincides with the first peak of *g*(*r*), measures how precisely cells maintain the preferred spacing *r*_0_: higher *K* indicates stiffer confinement.

### Step response quality gate

The signal *g*(*r*) − 1 is expected to exhibit damped oscillatory behaviour for well-ordered endothelial monolayers. Rather than trimming the exclusion zone and fitting the oscillatory portion alone, the entire *g*(*r*) curve is fitted with an underdamped step response model (Eq. 1):

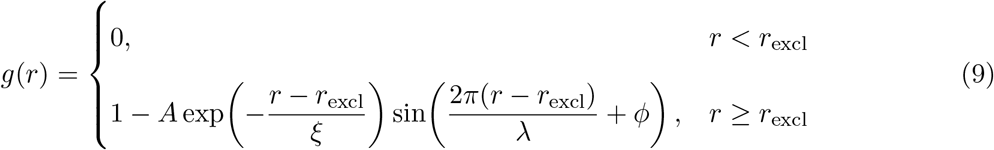

where the transition at *r*_excl_ is smoothed by a logistic function of fixed width 0.3 px for differentiability, *A* is the oscillation amplitude, *ξ* is the correlation length, *λ* is the wavelength, *r*_excl_ is the exclusion zone boundary and *ϕ* is the phase offset. Parameters are estimated via bounded nonlinear least squares (`scipy.optimize.curve_fit`, trust region reflective) with bounds:

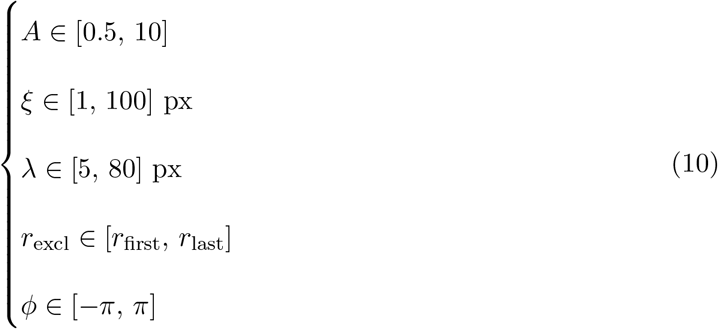

Reliability is assessed by a composite criterion combining four checks:

#### Form check

The fitted oscillatory component (evaluated for *r > r*_excl_) must exhibit a positive peak with amplitude *>* 0.02 in the first half of the fitted range, followed by a zero crossing, followed by a negative trough with amplitude *<* −0.01.

#### Parameter sanity

The fitted parameters must fall within permissive bounds that exclude degenerate fits: *ξ <* 95 px (55 *µ*m) and 8 *< λ <* 70 px (4.6–41 *µ*m). These are wide relative to the observed values, as the median fitted wavelength is 28.8 px (16.7 *µ*m), close to the median first-peak spacing of 17.9 *µ*m, so the gates reject only very short ripples or wavelengths longer than the field of view can support. The thresholds are defined at the reference pixel size (*p*_ref_ = 0.58 *µ*m/px); for images acquired at a different calibration *p*, the upper bounds are multiplied by *p*_ref_ */p* and the lower bound divided by it, so the gates track physical length across instruments.

#### Peak height

The raw *g*(*r*) must show a meaningful oscillation above unity: its maximum over the first half of the radial range must exceed 1.10. *Goodness of fit and noise diagnostics*. *R*^2^ is computed on the oscillatory region only (*r>r*_excl_) to prevent the step rise from inflating goodness of fit. The fit must satisfy either 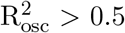(strong fit), or 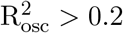 together with residuals that are both uncorrelated (lag-1 autocorrelation |ACF_1_| *<* 0.4) and small relative to the oscillation (SNR = *A/ σ*_residual_ *>* 1.2).An image is classified as reliable if and only if all four checks pass:

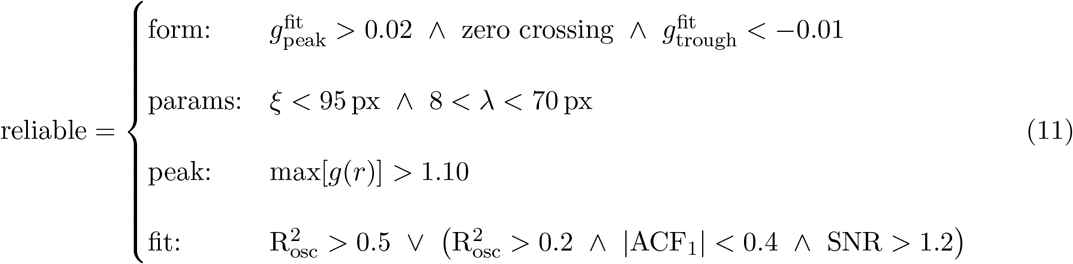

Of the 1,585 clinical images, 1,583 (99.9%) passed the quality gate.

### Spring constant extraction

Five estimates of *K* are computed per image: two parametric models fitted to raw *V* (*r*) under two region-selection strategies, plus *K*_ds_ from the step response fit.

#### Quadratic fit

Following Yamamoto *et al*. ^10^, raw *V* (*r*) is approximated as a degree-2 polynomial near its minimum, yielding *K* = 2|*a*_2_| where *a*_2_ is the leading coefficient.

#### Morse potential fit

The Morse potential admits asymmetry about the minimum, which the parabola cannot. It is fitted to raw *V* (*r*) via nonlinear least squares:

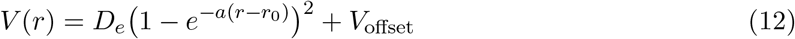

with initial guesses taken from the data (*D*_*e*_ from the observed well depth, *r*_0_ from the minimum location, *a* = 0.1) and positivity bounds on *D*_*e*_, *a* and *r*_0_, yielding 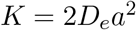.

Both models are fitted under two region strategies:

- *Raw region (Yamamoto):* ±10 px symmetric window around the argmin of raw *V* (*r*).
- *Guided region:* boundaries determined analytically from the step response fit. The left boundary is the zero crossing of *V*_fit_(*r*) before its minimum minus 3 data points; the right boundary is the first hump after the zero crossing beyond the minimum plus 3 data points. The margin points enforce curvature at the edges and prevent plateau fits.

#### *K*_ds_ *(recommended)*

The step response fit already provides a smooth analytical approximation to *g*(*r*), constrained by the entire curve rather than by the well alone. *K*_ds_ is taken from that same fit, so one fit yields both the reliability assessment and the spring constant. The fitted curve is evaluated on a dense grid of 1,000 points spanning the oscillatory region (*r > r*_excl_), converted via *V*_fit_(*r*) = − ln[*g*_fit_(*r*)], and its curvature at the first minimum gives 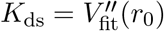.

All RMSE values are evaluated on the guided well region for fair comparison, and all *K* values are converted to physical units via *K*_*µ*m_ = *K*_px_*/p*^2^, with *p*^2^ = *s*_*x*_*s*_*y*_ for non-square pixels.

### Synthetic ground-truth validation

To benchmark the quality gate and the five extraction methods against a known answer, we generated synthetic *g*(*r*) curves from the same underdamped step-response model used for fitting (Eq. 1). For each of 1,000 random parameter combinations, with *A, ξ, λ, r*_excl_ and *ϕ* drawn uniformly from *A* ∈ [0.5, 1.5], *ξ* ∈ [5, 35] px (2.9–20 *µ*m), *λ* ∈ [15, 45] px (8.7–26 *µ*m), *r*_excl_ ∈ [10, 22] px (5.8–13 *µ*m) and *ϕ* ∈ [−*π/*2, *π/*2], the ground-truth spring constant *K*_true_ was obtained by double numerical differentiation of the noiseless potential on a 5,000-point grid. These intervals sit well inside the optimizer bounds, so fitting is never constrained at the edges of the generated space.

Each parameter set was rendered at 10 signal-to-noise ratios spanning SNR 1.5–50, yielding 10,000 synthetic curves in total. SNR is defined as the peak-to-trough amplitude of the noiseless oscillation divided by the noise standard deviation. Noise comprised an AR(1) component with coefficient 0.35 carrying 60% of the variance, plus additive white noise carrying the remainder. To test the gate against curves that should be rejected, an additional 2,000 non-oscillatory curves (flat, single bump, step rise and random walk; 50 of each type, 200 per SNR level, with noise standard deviation 0.3*/*SNR) were generated as negatives. Sensitivity is the fraction of genuine oscillatory curves the gate admits (one minus the false negative rate); specificity is the fraction of non-oscillatory curves it rejects (one minus the false positive rate).

## Statistical methods

### Statistical analysis

#### Clinical cohort

BS and AS values were compared within donors, and no parametric assumption was made. Per-donor percentage change is Δ% = 100 × (AS − BS)*/*BS. For the comparison across metrics (Fig. 5f) we use the symmetric percentage change, Δ%_sym_ = 100 ×(AS−BS)*/*[(AS + BS)*/*2], which counts rises and falls of equal size equally; its mean is reported with a 95% confidence interval (mean ±1.96 s.e.m.), and *P* values are from Wilcoxon signed-rank tests. In the binned analyses (Fig. 5h,i), the share of donors whose value rose is given with Wilson 95% confidence intervals and the median change with bootstrap 95% confidence intervals; trends are Spearman’s *ρ* across donors, with the *R*^2^ of the fitted curves across deciles. Correlations between metrics and between their paired changes are Spearman’s *ρ*, except the baseline correlations of *K*_ds_ with cell density and hexagonality (Fig. 5a,b), which are Pearson’s *r*. The share of donors whose *K*_ds_ rose after stripping is tested against 50% with an exact binomial test.

#### Trimming

No outliers were removed from the reliable pairs, except in the analysis-specific subsets listed in Table 2: a 1–99% trim on *K*_ds_ (BS and AS) for the binned and density-band analyses (*n* = 750); a 2–98% clip on both axes for the paired scatter plots (Fig. 5c, *n* = 718; Fig. 5g, *n* = 722); a 1–99% clip on both axes for the change scatter plots of Fig. 6e,f; and removal of the top 1% of baseline *K*_ds_ for Fig. 6g (*n* = 774). In Fig. 3f the per-method *K* distributions are shown after excluding values outside *Q*_1_ − 1.5 IQR to *Q*_3_ + 1.5 IQR; this affects only the display and the coefficients of variation. In Fig. 3g no values are excluded; for *K*_ds_, points of the well that fall inside the model’s hard core (fitted *g <* 0.05) are excluded from its RMSE, where − ln *g* is undefined.

#### Cultured cells

For the cultured cells, images are positions within a culture and are not independent, so the biological replicate is the culture from each source cornea (three culture lines for the passage series, three corneas for the media conditions). Culture data are shown as SuperPlots^18^. For media conditions, pairs of conditions were compared with a linear mixed-effects model on log-transformed values with cornea as a random intercept, fitted to all images, with Bonferroni correction for three comparisons. For the passage series, adjacent passages were compared with paired *t*-tests on the line means (*n* = 3), and the reproducibility of the decay was assessed from the exponential decay rate fitted to each line separately.

A significance threshold of *P <* 0.05 is used throughout.

### Regression to the mean and the correction applied

Every donor in the paired dataset receives the same intervention and acts as its own control, so the design is a single-arm pre–post study. Regressing a change score on a baseline value in such a design is biased by two distinct artifacts^19,20^. Mathematical coupling arises because Δ*Y* = *Y*_2_ − *Y*_1_ contains *Y*_1_ by construction, inducing a negative association between them even when the intervention does nothing. Regression to the mean, on the other hand, stems from inherent measurement noise: donors with extreme low initial values tend to rise closer to the mean on remeasurement, while those with extreme high values tend to fall, producing a bilateral convergence toward the mean from both sides. Both artifacts push in the same direction and together produce the appearance of a restoring tendency pulling extreme tissues toward a common value.

We address this in three ways, applied at different points in the analysis.

#### Cross-variable binning

Measuring changes in a variable while binning or selecting on the same variable leads to regression to the mean^19^. To avoid this, binned displays select bins on one baseline variable and measure the change in a different one: baseline cell density against Δ*K*_ds_. Indirect leakage remains possible in proportion to the baseline correlation between the two variables ^40^ (Supplementary Table S5). Cell density and *K*_ds_ share 29% of their baseline variance, so each donor’s change is first corrected for regression to the mean (see below); uncorrected, the leak pushes the cross-variable trend downward (Supplementary Section S5.3). *K*_ds_ and *r*_0_ share 3%. Cell density and *r*_0_ are strongly coupled (*R*^2^ = 0.37), so they were not used for cross-analysis.

#### Correction for regression to the mean

We correct each donor’s change with the method of Varadhan *et al*. ^20^ for single-arm pre–post studies. The method compares the stripped donors with an imagined control arm, the same donors left unstripped, and summarizes that arm in a single constant *k*, which compares how tightly baseline and follow-up would track each other without the intervention with how tightly they track with it. Without stripping, baseline and follow-up are two measurements of the same tissue, so *k* can be measured from the agreement between the left and right halves of each image (Supplementary Section S5.4).

The correction is built by writing the same regression for both an imagined control arm and the treatment arm:

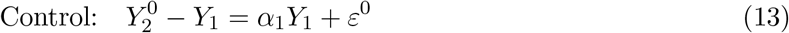

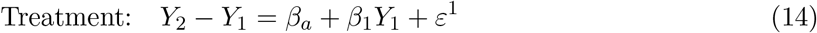

The naive estimate from the single-arm study is *β*_1_, but it contains the artifact *α*_1_ on top of the real effect 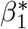, so 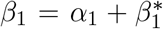. To isolate the real effect we need *α*_1_. Assuming the control-arm error is uncorrelated with baseline (*E*[*ε*^0^|*Y*_1_] = 0), solving for *α*_1_ gives:

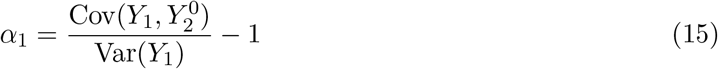

The −1 is the mathematical coupling term, present by construction. The covariance ratio is the regression to the mean component. Since Covc 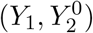 belongs to the unobserved control arm, we express it as a multiple of the observable covariance through the parameter *k*:

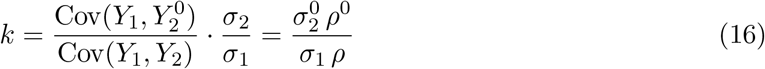

which gives:

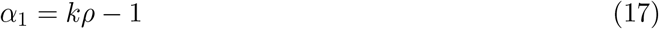

where *ρ* is the paired correlation between *Y*_BS_ and *Y*_AS_. When covariates *X* are present, *Y*_1_ is correlated with *X*, so the bias on the baseline coefficient and the covariate coefficients become entangled and must be solved together:

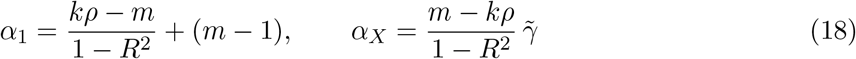

where *R*^2^ and 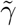come from regressing *Y*_BS_ on *X*_BS_, and *m* (set to 1) assumes that without intervention, the relationship between covariates and outcome does not change. The corrected coefficients are 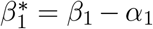 and 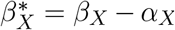, where *β*_1_ and *β*_*X*_ are the naive regression coefficients. Both *ρ* and *R*^2^ use the Olkin–Pratt unbiased estimators^20^. The full derivation is given in Supplementary Section S5.

For each image, the Spearman–Brown-corrected agreement between its left and right halves gives the reliability *R* of the full-image value (*K*_ds_: *R* = 0.46; cell density: *R* = 0.87; Supplementary Section S5.4), and *k* = *R/ρ*. For *K*_ds_ this gives *k* ≈ 1.0, the classical correction^21,22^. We therefore correct each donor’s change in *K*_ds_ at *k* = 1, 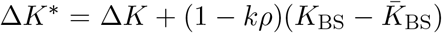, before binning on baseline cell density (Fig. 5h). The procedure is a Python reimplementation of the R package `resilience`.

### Shared segmentation error

*K*_ds_ and cell density are computed from the same centroids, so a segmentation error in one image changes both, and this shared error could by itself correlate their paired changes. We tested this with simulation–extrapolation (SIMEX), which applies to variables calculated from a common set of measurements with coupled error^41^. For each of the 779 donors with a reliable BS/AS pair (one pair per donor), additional segmentation error was added to the BS and AS centroids independently, either as missed cells (random removal) or as false detections (random addition), at levels *λ* = 0.025–0.20 of the cell count in steps of 0.025, with 100 replicates per level. At each level, *K*_ds_ was recomputed with the same pipeline, cell density as the interior cell count per interior area, and the correlation between Δ*K*_ds_% and ΔCD% exactly as in Fig. 5c. Perturbed images were flagged by the quality gate, and results were computed both with and without excluding flagged pairs. The correlation was extrapolated on the Fisher-*z* scale (linear and quadratic) back to assumed existing error rates of 2, 5 and 10%, since the true error rate is unknown; 95% confidence intervals were obtained by bootstrapping over pairs (1,000 resamples).

### Comparison of the two cohorts

Each eye’s change in *K*_ds_ was corrected for regression to the mean with the constant *k* measured within its own cohort (primary *k* = 1.0, second *k* = 2.3; Supplementary Section S1). Both cohorts were then pooled, and one logistic regression (direction of change) and one linear regression (median percentage change) on baseline cell density were fitted to all eyes. Whether the two cohorts need separate offsets was tested by adding a cohort indicator: a likelihood-ratio test for the logistic fit and an *F* -test for the linear fit. *R*^2^ is reported for each fit against the decile medians of both cohorts.

## Mechanical analysis

### Delaunay truss model and residual force computation

Interior cell centroids, those at least 15% of the image’s shorter dimension from any edge, were connected via Delaunay triangulation, a purely geometric construction that links each cell to those immediately surrounding it and so defines the tissue’s neighbour network without any free parameters; boundary cells subject to truncation artefacts are excluded. For non-square pixels, node positions use the pixel size along each axis (Supplementary Section S8).

Following the statistical mechanics framework of Yamamoto *et al*. ^10^, the effective interaction potential between cells is related to the radial distribution function by the potential of mean force:

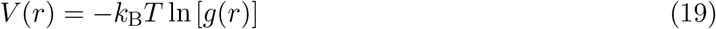

where *g*(*r*) is the damped oscillatory radial distribution function with parameters (*A, ξ, λ, r*_excl_, *ϕ*) obtained from the Cornealia pipeline. Setting *k*_B_*T* = 1, the pairwise force on each edge follows by differentiating the potential:

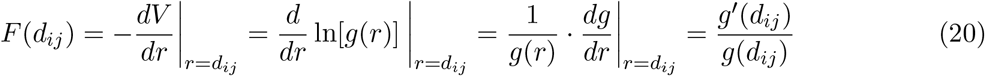

where *d*_*ij*_ is the measured distance between nodes *i* and *j*, and *g*(*r*) and *g*^′^(*r*) are evaluated analytically from the fitted step response parameters converted to physical units (*µ*m). Each edge carries a force determined by its length relative to the potential landscape: edges shorter than the equilibrium spacing *r*_0_ (the first maximum of *g*(*r*)) are under compression and thereby repulsive, while edges longer than *r*_0_ are under tension and attractive; its magnitude scales with the local gradient of *g*(*r*). Edges where *g*(*d*_*ij*_) *<* .05 are assigned zero force to avoid numerical instability in low-density regions.

The residual force on each node is the vector sum of all pairwise contributions from its Delaunay neighbours:

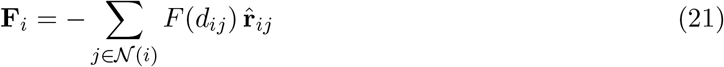

where *N*(*i*) denotes the set of Delaunay neighbours of node *i* and 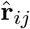is the unit vector from node *i* to node *j*. The leading minus sign carries the convention of Eq. 20: a compressed edge has *F >* 0 and pushes node *i* away from its neighbour, along 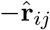, while a stretched edge has *F <* 0 and pulls it toward the neighbour. A node in perfect force balance has |**F**_*i*_| = 0; non-zero residual forces indicate local mechanical disequilibrium. The tissue-level summary statistic is the mean residualforce magnitude:

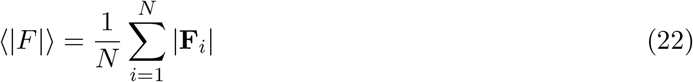

where *N* is the number of interior nodes. Summing magnitudes rather than vectors means opposing forces do not cancel, so ⟨|*F* |⟩ measures how far the monolayer sits from local mechanical equilibrium.

### Local release computation

Stripping triggers a field-spanning reorganization of the cells. Here we test whether local release at the highest-stress regions alone can account for it: if it can, letting the most strained cells relax along their force vectors should reproduce the measured *K*_ds_ after stripping. The test was run on every baseline image of the paired donors.

Three release sites were chosen from the top 10% of nodes by |**F**_*i*_|, taking the highest first and requiring each subsequent site to lie at least 4*r*_0_ from those already chosen. Each node *i* was assigned a mobility

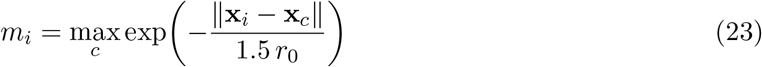

the maximum taken over the three release centres *c*, so that nodes near a site are free to move and those beyond a few cell spacings are effectively fixed. Every node was then displaced along its own residual force direction:

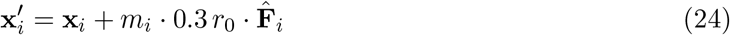

a step of at most a third of the equilibrium spacing. This is a single geometric step, not an energy minimization, so it captures the immediate local effect of release rather than the global equilibrium the tissue would eventually settle into. *K*_ds_ and mean|*F* | were then recomputed from the displaced centroids through the standard Cornealia pipeline and compared against the donor’s measured post-stripping values (Supplementary Fig. S4).

## Implementation and availability

### Implementation

Cornealia is implemented in Python and runs entirely on CPU, with no GPU, cloud, or deep learning dependency. Given a specular microscopy image and a pixel calibration, it executes the full pipeline end-to-end: segmentation (CellADE), classical morphometry (CD, CV, HEX), the radial distribution function *g*(*r*), the quality gate, collective order extraction (*K*_ds_), and the residual force field, using only standard scientific Python libraries (NumPy, SciPy, OpenCV, scikit-image). Batch processing parallelizes across CPU cores, achieving ∼1.8 s/image on a standard workstation with 10 workers and completing the 1,607-image clinical dataset in under 30 minutes.

### Software availability

Cornealia can be used in two ways. For programmatic access, it is a standalone Python `library: from cornealia import cornealia` provides the full API for single-image analysis and parallel batch processing, including command-line entry points for scripted workflows. For users without programming experience, Cornealia is integrated into CelLAB, an open-source desktop platform providing a graphical interface with structured workflows for image loading, parameter configuration, batch execution and interactive result visualization. CelLAB also includes CellIntella, a locally run assistant based on Llama 3.2 3B that answers questions about the platform using retrieval-augmented generation over its documentation. Pre-built packages for Windows, Linux and macOS, requiring no installation or GPU, are archived on Zenodo (https://doi.org/10.5281/zenodo.22018616), and source code is available at https://github.com/HanLab-BME-MTU/CelLAB. Cornealia and CelLAB are open-source software released under the GNU Affero General Public License v3.0 (AGPL-3.0). Commercial licenses that permit use without the AGPL obligations are available; inquiries should be directed to the corresponding author. The Llama 3.2 model used by CellIntella is not part of the CelLAB distribution under AGPL-3.0 and remains subject to the Llama 3.2 Community License.

## Acknowledgements

The authors thank the Singapore Eye Research Institute for providing the clinical dataset. We thank Lions VisionGift for providing the second clinical cohort, and the laboratories at the University of Texas Southwestern Medical Center and the Singapore Eye Research Institute for providing the cultured cell images. This work was supported by NIH GMO250807 PO0000003913 and the Michigan Technological University Health Research Institute for research support and student funding.

## Author contributions

Majid Rostami conceived and developed the pipeline, performed the analysis and wrote the manuscript. Mason Steele and Samuel Haarman contributed to software development. Addie Richard contributed to data curation. Darren Sai Kyauk contributed to clinical data acquisition. Saad Asim performed the cultured cell experiments. David Moore and Lisa Moul provided the independent cross-validation dataset. Gary Peh and Muhammad Rizwan contributed to concept development, discussion and manuscript revision. Jodhbir S. Mehta provided the clinical dataset, contributed to concept development, discussion and manuscript revision. Sangyoon J. Han supervised the project, provided funding and reviewed the manuscript.

## Competing interests

The authors declare no competing interests.

## Data availability

The primary clinical dataset is available from the Singapore Eye Research Institute upon reasonable request. The second clinical cohort was provided by Lions VisionGift (Tampa, FL, USA) and is available from the eye bank upon reasonable request. The cultured rabbit corneal endothelial cell images were generated at the University of Texas Southwestern Medical Center, and the cultured human corneal endothelial cell images at the Singapore Eye Research Institute; both are available from the respective laboratories upon reasonable request. The benchmark dataset of Daniel *et al*. ^14^ is available as described in the original publication.

## Code availability

Cornealia is part of the CelLAB platform and is available at https://github.com/HanLab-BME-MTU/CelLAB and on Zenodo (https://doi.org/10.5281/zenodo.22018616) under the GNU Affero General Public License v3.0 (AGPL-3.0). Commercial licenses are available; inquiries should be directed to the corresponding author.

## Supplementary Information

### S1. A second clinical cohort at lower cell density

To test whether the density dependence of the response to stripping holds in an independent setting, we analysed a second cohort from a different eye bank, Lions VisionGift (Tampa, FL), whose images were acquired with a HAI Labs EB-3000xyz specular microscope with non-square pixels (Methods; Supplementary Section S8). Of 570 images, 530 (93%) passed the quality gate. After pairing images before (*quarantine*) and after stripping (*post cut* ) by donor and eye, and trimming the 1st and 99th percentiles of *K*_ds_, 233 eyes remained. Their baseline cell density (median about 2,140 cells/mm^2^, range 1,690–2,716) lies mostly below that of the primary cohort (2,072– 3,303 cells/mm^2^). CellADE segmented these images well after a quick recalibration of its scale and detection ranges for this dataset, including images of very low quality in which the endothelial mosaic is faint or partly obscured (Supplementary Fig. S1a–c).

In this sparser cohort, stripping reduced all three quantities: cell density by 3.6% (*P* = 4 × 10^−6^), collective order by 19% (*P* = 6 × 10^−15^) and mean force by 23% (*P* = 6 × 10^−19^; Wilcoxon signed-rank tests on symmetric percentage changes), whereas in the primary cohort all three rose slightly. Order rose in only 29% of eyes (68 of 233; Supplementary Fig. S1d).

To compare the two cohorts on one density axis, we corrected each eye’s change for regression to the mean, using a constant *k* measured within each cohort. Each image was split into left and right halves (320 × 480 px, 250 × 450 *µ*m), and *K*_ds_ was recomputed on each half. The two halves are simultaneous measurements of the same tissue, so their agreement across images reflects measurement noise only. Their correlation (*r* = 0.44) gives the reliability of the full-image value by the Spearman–Brown formula, *R* = 2*r/*(1 + *r*) = 0.61. Dividing it by the correlation between baseline and follow-up order (*ρ* = 0.27, Olkin–Pratt corrected) gives *k* = *R/ρ* = 2.3. The same procedure gives *k* = 1.0 in the primary cohort (Supplementary Section S5.4).

The density dependence of the effect of stripping on collective order extends to this cohort: a single density curve describes both cohorts (Supplementary Fig. S1e), with the share of eyes gaining order rising with baseline density (logistic slope *P* = 4 × 10^−24^; *R*^2^ = 0.78 across deciles), as does the median change (*P* = 2.5 × 10^−37^; *R*^2^ = 0.62). The second cohort extends this curve to densities the primary cohort does not reach, where most eyes lose order, with a modest downward shift (an offset of −0.9 on the log-odds scale, *P* = 2 × 10^−4^; −16% in median change, *P* = 4 × 10^−15^) that may reflect differences in instrument, storage and preparation. Within the second cohort’s narrower density range alone, no trend is detectable (*ρ* = −0.01). Together, these results show that both the method and the clinical finding generalize to an independent cohort, instrument and density range.

**Figure S1.**
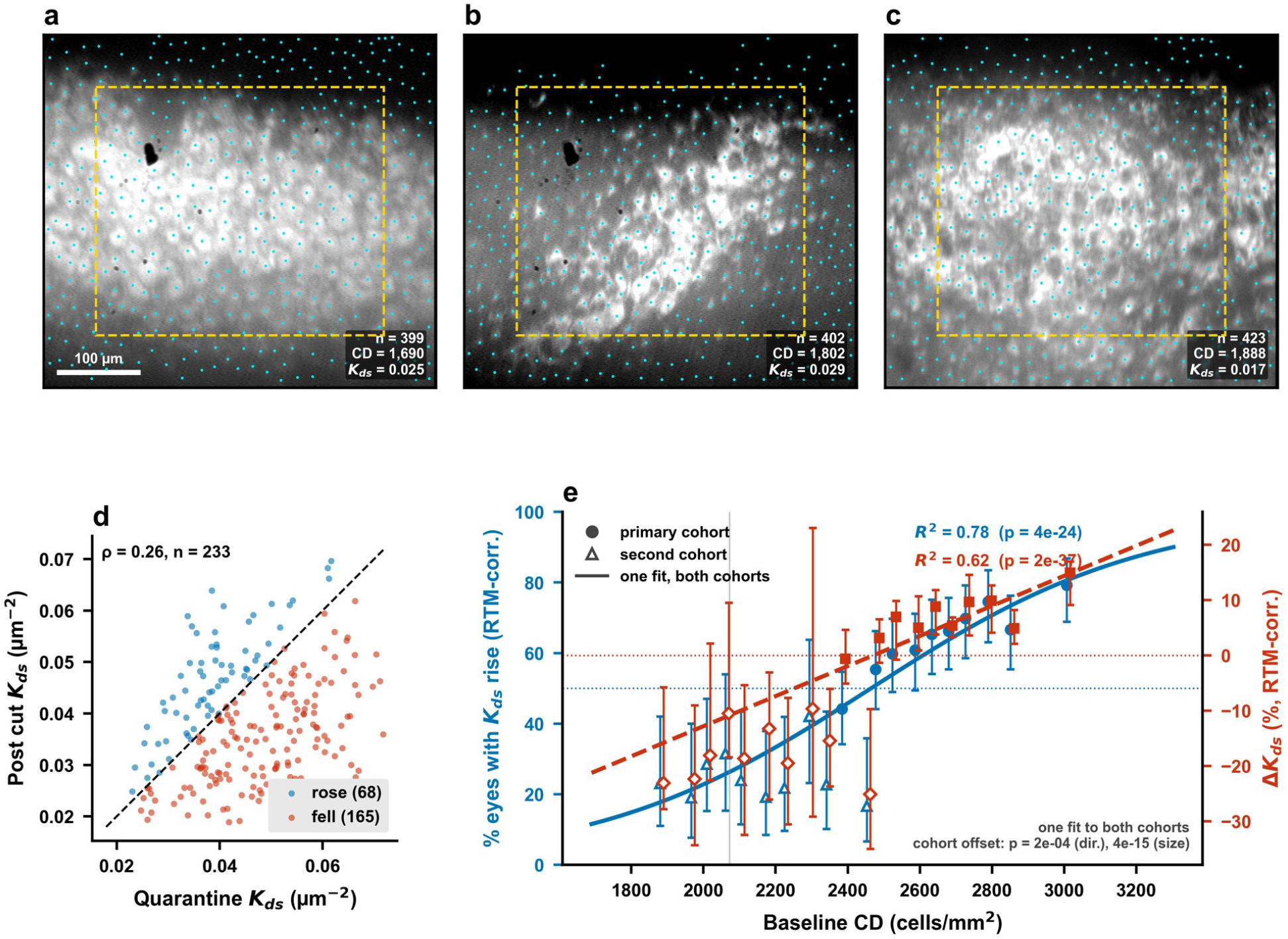
Second clinical cohort. **(a–c)** Three of the lowest-quality images (lowest signal-to-noise ratio of the *g*(*r*) fit), with the detected cells (cyan) and the interior region used for analysis (dashed yellow); images are shown in micrometres, correcting for the non-square pixels.**(d)** Paired *K*_ds_ before (*quarantine*) and after stripping (*post cut* ); the dashed line marks no change. **(e)** Change in *K*_ds_ per decile of baseline cell density in both cohorts (primary: filled markers; second: open markers), each corrected for regression to the mean with its own measured *k*. Left axis (blue): percentage of eyes whose *K*_ds_ rose, with Wilson 95% confidence intervals. Right axis (red): median percentage change, with bootstrap 95% confidence intervals. Lines: one fit to both cohorts (logistic and linear); the grey line marks the lowest density in the primary cohort.

### S2. Representative fitting examples across reliability levels

Four images spanning the signal quality spectrum illustrate how the quality gate and each spring constant extraction method respond to varying data quality (Supplementary Fig. S2). Row 1 shows a strong oscillatory signal 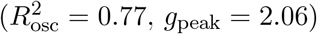 where all four checks pass and the five methods converge. Row 2 passes on every check as well 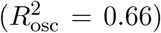 with a longer correlation length and a broader well; the five estimates still agree. Row 3 is rejected on the form check alone: the fitted oscillation is heavily damped (*ξ* = 2.7 px) and decays before producing the required trough, even though the fit quality 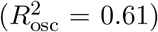 and peak height (*g*_peak_ = 1.33) would otherwise be acceptable. Row 4 fails both the form and fit checks 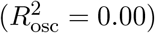the curve rises to unity and stays there, so the step response captures the exclusion transition but no periodic structure.

## S3. Segmentation comparison and recalibration

### S3.1. Examples on the Daniel *et al*. benchmark

Supplementary Fig. S3a shows representative images from the Daniel *et al*.^14^ benchmark, grouped into three categories: cases where CellADE succeeded and Cellpose-SAM returned no detections, cases where both performed comparably, and cases where CellADE over-detected because automatic scale estimation misjudged atypically large cells. In the last group, narrowing the LoG scale range from the estimated *σ* = [6, 18] to *σ* = [14, 16] raised F1 from 0.24 to 0.78, a correction path unavailable with a trained model.

### S3.2. Recalibration on clinical images

With automatic scale estimation, CellADE overcounted the clinical images relative to the device (Table 1). We estimated the expected LoG scale from the device-reported density: for CD= 2,506–3,623 cells/mm^2^, the mean cell area 10^6^/CD is 276–399 *µ*m^2^, the equivalent diameter 18.7–22.5 *µ*m (32–39 px at 0.58 *µ*m/px), and the LoG scale 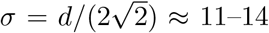 px. We tested this and nearby settings on the 350 images that disagreed most with the device (316 overcounted, 34 undercounted; Table S1). *σ* = 11–14 px overcorrected, while *σ* = 8–12 px with the LoG threshold lowered from 0.01 to 0.005 restored agreement; a further reduction to 0.003 changed little. The full dataset was then reprocessed with *σ* = 8–12 px and a threshold of 0.003 (Table 1). Supplementary Fig. S3b shows three clinical images that automatic scale estimation overcounted by 1.5–2.0×; after recalibration, CellADE came within 7% of the device count in each, whereas Cellpose-SAM detected only 20–26% of the cells in two of them.

**Figure S2.**
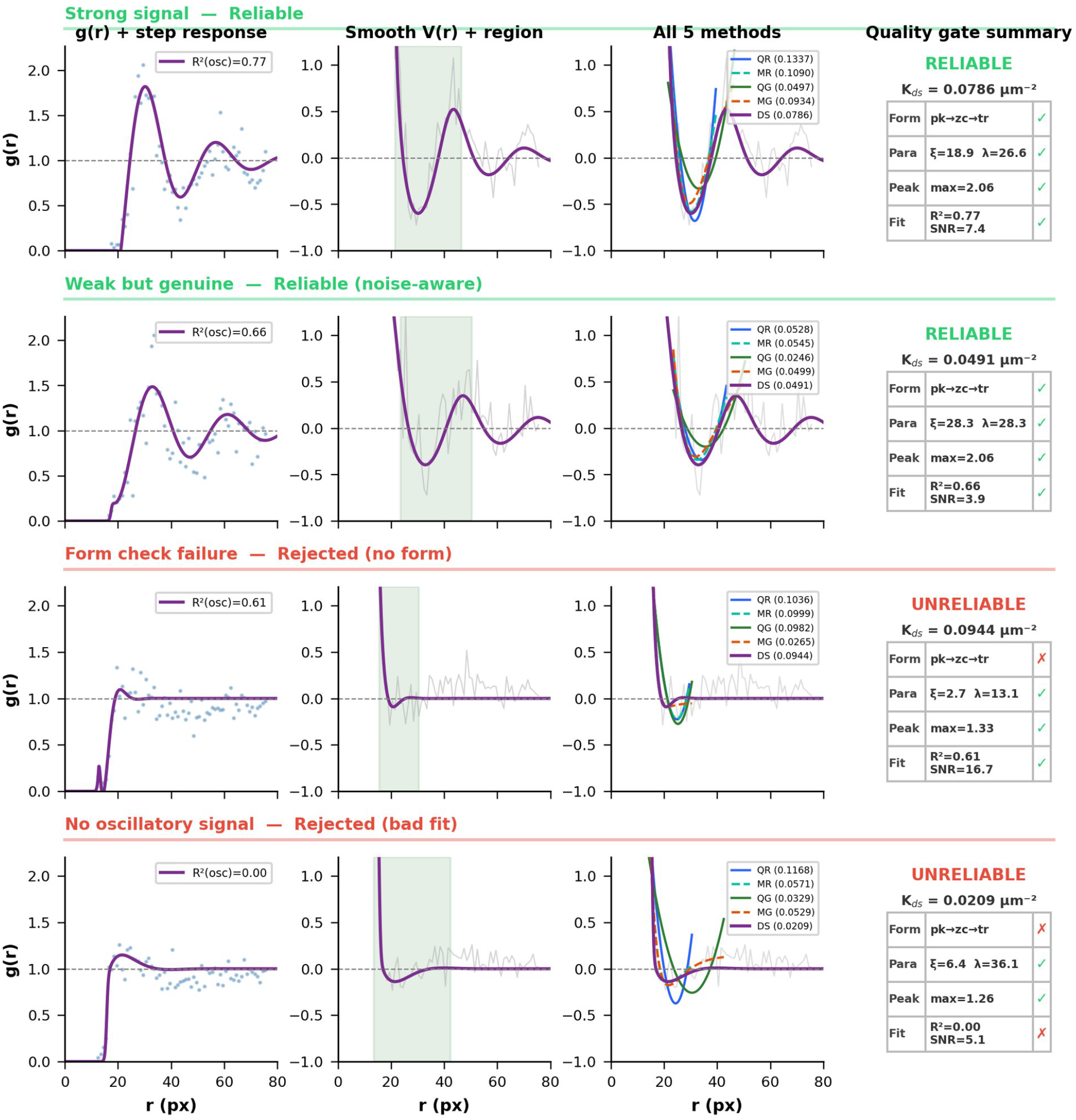
Representative fitting examples across reliability levels. Four rows show images spanning strong signal (top) to form check failure (bottom). **Column 1:** Raw *g*(*r*) with step response fit and 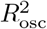. **Column 2:** Smooth *V* (*r*) derived from the fit with quality-gate-guided region (green shading). **Column 3:** All five extraction methods overlaid on raw *V* (*r*). **Column 4:** Quality gate verdict with diagnostic parameters. Rows 1–2 pass the composite reliability criterion; row 3 is rejected on the form check alone and row 4 on both the form and fit checks.

**Figure S3a.**
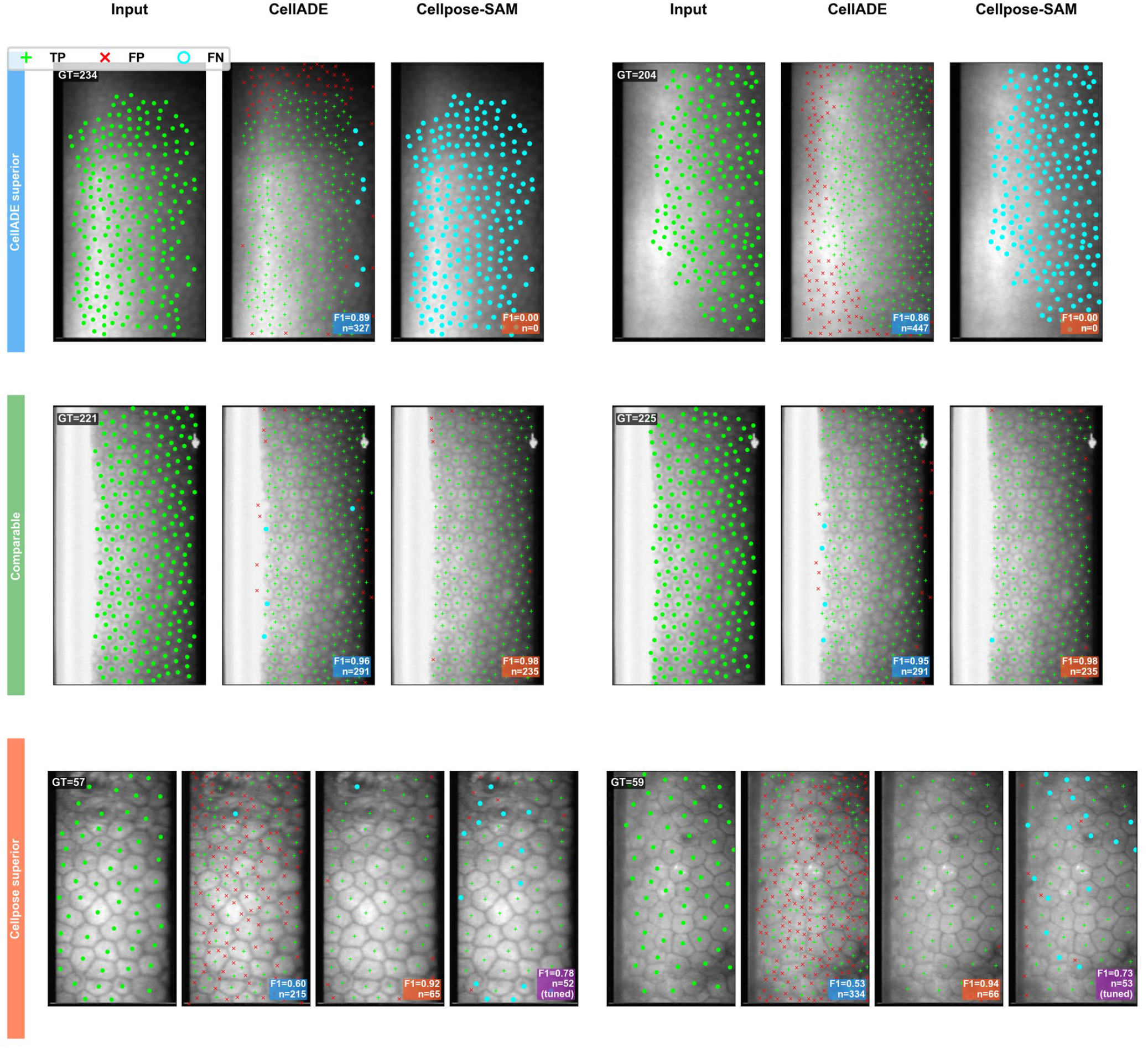
Segmentation comparison on the Daniel *et al*. benchmark. Six representative images grouped by performance category: CellADE superior, comparable, and Cellpose superior. Green crosses: true positives; red crosses: false positives; cyan circles: false negatives. Match radius = 15 px. In the Cellpose superior cases, adjusting a single parameter (LoG scale range *σ*) recovered detection performance (purple panels), demonstrating a correction path unavailable with opaque deep learning models.

**Table S1.** CellADE settings tested on the 350 clinical images that disagreed most with the device-reported density (count ratio to the device, *R*^2^ and Spearman *ρ* of per-image counts against the device).

| Setting | Median ratio to device | $R^2$ | Spearman $\rho$ |
| --- | --- | --- | --- |
| Automatic scale (FFT) | 1.41 | 0.05 | 0.38 |
| $\sigma$ 11–14 px, threshold 0.01 | 0.76 | 0.00 | 0.07 |
| $\sigma$ 8–12 px, threshold 0.005 | 1.03 | 0.25 | 0.50 |
| $\sigma$ 8–12 px, threshold **0.003** | **1.04** | **0.25** | **0.49** |

**Figure S3b.**
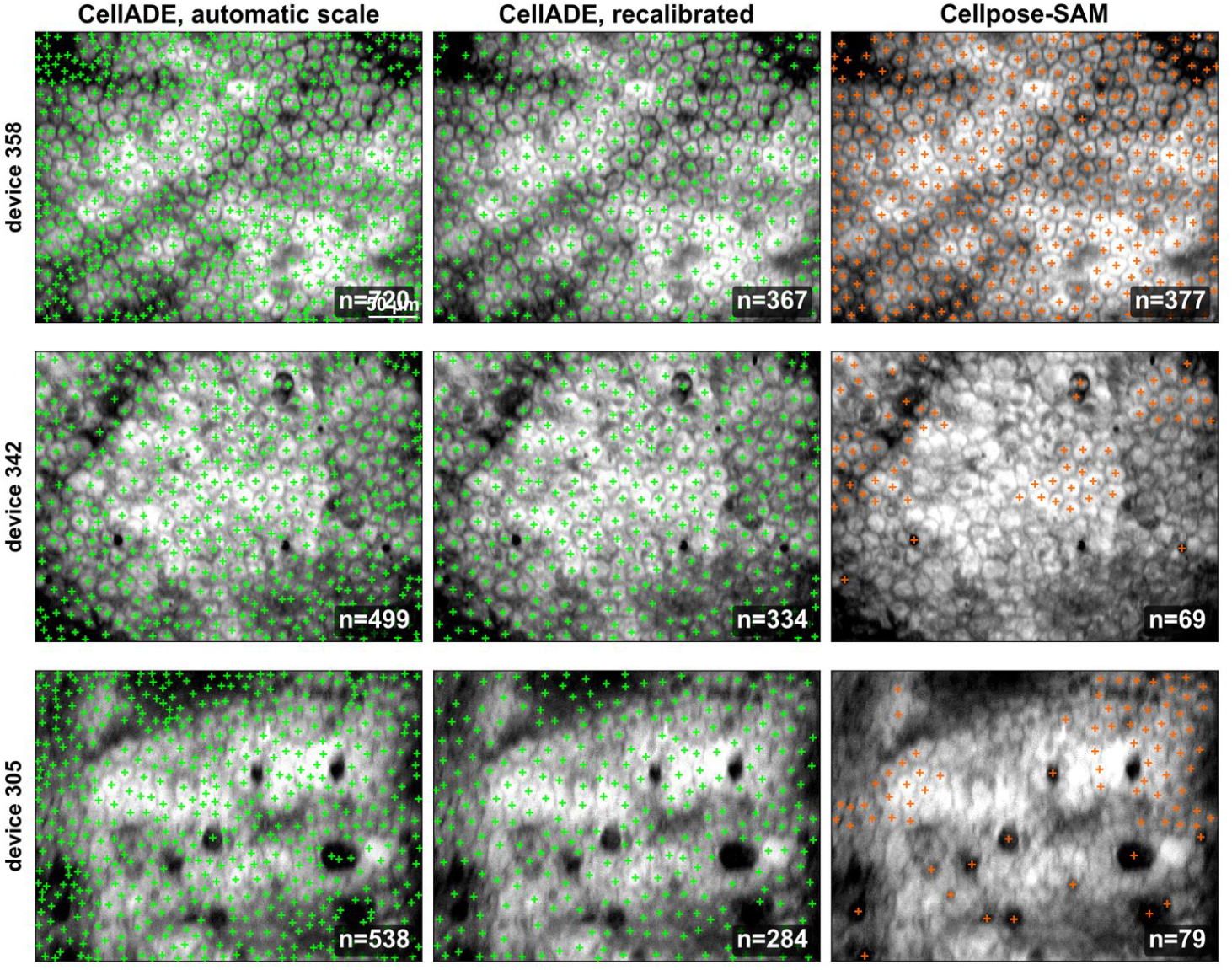
Recalibration of CellADE on clinical images. Three clinical images (rows) with CellADE detections using automatic scale estimation (left), after recalibration (*σ* = 8– 12 px, LoG threshold 0.003; middle), and Cellpose-SAM detections (right). The device count (density × imaged area) is given at the left of each row and the detected count in each panel. Automatic scale estimation overcounted all three images (720, 499 and 538 cells against 358, 342 and 305); after recalibration CellADE detected 367, 334 and 284. Cellpose-SAM matched the device in the first image (377) but detected only 69 and 79 cells in the other two. Scale bar: 50 µm; contrast adjusted for print.

### S4. Local rearrangement cannot reproduce the observed change in collective order

A change in collective order requires a field-reaching reorganization of all cells and cannot be produced by local adjustment alone. Since stripping produced significant changes in collective order, we tested whether a simulated local release could reproduce them.

Local release moved *K*_ds_ in the same direction as stripping in 60% of 782 donors, barely above the 54% expected by chance, since both mostly increased (Supplementary Fig. S4a–c).More importantly, it did not reproduce the forces behind the change. After real stripping, the field-wide mean force and *K*_ds_ rise together (*ρ* = 0.44; same direction in 69% of donors), whereas local release lowers the force (median −23%) while raising *K*_ds_, so the two move in opposite directions in two-thirds of donors (Supplementary Fig. S4d). The response to stripping is therefore a global reorganization of the monolayer, not a sum of local relaxations around stress hot spots.

### S5. Regression to the mean: correction and validation

Every donor cornea in this study was stripped exactly once, with one image taken before and one after, and no unstripped control group. This is a single-arm pre–post design. We want to know whether a donor’s baseline cell density or collective order predicts how it changes after stripping. The answer is not straightforward, because two artifacts contaminate every such analysis.

The first is mathematical coupling. The change Δ*Y* = *Y*_2_ − *Y*_1_ contains *Y*_1_ by construction, so regressing Δ*Y* on *Y*_1_ builds in a negative association mechanically, even if the intervention does nothing. The second is regression to the mean^19^: every measurement carries noise, so a donor whose baseline reading was unusually high was partly high by luck, and that luck does not repeat on the second measurement. On re-measurement, such donors drift back toward the population mean. The same happens in reverse for unusually low readings. Heterogeneous responses of individual donors to the same intervention have the same effect^S1^. Neither artifact requires anything to have changed; both produce the appearance of a restoring force pulling extreme tissues toward a common value.

The severity of both artifacts grows as the correlation between a variable’s baseline and follow-up values falls: with equal variances and no real effect, change and baseline still correlate at 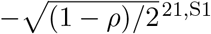. Here that correlation is moderate: *ρ*(*K*_BS_, *K*_AS_) = 0.47 and *ρ*(CD_BS_, CD_AS_) =0.51 (Olkin–Pratt corrected), which alone predicts spurious correlations of about −0.51 and −0.49, so the contamination is substantial. This section shows what the artifact looks like in our data, how the classical test for a differential effect and a per-donor correction separate it from real effects, and why the relationships reported in the main text survive its removal.

**Figure S4.**
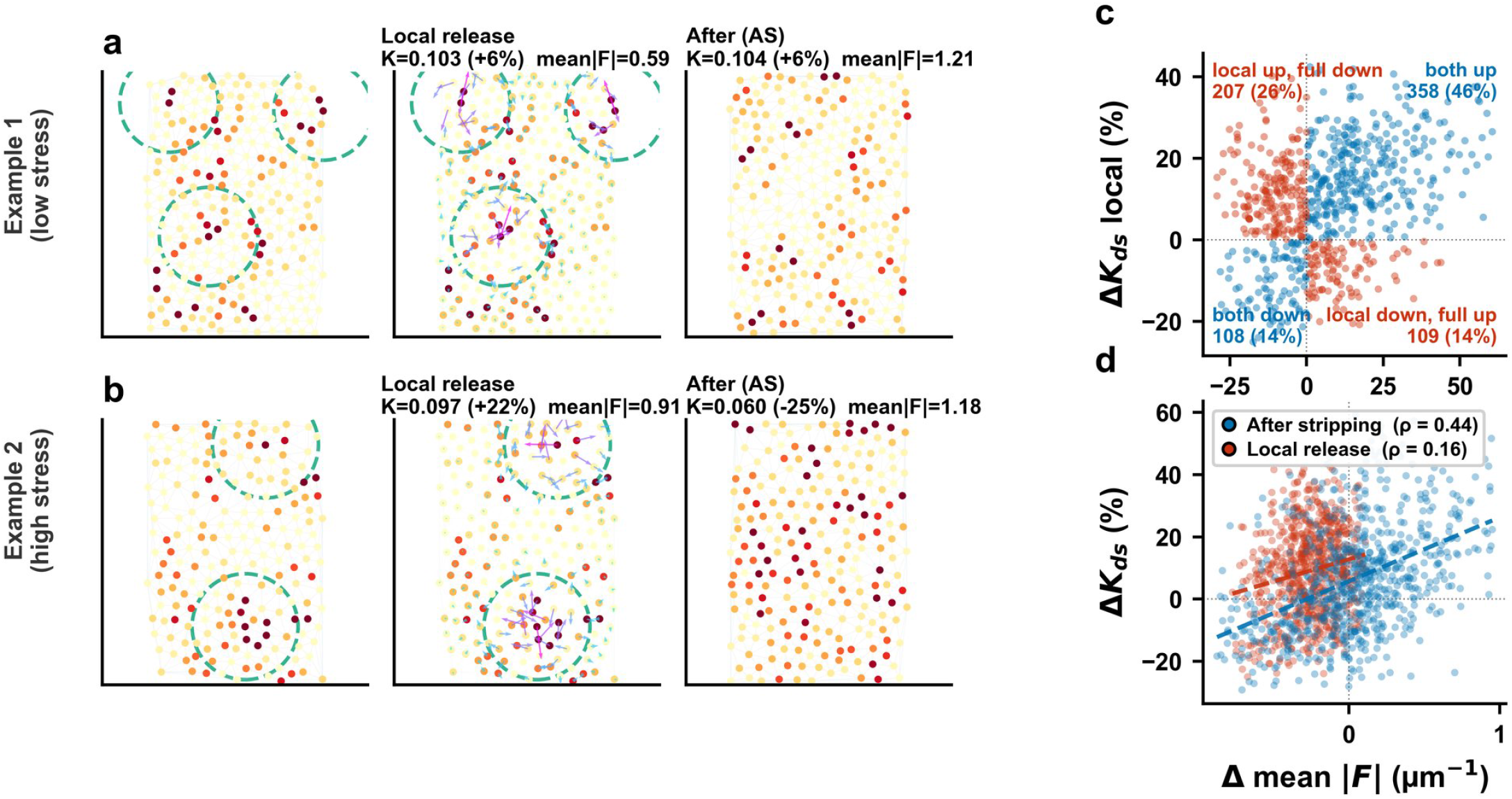
Local stress relief does not reproduce the observed reorganization. **(a**,**b)** Two representative donors, one starting from low and one from high mean force. Each row shows the baseline force field (left), the same tissue after the local-release perturbation with displacement vectors and release sites marked by dashed circles (centre), and the real tissue after stripping (right); *K*_ds_, its percentage change, and mean|*F* | are annotated. Node colour is per-cell residual force magnitude. **(c)** Percentage change in *K*_ds_ under local release against the change actually observed after stripping, one point per donor; colours mark donors in which the two changes agree (blue) or disagree (red) in sign, with the count in each quadrant. **(d)** Change in mean|*F* | against change in *K*_ds_ for the same donors, after real stripping (blue) and under local release (red), with least-squares trends dashed. Force and order rise together in the real tissue, whereas local release lowers the force while raising *K*_ds_. All forces in this figure are computed with the same reconstruction from the fitted *g*(*r*).the classical test for a differential effect and a per-donor correction separate it from real effects, and why the relationships

### S5.1. The artifact in our data

Regressing each metric’s change on its own baseline produces a strong negative slope in every case (Table S2). The uniformity across six different quantities, each measuring a different aspect of tissue organization, is itself the signature of an artifact rather than six independent biological findings. Each slope is close to the value expected from coupling and RTM alone,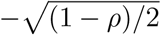given that metric’s own pre–post correlation.

**Table S2.** Naive correlation of change against own baseline for all six metrics (*n* = 750), and the value expected from coupling and RTM alone, 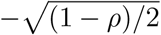 Every metric shows a strong negative slope close to that expectation.

| Metric | $\rho(Y_{BS}, Y_{AS})$ | Calculated naive $\rho(\Delta Y, Y_{BS})$ | Expected, no effect |
| --- | --- | --- | --- |
| $K_{ds}$ | +0.473 | −0.560 | −0.51 |
| CD | +0.510 | −0.422 | −0.49 |
| $r_0$ | +0.663 | −0.376 | −0.41 |
| HEX | +0.253 | −0.528 | −0.61 |
| CV | +0.457 | −0.505 | −0.52 |
| mean\|F\| | +0.226 | −0.628 | −0.62 |

Binning on baseline *K*_ds_ and measuring Δ*K*_ds_, the configuration in which the artifact is strongest, produces a monotonic trend from +0.014 *µ*m^−2^ in the lowest bin to −0.009 in the highest, with a rank correlation of *r* = −0.98 across bin medians (Supplementary Fig. S5a). Read at face value this would be a restoring force pulling every tissue toward a common value. As the table shows, the pre-post correlation for *K*_ds_ is *ρ* = 0.47, which alone predicts a naive slope near *ρ* − 1 = −0.53; the observed −0.56 is entirely consistent with this expectation. The correction of Section S5.4 removes it: after correction, the trend collapses to *r* = +0.12 (Supplementary Fig. S5d). What appears to be a strong biological signal is entirely attributable to the two artifacts.

### S5.2. Collective order converges beyond regression to the mean

The previous section showed that regression to the mean is present. The next question is whether anything real sits on top of it. Kelly & Price^S2^ distinguish two kinds of real effect. An additive effect shifts all donors equally: every cornea gains or loses the same amount, so the mean changes but the spread does not. A differential effect acts more strongly on some donors than others, depending on their baseline value, which changes the spread. RTM alone preserves the spread, so the test for a differential effect is whether the variance changed between baseline and follow-up. This is Oldham’s method, the recommended test of whether change depends on initial value^21,22,S1^. It can mislead when only donors with extreme starting values are studied, which is not the case here: every stripped cornea was included.

The null hypothesis is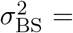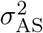, tested with Pitman’s paired statistic (equivalently, Oldham’s correlation between the change and the mean of the two measurements^S1^):

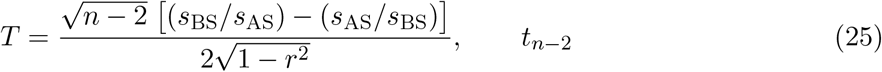

where *r* is the paired correlation. If the variances are equal, the observed slope of change on baseline is fully accounted for by RTM. If not, a differential effect is present, and its direction is read from the variance ratio *v* = *s*_AS_*/s*_BS_: *v >* 1 means the spread grew (divergence), *v <* 1 means it shrank (convergence).

**Table S3.** Variance-ratio test (Oldham’s method; Pitman’s paired statistic) on the trimmed analysis set (*n* = 750). *v* = *s*_AS_*/s*_BS_ is the variance ratio; *T* is Pitman’s paired statistic. Metrics whose variance changed significantly carry a differential effect on top of RTM; those that did not are fully explained by RTM alone.

| Metric | $s_{BS}$ | $s_{AS}$ | $v$ | $r$ | $T$ | $p$ | Verdict |
| --- | --- | --- | --- | --- | --- | --- | --- |
| $K_{ds}$ | 0.0136 | 0.0126 | 0.929 | +0.473 | +2.28 | 0.023 | converged |
| HEX | 4.758 | 5.265 | 1.106 | +0.253 | −2.86 | 0.004 | diverged |
| CD | 190.5 | 200.4 | 1.052 | +0.511 | −1.60 | 0.109 | RTM alone |
| $r_0$ | 0.656 | 0.676 | 1.031 | +0.664 | −1.10 | 0.270 | RTM alone |
| CV | 2.722 | 2.734 | 1.004 | +0.457 | −0.14 | 0.893 | RTM alone |
| mean$\vert F\vert $ | 0.0865 | 0.0844 | 0.975 | +0.226 | +0.71 | 0.476 | RTM alone |

Two metrics show a differential effect, in opposite directions (Table S3; Supplementary Fig. S5c). *K*_ds_ converged: its spread shrank after stripping (*v* = 0.93, *P* = 0.023), so tissues moved toward a more common level of order beyond what regression to the mean produces. Hexagonality diverged (*v* = 1.11, *P* = 0.004). CD, *r*_0_, CV and mean|*F* | show no differential effect; their naive slopes are fully accounted for by RTM. Collective order is thus the only metric that converges after stripping, consistent with its being the metric most responsive to it (main text, Fig. 5f).

The per-donor correction used for the cross-variable analyses (main text, Fig. 5h,i; *k* = 1, Section S5.5) subtracts from each donor’s change the part expected from RTM^S2^. After this correction, every metric’s slope against its own baseline falls to within ±0.06 of zero (Table S4), so the strong naive slopes of Section S5.1 are removed before donors are binned on another variable. The small differential effects detected above (convergence of *K*_ds_, divergence of hexagonality) are too weak to appear as a slope after this correction.

**Table S4.** Per-donor correction (*k* = 1; Kelly & Price per-case adjustment, *n* = 750). After removing the RTM component, the correlation between corrected change and own baseline collapses for every metric.

| Metric | $\hat{r}$ | Naive $\rho$ | Adjusted $\rho$ | $p$ |
| --- | --- | --- | --- | --- |
| $K_{\text{ds}}$ | +0.473 | −0.560 | −0.033 | 0.37 |
| CD | +0.510 | −0.422 | +0.057 | 0.12 |
| $r_0$ | +0.663 | −0.376 | +0.025 | 0.50 |
| HEX | +0.253 | −0.528 | +0.036 | 0.32 |
| CV | +0.457 | −0.505 | +0.013 | 0.73 |
| mean$\vert F\vert $ | +0.226 | −0.628 | −0.018 | 0.62 |

### S5.3. Cross-variable binning

The artifact requires the binning variable and the measured variable to share their noise. Binning on one baseline quantity and measuring the change in a different one removes that shared-noise pathway only if the two are weakly correlated at baseline; otherwise the artifact can leak from one variable into the other^40^. Table S5 shows the baseline cross-correlations for all pairs.

Cell density and *K*_ds_ share 29% of their baseline variance, so binning one against the other’s change does not by itself remove the artifact: it leaks through the shared variance, and cross-variable binning therefore needs the same correction as own-baseline analyses. Because the two are positively correlated, the leak has a fixed direction. Donors in the dense bins also tend to have high baseline *K*_ds_, which regresses downward at follow-up; donors in the sparse bins tend to have low baseline *K*_ds_, which regresses upward. The artifact therefore lowers the apparent change in order in dense tissue and raises it in sparse tissue, producing a spurious negative trend of Δ*K*_ds_ against cell density. The same holds in reverse for ΔCD against baseline *K*_ds_. The leak can push the cross-variable trends downward, never upward. The uncorrected trend is accordingly negative: Δ*K*_ds_ against baseline cell density, *r* = −0.50 (Supplementary Fig. S5b). Cell density and *r*_0_ share 37% of their variance, so binning one against the other’s change was not used.

**Table S5.** Baseline cross-correlations between all metric pairs (*n* = 753). Pairs with low *R*^2^ are safe for cross-variable binning; pairs with high *R*^2^ are not, because indirect leakage through the shared variance reintroduces the artifact.

| Pair | $r$ | $R^2$ | $p$ | Safe for cross-binning? |
| --- | --- | --- | --- | --- |
| CD vs $K_{\text{ds}}$ | +0.534 | 0.286 | $7.5 \times 10^{-57}$ | needs correction |
| $K_{\text{ds}}$ vs $r_0$ | -0.184 | 0.034 | $3.5 \times 10^{-7}$ | yes |
| CD vs HEX | +0.181 | 0.033 | $5.6 \times 10^{-7}$ | yes |
| $K_{\text{ds}}$ vs HEX | +0.397 | 0.158 | $6.7 \times 10^{-30}$ | marginal |
| $r_0$ vs HEX | +0.230 | 0.053 | $1.6 \times 10^{-10}$ | yes |
| CD vs $r_0$ | -0.606 | 0.368 | $9.5 \times 10^{-77}$ | no |

### S5.4. Correcting each donor’s change without a control arm

Sections S5.2 and S5.3 showed that each donor’s change has to be corrected for the artifact before donors are binned, whether on their own baseline or on another variable. The correction subtracts from each change the part that the artifact alone would have produced. Ideally, that part would be read from a control arm, the same donors imaged twice without stripping, which this study does not have. Varadhan *et al*. ^20^ showed that the missing control arm enters the correction through a single constant, *k*: the ratio between the baseline–follow-up covariance that would have been observed without the intervention and the one actually observed. Because single-arm data contain no information on *k*, they treat it as unknown and examine a range of plausible values. Without stripping, however, baseline and follow-up are simply two measurements of the same tissue, so their covariance is set by how closely repeated measurements agree. That agreement can be measured within our own data: the two halves of each image are two simultaneous reads of the same tissue. The correction then follows from their equations with *k* measured rather than assumed. This section derives it.

#### Setup

Let *Y*_1_ and *Y*_2_ denote the baseline and follow-up values of a metric (e.g., *K*_ds_). If the two are bivariate normal with means *µ*_1_, *µ*_2_, standard deviations *σ*_1_, *σ*_2_, and correlation *ρ*, the conditional expectation of *Y*_2_ given *Y*_1_ = *y*_1_ is:

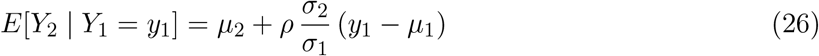

Subtracting *y*_1_ from both sides gives the expected change:

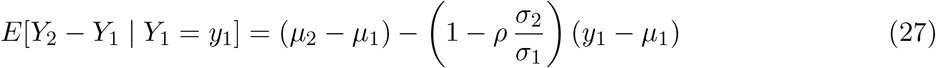

This is a linear function of *y*_1_ with slope *ρ σ*_2_*/σ*_1_ − 1. Since 0 ≤ *ρ* ≤ 1 in this context and*σ*_2_*/σ*_1_ is typically close to 1, the slope is almost always negative. If the intervention has no differential effect, the two variances are equal and the slope reduces to *ρ* − 1: negative for any imperfect correlation, and −1 when baseline and follow-up are uncorrelated, so that a donor one unit above the mean at baseline appears to lose one full unit. This part of the slope is the artifact; only a departure of *σ*_2_*/σ*_1_ from 1 reflects a real differential effect.

#### The counterfactual control group

Let 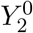 denote the counterfactual follow-up of the same donors, not stripped. We assume the control arm shares the same baseline distribution 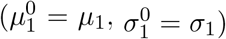, and write the expected change in the control arm by the same logic:

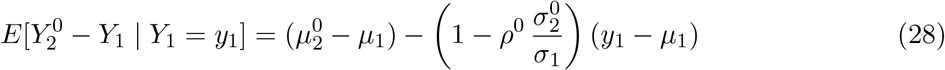

The real treatment effect at baseline *y*_1_ is the difference between the treatment and control changes:

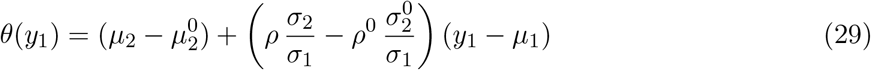

The artifact cancels in the subtraction. What remains is genuine.

#### Reducing to a single parameter

The control-arm parameters *ρ*^0^ and 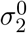are unobservable.They can be combined into a single constant:

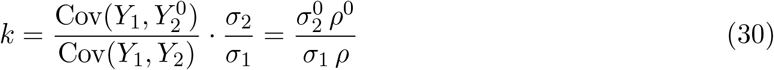

which compares how tightly baseline and follow-up would track each other without the intervention with how tightly they track with it. This simplification does not require normality. Writing the control and treatment arms as linear models:

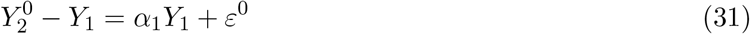

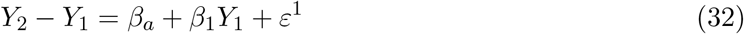

and assuming only that the control-arm error is uncorrelated with baseline (*E*[*ε*^0^ | *Y*_1_] = 0), solving for *α*_1_ gives:

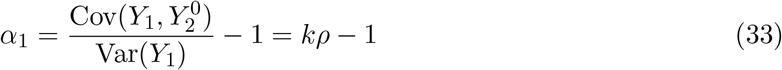

The −1 is the mathematical coupling. The *kρ* is the regression to the mean, scaled by *k* because we do not know how tightly baseline and follow-up would have tracked each other without the intervention. The corrected slope is 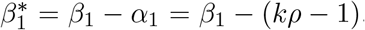 . Without stripping, the follow-up has the same spread as the baseline (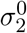 =_1_), and *ρ*^0^ is the agreement between repeated measurements of the same tissue, its reliability *R*; hence *k* = *R/ρ*.

#### With covariates

When a covariate *X* (e.g., baseline cell density) is included, *Y*_1_ is correlated with *X*, so the biases on the baseline coefficient and the covariate coefficient become entangled. The full model is:

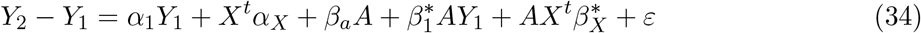

where *A* ∈ {0, 1} indicates treatment versus control. In the single-arm study we observe only the combined coefficients *β*_1_ = *α*_1_ + 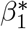 and *β*_*X*_ = *α*_*X*_ + 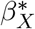 . Solving the estimating equations under the same exogeneity assumption, and introducing a second parameter *m* (set to 1) that assumes therelationship between covariates and outcome does not change without intervention, the bias terms are:

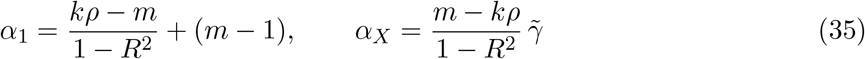

where *R*^2^ and 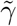 come from regressing *Y*_BS_ on *X*_BS_. When *m* = 1 and there are no covariates (*R*^2^ = 0), this reduces to *α*_1_ = *kρ* − 1, recovering the simple case above. The term *α*_*X*_ is the formal counterpart of the leak described in Section S5.3: it is the part of the artifact that a covariate correlated with the baseline absorbs.

#### Estimation

The sample correlation *r* and sample *R*^2^ are biased estimators. We use the Olkin– Pratt unbiased estimators:

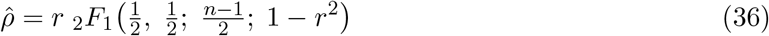

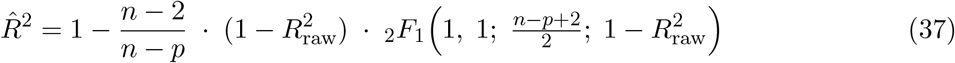

where 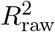is the sample coefficient of determination from regressing *Y*_BS_ on *X*_BS, 2_*F*_1_ is the Gauss hypergeometric function, *n* is the sample size, and *p* is the number of covariates. For our sample sizes (*n* = 753) the correction is small but we include it for completeness. The corrected coefficients are 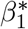 = *β*_1_ − *α*_1_ and 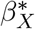 = *β*_*X*_ − *α*_*X*_.

#### Measuring *k*

The parameter *k* cannot be estimated from the pre–post data alone, but it can be measured from the images themselves. We split each image into left and right halves (342 × 514 px each) and recomputed *K*_ds_ and cell density on each half. The two halves are simultaneous measurements of the same tissue, so their agreement across images reflects measurement noise only. The correlation between halves, *r*_half_, gives the reliability of the full-image value by the Spearman– Brown formula, *R* = 2*r*_half_ */*(1 + *r*_half_ ), and *k* = *R/ρ*. For *K*_ds_, *R* = 0.46 (all 1,605 reliable images), giving *k* = 0.46*/*0.47 ≈ 1.0; for cell density, *R* = 0.87, giving *k* ≈ 1.7. Splitting into top and bottom halves gave the same reliability for cell density, but halves only 257 px tall proved too short for a stable fit of *g*(*r*), so *K*_ds_ was measured on left–right halves only. For *K*_ds_ the measured value thus coincides with *k* = 1, the classical correction^21,22^.

The derivation follows Varadhan *et al*. ^20^; our implementation is in Python, reimplementing the R package resilience with the addition of the Olkin–Pratt unbiased estimators for *ρ* and *R*^2^, and with *k* measured from split images rather than assumed.

### S5.5. The correction applied to our data

With *k* = 1 measured for *K*_ds_ (Section S5.4), each donor’s change in *K*_ds_ is corrected as 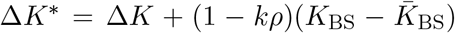. The same-variable trend then collapses: Δ*K*_ds_ binned on baseline *K*_ds_ goes from *r* = −0.98 before correction to *r* = +0.12 after (*P* = 0.78; Supplementary Fig. S5a,d), so the artifact is removed. Binned on baseline cell density instead, the corrected change in *K*_ds_ rises with density, from *r* = −0.50 before correction to *r* = +0.86 after (*P* = 0.007; Supplementary Fig. S5b,e). This is the trend reported in the main text (Fig. 5h), where the same corrected changes are summarized by decile as the share of donors whose order rose and the median percentage change.

**Figure S5.**
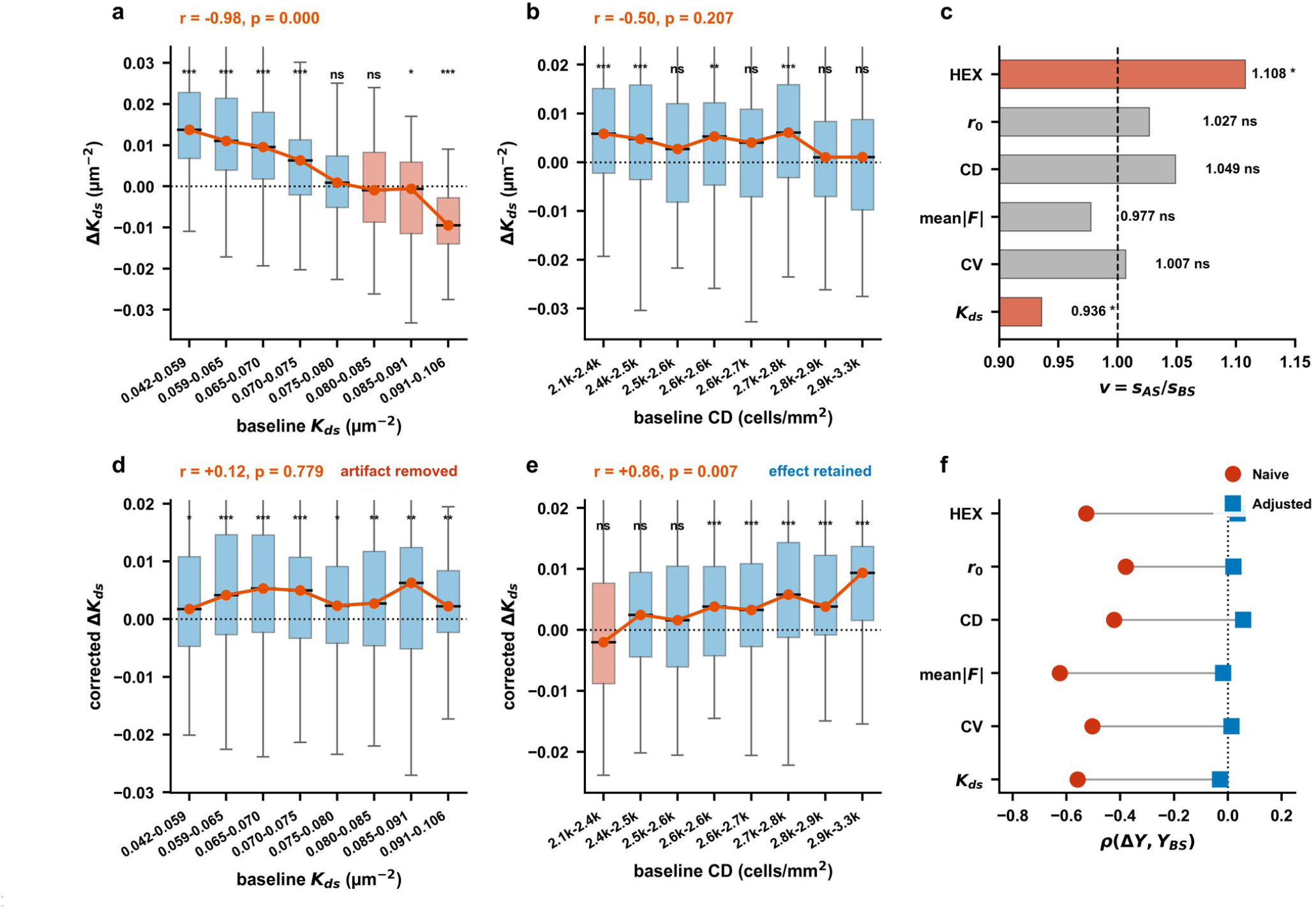
Regression to the mean in the paired donors, and its correction. **(a)** Δ*K*_ds_ binned on baseline *K*_ds_ (same variable); the paired correlation *ρ*(BS, AS) is annotated. **(b)** Δ*K*_ds_ binned on baseline cell density (cross-variable); the baseline *R*^2^ between the two variables is annotated. Boxes show the distribution of the change within each bin, coloured by the sign of the bin median; the line joins the medians; *r* and *p* are the rank correlation across bin medians; stars give the Wilcoxon test of each bin against zero. **(c)** Variance ratio *v* = *s*_AS_*/s*_BS_ for all six metrics (Oldham’s variance-ratio test, Pitman’s statistic); red bars mark a differential effect on top of RTM (*p <* 0.05), grey bars RTM alone. **(d**,**e)** As (a,b) after the per-donor correction at the measured *k* = 1 (Sections S5.4, S5.5): **(d)** the same-variable trend collapses (*r* = +0.12, *P* = 0.78); **(e)** the cross-variable trend becomes positive (*r* = +0.86, *P* = 0.007). **(f)** Naive correlation between change and own baseline (red circles, *ρ* ≈ −0.38 to −0.63) and after the per-donor correction (blue squares), for all six metrics.

## S6. Segmentation error does not create the Δ*K*_ds_–ΔCD correlation

### S6.1. The problem

*K*_ds_ and cell density are both computed from the same cell centroids, so a single segmentation error (a missed cell or a false detection) changes both at once. Each measured value is the true value plus an error,

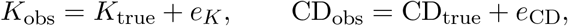

and because both come from the same centroids, *e*_*K*_ and *e*_CD_ are correlated. For each BS/AS pair, the measured change is the true change plus the difference in errors:

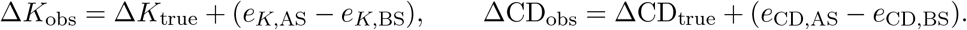

If the BS and AS errors are independent of each other and of the true values, the observed covariance splits into two parts:

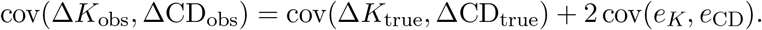

The first part is the real co-change; the second comes only from shared segmentation error. The question is whether the second part could explain the observed correlation. This is a different problem from regression to the mean (Section S5), which concerns a single variable’s change compared with its own baseline. Here, two different variables share the same error within each image ^41^.

### S6.2. Method

We used simulation–extrapolation (SIMEX; Methods). We added extra segmentation error to the real BS and AS centroids, independently, at increasing levels, recomputed the correlation each time, and followed how it changed.

### S6.3. How to read the result

- If the correlation gets weaker as error is added, error is hiding part of the real relationship, and the measured value is an underestimate.
- If it gets stronger, error is creating part of the relationship, and the measured value is inflated.
- If it stays the same, the measured value is not affected by that error.

### S6.4. Results

We added two kinds of segmentation error and watched what happened to the correlation between Δ*K*_ds_ and ΔCD (*ρ* = +0.26 in the real data).

#### Missed cells

Removing up to 20% of cells at random changed the correlation very little,from +0.26 to +0.23. Projecting back to the error-free case gave +0.26 to +0.27.*False detections*. Adding fake cells weakened the correlation sharply, to about +0.07 already at 5%, after which it flattened. Because this curve falls steeply at first and then flattens, projecting it back to the error-free case is unreliable, so we use only its direction (all projections stay positive).

In both cases, adding error made the correlation weaker, never stronger. Segmentation error therefore cannot create the observed correlation; if anything, the true correlation is at least as strong as measured. The result was the same whether or not images that failed the quality gate after error was added were excluded (Supplementary Fig. S6). Supplementary Fig. S6c also offers indirect support for the method itself: removing real cells left nearly all images passing the quality gate (≤1%), whereas adding randomly placed points caused increasingly many to fail (up to 40%), indicating that the damped oscillatory *g*(*r*) arises from the spatial arrangement of real cells and that the gate detects its loss.

## S7. CellADE segmentation algorithm

### S7.1. Pipeline overview

Supplementary Fig. S7 illustrates the CellADE pipeline on a benchmark image containing a non-cellular bright region along its left edge, chosen because it exercises the region-selection and scale-estimation stages. The pipeline proceeds through three stages, followed by a gap-filling pass described in Section S7.7:

*Preprocessing* (panels a–d): The raw image is contrast-enhanced via CLAHE. Strip-based region-of-interest detection divides the normalized (pre-CLAHE) image into eight vertical strips. The top and bottom 10% of each strip are trimmed to exclude edge artefacts, and the mean absolute first derivative (“jaggedness”) of each strip’s vertical intensity profile is computed. Strips whose jaggedness falls below a threshold, taken as the greater of an absolute floor (0.002) and 40% of the maximum jaggedness, are classified as non-cellular. Seeds subsequently detected inside these strips are discarded.

*FFT sigma estimation* (panels e–h): A centre crop (35% area) positioned at the centroid of cell-containing strips is mirror-tiled into a 4 × 4 mosaic to increase frequency resolution. The 2D power spectrum is computed, radially averaged, whitened by 1*/f* trend removal, and low-pass filtered (5 Fourier coefficients). The dominant peak frequency is converted to the LoG scale parameter *σ*, which sets the adaptive scale range for seed detection. When fewer than 75% of strips are cell-containing, the spectral estimate is skipped and the default range is used instead.

**Figure S6.**
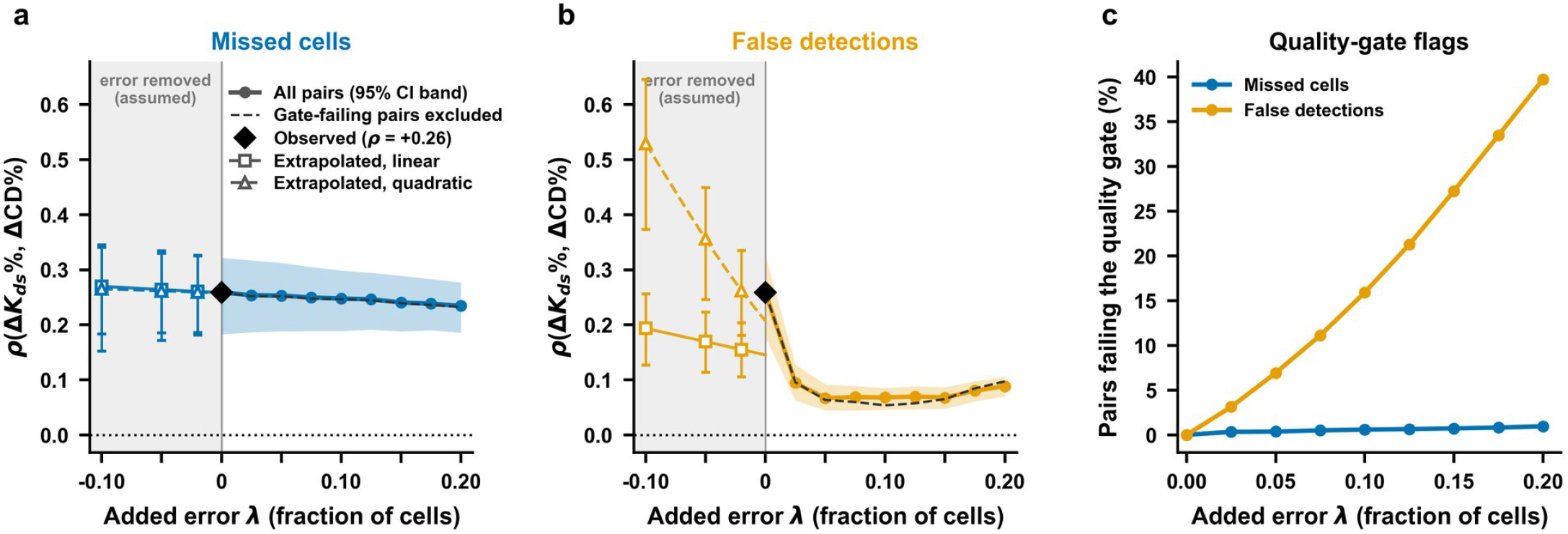
Segmentation error weakens, and does not create, the Δ*K*_ds_–ΔCD correlation (SIMEX). **(a)** Correlation between Δ*K*_ds_% and ΔCD% as missed cells are added at random to the BS and AS centroids of the 779 reliable pairs (*λ*, fraction of cells; 100 replicates per level). Solid line and band, mean and 95% bootstrap confidence interval over all pairs; dashed black line, pairs that failed the quality gate after error was added excluded; black diamond, observed value. The shaded region shows extrapolation back to assumed existing error rates of 2, 5 and 10% (linear, squares; quadratic, triangles; Fisher-*z* scale). **(b)** The same for false detections added at random positions. **(c)** Percentage of perturbed pairs in which the BS or AS image failed the quality gate.

*Segmentation* (panels i–l): Laplacian-of-Gaussian blob detection identifies cell seed positions.Local fuzzy C-means clustering extracts per-cell foreground masks. Marker-controlled watershed produces the labelled segmentation, from which cell centroids are extracted for downstream *g*(*r*) computation.

### S7.2. Preprocessing

The input image is converted to grayscale and normalized to [0, 1]:

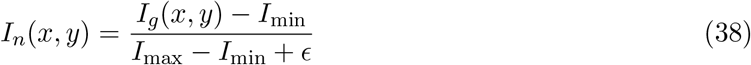

where *ϵ* = 10^−8^. Contrast-limited adaptive histogram equalization (CLAHE) is applied with clip limit *c* = 0.03:

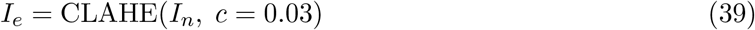

CellADE detects cells as bright objects on a dark background, which is how they appear in specular microscopy. For modalities with the opposite polarity, such as bright-field images of cultured cells, an optional inversion step is applied before CLAHE and the image is re-normalized. Inversion is disabled for both clinical cohorts in this study.

### S7.3. Spectral cell-size estimation

The LoG detector requires a scale range [σ_min_, *σ*_max_] that brackets the expected cell size. A fixed range either misses cells outside the assumed size or wastes computation searching irrelevant scales. We estimate the dominant cell size from the image itself using the radial power spectrum, then derive an adaptive sigma range.

#### Adaptive cropping

Specular microscopy images often contain non-cellular regions (bright reflections, empty margins) at the periphery. To ensure the frequency analysis captures cell periodicity rather than large-scale illumination structure, a crop of area fraction *f* = 0.35 is extracted and centred at the horizontal centroid of cell-containing strips (identified by the strip jaggedness analysis) and the vertical midpoint of the image:

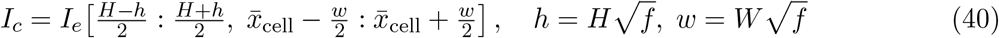

where 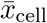 is the mean horizontal position of strips classified as cell-containing. This positions the frequency analysis over the densest cell region rather than the geometric image centre, which may contain non-cellular areas in partial-coverage fields.

**Figure S7.**
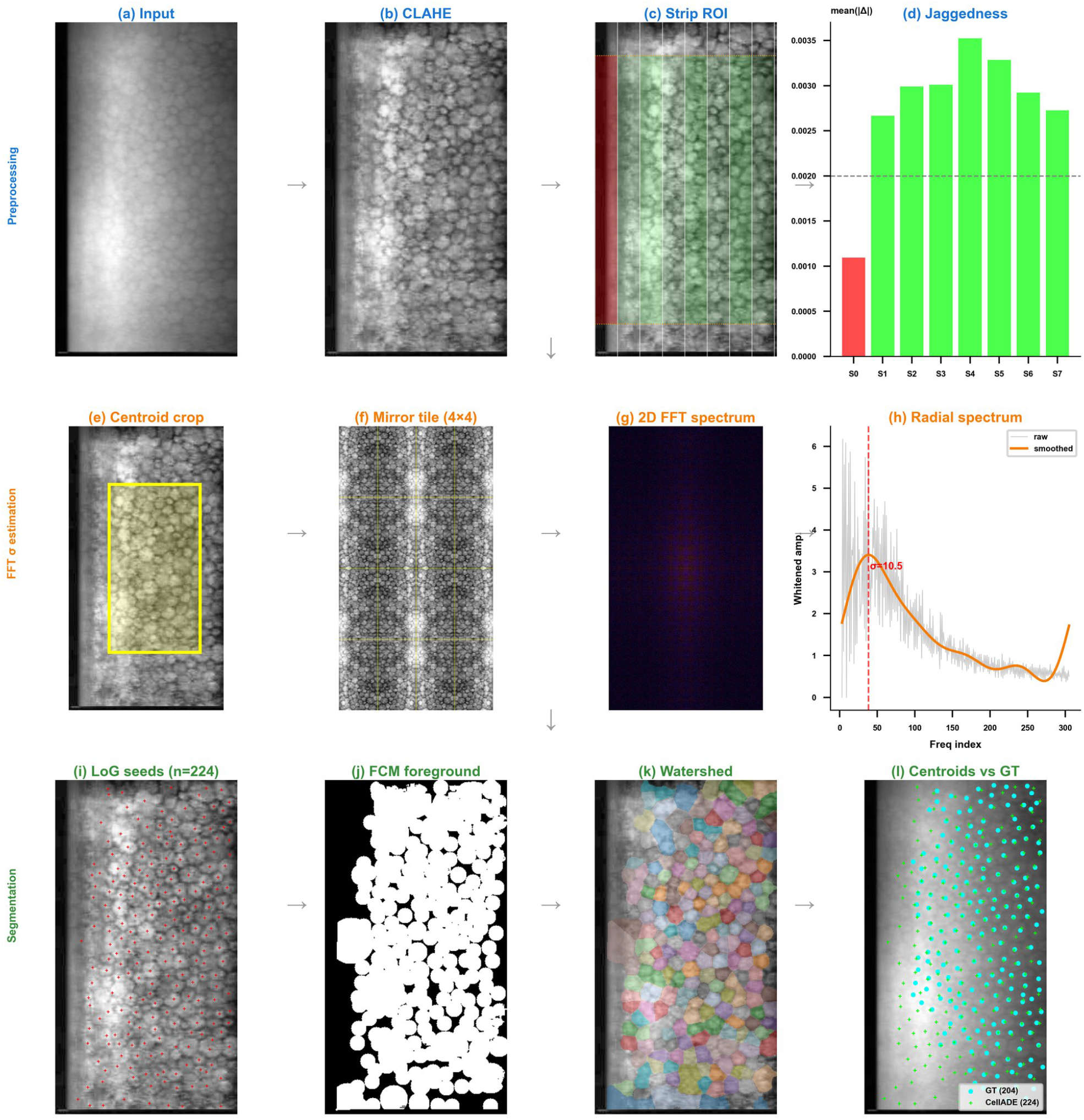
CellADE cornealia pipeline. Processing stages illustrated on image 72324, which contains a non-cellular bright region (left side). **Row 1, Preprocessing:** (a) Input image. (b) CLAHE-enhanced. (c) Strip ROI detection (green = cell-containing, red = excluded).(d) Jaggedness metric per strip; dashed line indicates threshold. **Row 2, FFT sigma estimation:** (e) Centre crop positioned at cell-strip centroid. (f) 4 × 4 mirror-tiled mosaic. (g) 2D power spectrum. (h) Whitened radial spectrum with peak detection yielding = 10.5. **Row 3, Segmentation:** (i) LoG seed detection. (j) FCM foreground mask. (k) Watershed segmentation. (l) Final centroids (green) overlaid on ground truth (cyan).

#### Mirror tiling

Fourier analysis requires sufficient repetitions of the periodic pattern to resolve its frequency. For large cells, the crop may contain only 3–4 cells, yielding an ambiguous spectral peak. To increase frequency resolution without introducing seam artefacts, the crop is mirror-tiled into an *n* × *n* mosaic (*n* = 4 by default): odd-indexed copies are reflected horizontally or vertically so that cell boundaries align seamlessly across tile edges. The tiled image *I*_*t*_ has dimensions *nH*_*c*_ × *nW*_*c*_ and *n*^2^× finer frequency bins than the original crop.

#### Radial power spectrum

The 2D discrete Fourier transform of the mean-subtracted tiled image is computed and shifted so that the zero-frequency component is centred:

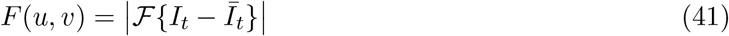

The radial average of *F* yields a one-dimensional power profile *P* (*k*), where *k* is the radial frequency index:

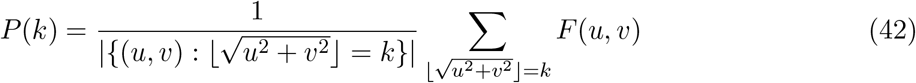

#### Spectral whitening

Natural images exhibit a characteristic 1*/f* ^*α*^ power-law decay that dominates the radial profile at low frequencies, masking the cell-periodicity peak. To remove this trend, a linear fit in log–log space is computed for *k* ≥ *k*_min_ (where *k*_min_ = max(3, ⌊0.01 · *k*_max_⌋) excludes the DC neighbourhood):

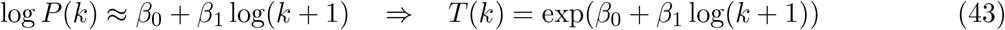

The whitened profile is obtained by dividing out the trend:

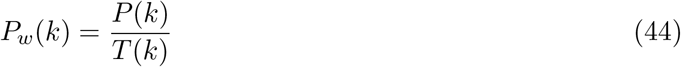

In *P*_*w*_, the cell-periodicity signal appears as a peak above a flat baseline rather than a bump on a steep slope.

#### Low-pass smoothing

The whitened profile is noisy at high frequencies. Rather than a moving average, which is sensitive to local outliers, we apply a spectral low-pass filter: the one-dimensional FFT of *P*_*w*_ is computed, and only the lowest *n*_keep_ = 5 Fourier coefficients are retained before inversetransformation:

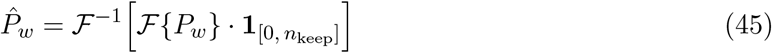

This extracts the smooth envelope of the whitened spectrum, capturing the overall spectral trend while suppressing noise.

#### Peak detection

The peak of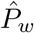 corresponds to the dominant spatial periodicity. To avoid selecting residual low-frequency energy that survives whitening, the search begins only after 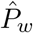 first exceeds its own median value (skipping the initial rising portion):

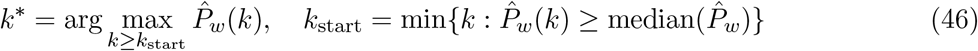

#### Sigma conversion

The peak frequency index *k*^∗^ is converted to a spatial scale. The cell-to-cell spacing in pixels is:

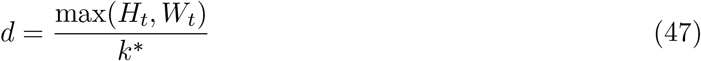

where *H*_*t*_, *W*_*t*_ are the tiled image dimensions. The estimated cell radius and corresponding LoG sigma are:

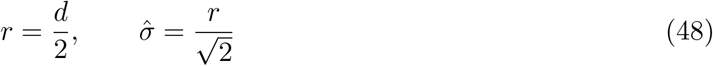

The factor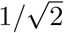arises because the LoG filter achieves its maximum response when 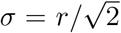for a circular blob of radius *r*.

#### Adaptive range and outlier handling

The LoG scale range is set symmetrically around 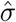:

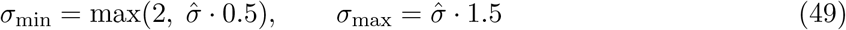

Images with poor cell contrast, partial ROI coverage, or dominant non-cellular structure can produce unreliable spectral peaks, typically manifesting as anomalously large 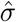 values. To handle such outliers, a hard clamp is applied: if 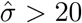 (corresponding to a cell radius *>* 28 px, which exceeds any plausible corneal endothelial cell in these images), the estimate is discarded and replaced by a data-driven calibrated range.

The calibrated range is derived from ground-truth centroid annotations across the benchmark dataset. For each annotated image, the nearest-neighbour distance *d* is computed for every cell centroid, and each is converted to a scale by 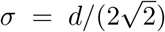 as above. The 5th and 95th percentiles of the resulting distribution across all cells define a population-level reference range [σ_cal,min_, σ_cal,max_]. In the present dataset (*N* = 42,147 cells across 324 images), this yielded [σ_cal,min_, σ_cal,max_] = [4.5, 11.7], corresponding to cell spacings of 12.7 to 33.1 px. When the FFT estimate exceeds the hard clamp, the pipeline falls back to this range, so images where spectral estimation fails still receive a data-informed scale range rather than arbitrary defaults.

### S7.4. Seed detection via Laplacian-of-Gaussian

Cell centers are detected as bright blobs using multi-scale Laplacian-of-Gaussian (LoG) filtering, as previously established for sub-resolution adhesion and bead localization ^S3,S4^. For each scale σ, the scale-normalized LoG response is:

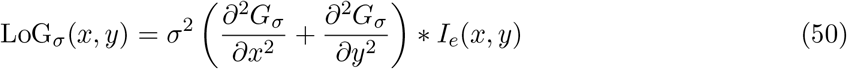

where *G*_*σ*_ is a Gaussian kernel with standard deviation σ. Local maxima across the three-dimensional scale space (*x, y*, σ) that exceed a response threshold *τ* yield a set of blob detections:

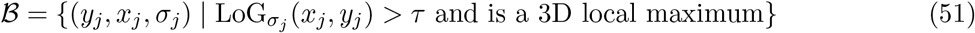

The sigma range [σ_min_, σ_max_] is determined adaptively by the spectral estimator (Section S7.3) when available, or set to default values (σ_min_ = 6, σ_max_ = 30, 15 scales, *τ* = 0.01).

### S7.5. Local fuzzy C-means foreground extraction

Rather than applying a global intensity threshold, CellADE performs two-class fuzzy C-means (FCM) clustering locally around each detected seed. For each blob *j*, a circular patch of radius 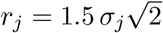 is extracted. The FCM objective minimizes:

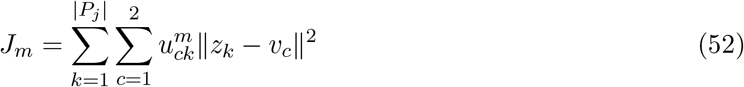

where *z*_*k*_ is the pixel intensity, *v*_*c*_ is the cluster centre, *u*_*ck*_ ∈ [0, 1] is the fuzzy membership satisfying 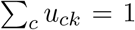, and *m* = 2 is the fuzziness exponent. Pixels with membership *>* 0.4 in the brighter cluster are classified as cell interior. The per-seed binary masks are combined via logical OR into a global foreground map **F**.

This local approach is the key design element of CellADE: each cell’s foreground is determined relative to its own local intensity distribution, making the algorithm robust to intensity gradients across the field of view that defeat global thresholding methods.

### S7.6. Marker-controlled watershed segmentation

The detected blob centres serve as markers on the global foreground mask **F**. A Euclidean distance transform from the markers provides the flooding landscape for the watershed algorithm^39^, producing a labelled image **L** where each cell receives a unique integer label.

### S7.7. Intercellular gap filling and guttae protection

The watershed of Section S7.6 is confined to the FCM foreground, so the thin dark boundaries between cells remain unlabelled and every cell is systematically undersized. A second flooding pass expands the existing labels into those boundaries, but it must not expand into genuinely dark structures such as guttae, which are absent cells rather than gaps between present ones.

The two are separated by their width. Inter-cell boundaries are one to three pixels across, whereas guttae span an appreciable fraction of a cell. Morphological opening of the background mask with a disk larger than the boundary width therefore erases the boundaries and leaves the large dark regions intact:

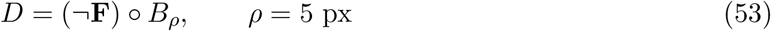

where **F** is the foreground mask, ° denotes opening and *B*_*ρ*_ a disk of radius *ρ*. Connected components of *D* smaller than 0.8× the mean labelled cell area are discarded as opening artefacts rather than true dark objects, leaving a mask of protected regions. Everything outside it is permissible territory:

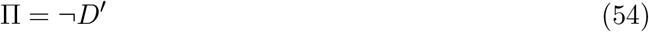

with *D*^′^ the size-filtered dark-object mask.

The labelled image **L** then serves as its own marker set for a second Euclidean distance transform watershed restricted to Π, so labels grow outward until they meet either each other or a protected dark region. Cells therefore acquire their full extent, including the boundary pixels shared with their neighbours, while guttae are left unlabelled.

Region properties are computed from the final label image, and objects smaller than 30 px are discarded as spurious fragments.

### S7.8. Pseudocode

#### Algorithm 1.

CellADE — Cornealia Mode

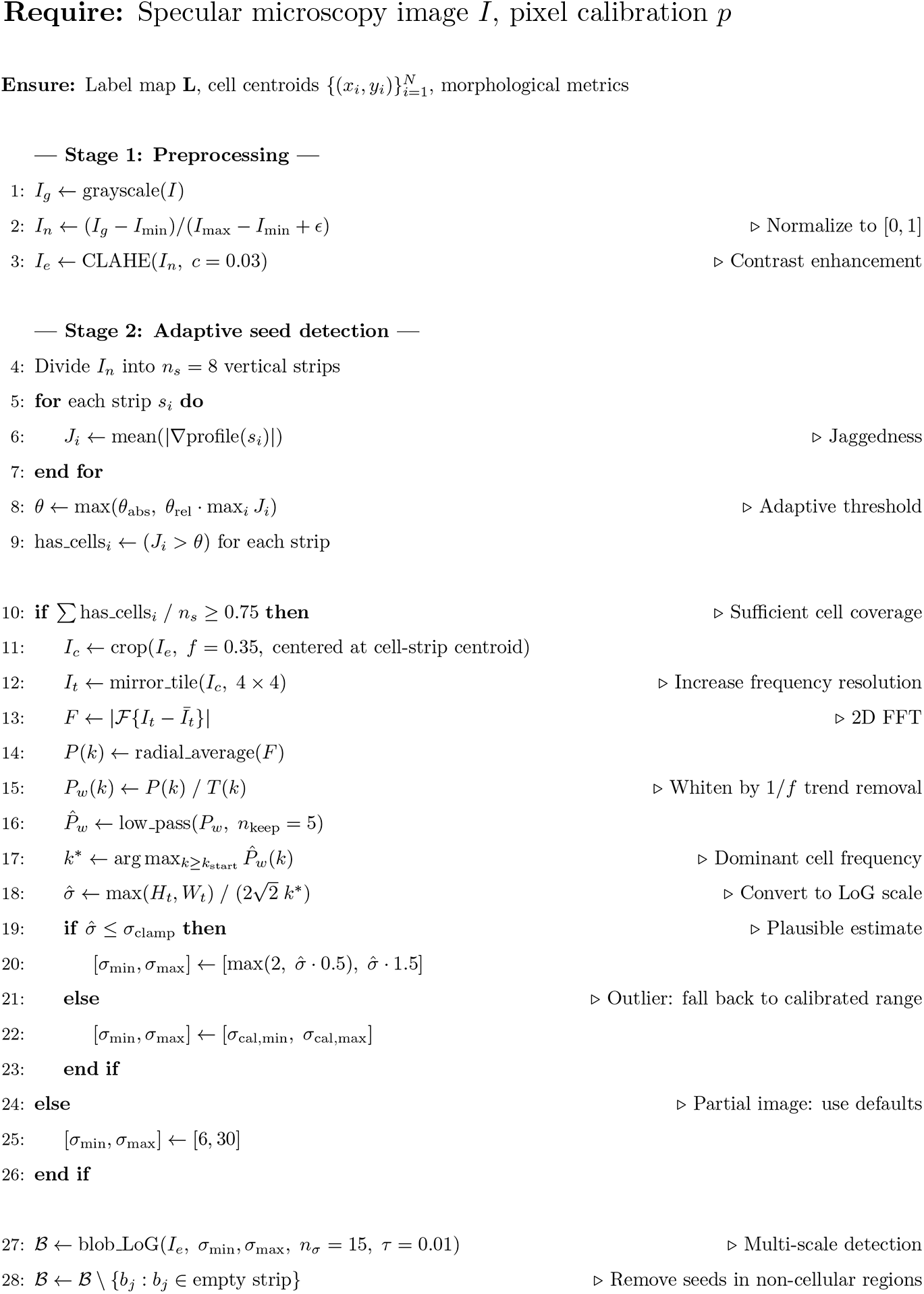
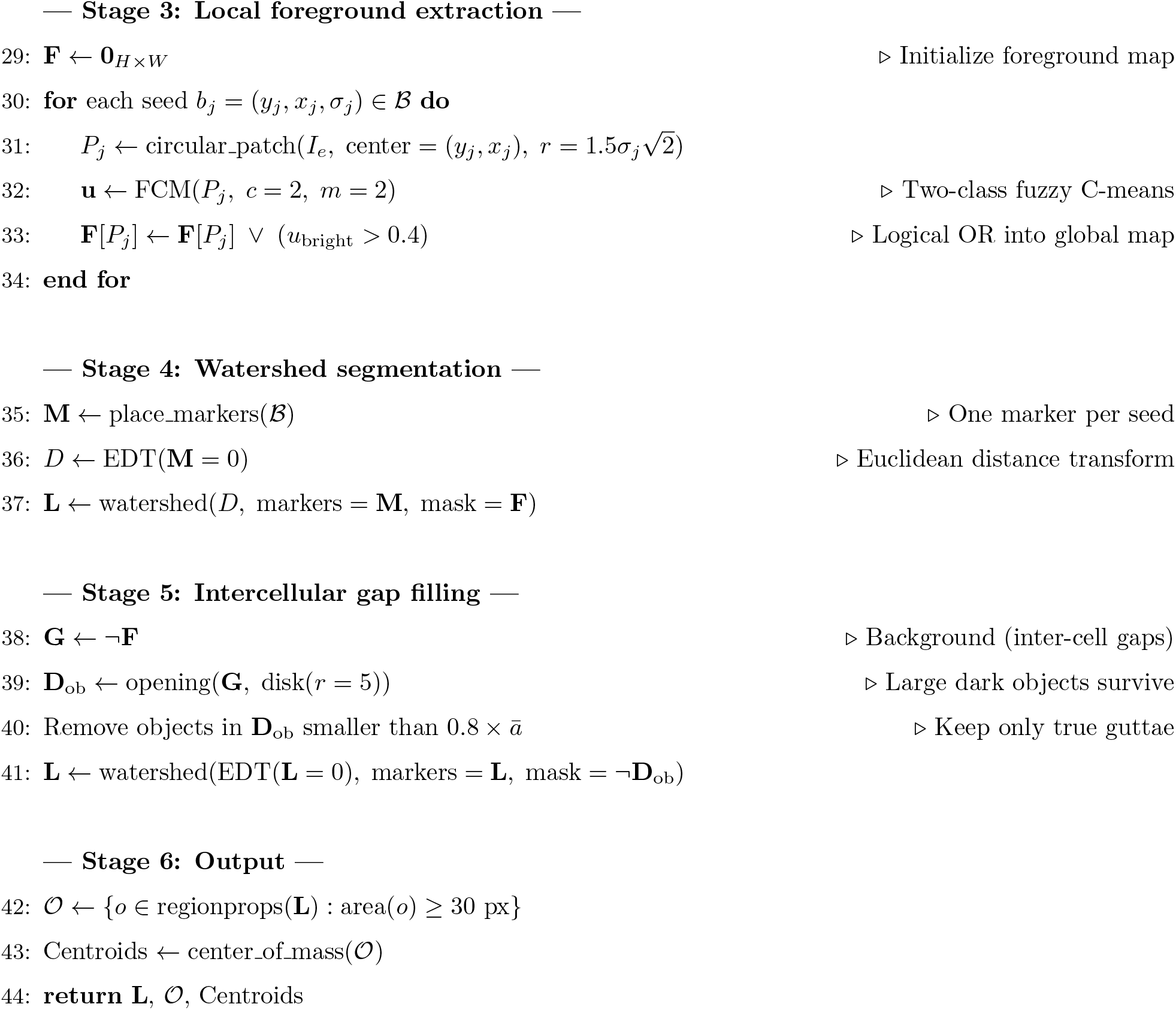

### S7.9. Parameter summary

Table S6 lists all CellADE cornealia mode parameters with their default values, physical interpretations, and tuning guidance. The three parameters most likely to require adjustment for new imaging conditions are the LoG scale range [σ_min_, σ_max_] (set automatically by FFT estimation but overridable), the LoG detection threshold, which sets how faint a seed may be, and the FCM membership threshold, which controls the foreground sensitivity.

**Table S6.** CellADE cornealia mode parameters. Parameters are grouped by pipeline stage. Bold entries indicate the three most important user-adjustable parameters.

| Parameter | Default | Physical interpretation | Tuning guidance |
| --- | --- | --- | --- |
| *Preprocessing* |  |  |  |
| `clahe_clip` | 0.03 | CLAHE contrast clip limit | Increase for very low-contrast images; decrease if enhancement saturates bright regions |
| `invert_image` | False | Inverts intensity so cells are bright on a dark background | Enable for bright-field or other modalities of opposite polarity |
| *FFT sigma estimation* |  |  |  |
| `fft_crop_frac` | 0.35 | Area fraction of center crop for spectral analysis | Increase if cells are sparse; decrease if periphery contains artifacts |
| `fft_tile_repeats` | 4 | Mirror-tile grid size ($n \times n$) | Higher values improve frequency resolution but increase computation |
| `fft_lowpass_n` | 5 | Number of Fourier coefficients retained in smoothing | Lower values give smoother envelope; too low may miss the cell peak |
| `fft_sigma_max_clamp` | 20 | Hard clamp on $\hat{\sigma}$; above this, fall back to calibrated range | Corresponds to cell radius $> 28$ px; adjust only for non-corneal tissues |
| `fft_calibrated_range` | (4.5, 11.7) | Data-driven $[\sigma_{\min}, \sigma_{\max}]$ fall-back from GT annotations | Recompute from annotated data if applying to a different instrument |
| *Strip-based ROI detection* |  |  |  |
| `fft_n_strips` | 8 | Number of vertical strips for jaggedness analysis | More strips give finer spatial resolution of cell-containing regions |
| `fft_strip_rel_thresh` | 0.4 | Jaggedness threshold relative to maximum strip | Lower values include more marginal strips; higher values are more conservative |

Table S6 continued
| Parameter | Default | Physical interpretation | Tuning guidance |
| --- | --- | --- | --- |
| `fft_strip_abs_thresh` | 0.002 | Absolute jaggedness floor, applied as a lower bound on the threshold | Raise if flat non-cellular regions are still admitted |
| `fft_strip_trim_frac` | 0.10 | Fraction trimmed from the top and bottom of each strip | Increase when image borders carry annotation or overlay artefacts |
| `fft_min_coverage` | 0.75 | Minimum fraction of cell-containing strips to proceed with FFT | Below this, FFT is skipped and defaults are used |
| *Seed detection (LoG)* |  |  |  |
| `log_min_sigma` | **6** | Minimum blob scale ≈ smallest cell radius $/\sqrt{2}$ (px) | **Primary tuning parameter.** Set automatically by FFT; override for atypically small or large cells |
| `log_max_sigma` | **30** | Maximum blob scale ≈ largest cell radius $/\sqrt{2}$ (px) | **Primary tuning parameter.** Paired with `log_min_sigma` |
| `log_num_sigma` | 15 | Number of LoG scales between $\sigma_{\min}$ and $\sigma_{\max}$ | More scales increase sensitivity to intermediate sizes but slow detection |
| `log_threshold` | **0.01** | Minimum LoG response to accept a blob | **Key detection control.** Lower detects more seeds in dim regions and low-quality images (including noise); higher is more selective |
| *Foreground extraction (local FCM)* |  |  |  |
| `fcm_mode` | local | FCM clustering strategy (local \| global \| global+local) | Local is robust to illumination gradients; global is faster but less adaptive |
| `fcm_clusters` | 2 | Number of FCM classes (foreground + background) | Fixed at 2 for binary foreground extraction |

Table S6 continued
| Parameter | Default | Physical interpretation | Tuning guidance |
| --- | --- | --- | --- |
| fcm_threshold | 0.4 | Fuzzy membership cutoff for foreground classification | **Key sensitivity control.**<br>Lower includes more borderline pixels (larger cells); higher is more selective (smaller cells) |
| *Gap filling and output* |  |  |  |
| dob_opening_radius | 5 | Disk radius for dark object exclusion (px) | Must exceed inter-cell gap width ($\sim 1\text{--}3$ px) but be smaller than guttae |
| dob_min_area_ratio | 0.8 | Minimum dark-object area, as a fraction of mean cell area | Lower to protect small guttae; raise if opening artefacts survive |
| min_area | 30 | Minimum cell area to retain (px) | Removes spurious fragments; scale with imaging resolution |

## S8. How non-square pixels are handled in Cornealia

### S8.1. Non-square pixels in the second cohort

The second cohort was imaged with a HAI Labs EB-3000xyz microscope, whose 500×450 *µ*m field of view is recorded on 640 × 480 px images. The pixels are therefore not square: 0.781 *µ*m wide and 0.938 *µ*m high, so a round cell appears 1.2 times wider than tall in the image.

### S8.2. How Cornealia handles them

Images and centroids stay in their native pixels; only distances are corrected. With pixel sizes *s*_*x*_ and *s*_*y*_, the effective pixel size 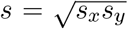 and the axis weights *a*_*x*_ = *s*_*x*_*/s* and *a*_*y*_ = *s*_*y*_*/s*, the distance between two cells is

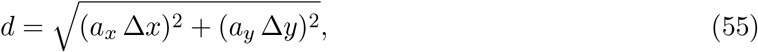

in effective pixels. This distance is used for *g*(*r*), hexagonality and the force network, and the interior margin is set from the physical size of the image. Because *a*_*x*_*a*_*y*_ = 1, each native pixel keeps its true area, so cell areas and density are unchanged. Physical units follow as *K*^*µ*m^ = and 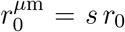For square pixels (*s*_*x*_ = *s*_*y*_), *a*_*x*_ = *a*_*y*_ = 1 and the method reduces exactly to the original.

### S8.3. Setting the pixel size in the API

The pixel size is passed as a single value for square pixels or as a pair for non-square ones (Supplementary Table S7). The pair is given as the width and height of one pixel in *µ*m, i.e. along the columns and the rows. No other setting changes: segmentation runs on the native image, and all reported quantities are returned in physical units.

**Table S7.**
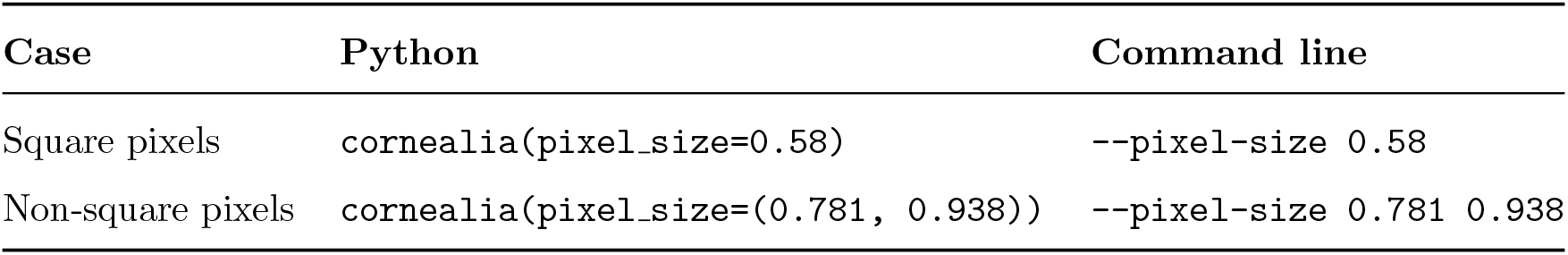
Setting the pixel size in Cornealia.

## Notes

### Competing Interest Statement

The authors have declared no competing interest.

## References

[1] Bonanno, J. A. Identity and regulation of ion transport mechanisms in the corneal endothelium. Prog. Retin. Eye Res. 22, 69–94 (2003).

[2] Edelhauser, H. F. The balance between corneal transparency and edema: the Proctor Lecture. Invest. Ophthalmol. Vis. Sci. 47, 1754–1767 (2006).

[3] Joyce, N. C. Proliferative capacity of the corneal endothelium. Prog. Retin. Eye Res. 22, 359–389 (2003).

[4] Joyce, N. C. Proliferative capacity of corneal endothelial cells. Exp. Eye Res. 95, 16–23 (2012).

[5] Bourne, W. M. Clinical estimation of corneal endothelial pump function. Trans. Am. Ophthalmol. Soc. 96, 229–242 (1998).

[6] Van Meter, W., Buckman, N., Philippy, B. & DeMatteo, J., on behalf of the Eye Bank Association of America. 2024 Eye Banking Statistical Report—Executive Summary. Eye Bank. Corneal Transpl. 4, e0042 (2025). doi:10.1097/ebct.0000000000000042

[7] Eye Bank Association of America. 2024 Eye Banking Statistical Report (EBAA, Washington, DC, 2025).

[8] Romano, V. et al. Quality assurance in corneal transplants: donor cornea assessment and oversight. Surv. Ophthalmol. 69, 465–482 (2024).

[9] McCarey, B. E., Edelhauser, H. F. & Lynn, M. J. Review of corneal endothelial specular microscopy for FDA clinical trials. Cornea 27, 1–16 (2008).

[10] Yamamoto, A. et al. A physical biomarker of the quality of cultured corneal endothelial cells and of the long-term prognosis of corneal restoration in patients. Nat. Biomed. Eng. 3, 953–960 (2019).

[11] Gain, P. et al. Global survey of corneal transplantation and eye banking. JAMA Ophthalmol. 134, 167–173 (2016).

[12] Ng, X. Y., Peh, G., Morales-Wong, F., Gabriel, R., Soong, P. L., Lin, K. H. & Mehta, J. S. Towards clinical application: calcium waves for in vitro qualitative assessment of propagated primary human corneal endothelial cells. Cells 13(23), 2012 (2024).

[13] Mehta, J. S., Kocaba, V. & Peh, G. S. A prognostic biomarker of corneal repair. Nat. Biomed. Eng. 3, 945–946 (2019).

[14] Daniel, M. C. et al. Automated segmentation of the corneal endothelium in a large set of ‘real-world’ specular microscopy images using the U-Net architecture. Sci. Rep. 9, 4752 (2019).

[15] Stringer, C., Wang, T., Michaelos, M. & Pachitariu, M. Cellpose: a generalist algorithm for cellular segmentation. Nat. Methods 18, 100–106 (2021).

[16] Schmidt, U., Weigert, M., Broaddus, C. & Myers, G. Cell detection with star-convex polygons. in MICCAI 265–273 (Springer, 2018).

[17] Peh, G. S. L., Chng, Z., Ang, H.-P., Cheng, T. Y. D., Adnan, K., Seah, X.-Y., George, B. L., Toh, K.-P., Tan, D. T., Yam, G. H. F., Colman, A. & Mehta, J. S. Propagation of human corneal endothelial cells: a novel dual media approach. Cell Transplant. 24, 287–304 (2015).

[18] Lord, S. J., Velle, K. B., Mullins, R. D. & Fritz-Laylin, L. K. SuperPlots: communicating reproducibility and variability in cell biology. J. Cell Biol. 219, e202001064 (2020).

[19] Barnett, A. G., van der Pols, J. C. & Dobson, A. J. Regression to the mean: what it is and how to deal with it. Int. J. Epidemiol. 34, 215–220 (2005).

[20] Varadhan, R., Zhu, J. & Bandeen-Roche, K. Identifying predictors of resilience to stressors in single-arm studies of pre–post change. Biostatistics 25, 1094–1111 (2024).

[21] Oldham, P. D. A note on the analysis of repeated measurements of the same subjects. J. Chronic Dis. 15, 969–977 (1962).

[22] Tu, Y.-K., Maddick, I. H., Griffiths, G. S. & Gilthorpe, M. S. Mathematical coupling can undermine the statistical assessment of clinical research: illustration from the treatment of guided tissue regeneration. J. Dent. 32, 133–142 (2004).

[23] Armitage, W. J. Assessment of corneal quality by eye banks. J. Ophthalmic Vis. Res. 6, 3–4 (2011).

[24] Bernard, A. et al. 3D images of the endothelial surface to increase accuracy of cell count in eye banks. IRBM 37, 69–75 (2016).

[25] Vidal-Villegas, B. et al. Corneal endothelial validation in the eye bank: differences in automated methods and repeatability. J. Fr. Ophtalmol. 47, 104022 (2024).

[26] Okumura, N., Kay, E. P., Nakahara, M., Hamuro, J., Kinoshita, S. & Koizumi, N. Inhibition of TGF-β signaling enables human corneal endothelial cell expansion in vitro for use in regenerative medicine. PLoS ONE 8, e58000 (2013).

[27] Hongo, A., Okumura, N., Nakahara, M., Kay, E. P. & Koizumi, N. The effect of a p38 mitogen-activated protein kinase inhibitor on cellular senescence of cultivated human corneal endothelial cells. Invest. Ophthalmol. Vis. Sci. 58, 3325–3334 (2017).

[28] Nakahara, M., Okumura, N., Nakano, S. & Koizumi, N. Effect of a p38 mitogen-activated protein kinase inhibitor on corneal endothelial cell proliferation. Invest. Ophthalmol. Vis. Sci. 59, 4218–4227 (2018).

[29] Filev, F. et al. Semiautomatic assessment of endothelial density and morphology in organ-cultured corneas—potential predictors for transplantation suitability and clinical outcome? Graefes Arch. Clin. Exp. Ophthalmol. 261, 2593–2602 (2023).

[30] Wakefield, M. J. et al. The impact of donor age and endothelial cell density on graft survival following penetrating keratoplasty. Br. J. Ophthalmol. 100, 986–989 (2016).

[31] Wilhelm, T. I. et al. Ten-year outcomes after DMEK, DSAEK, and PK: insights on graft survival, endothelial cell density loss, rejection and visual acuity. Sci. Rep. 15, 1249 (2025).

[32] Brookes, N. H. Riding the cell jamming boundary: geometry, topology, and phase of human corneal endothelium. Exp. Eye Res. 172, 171–180 (2018).

[33] Bi, D., Yang, X., Marchetti, M. C. & Manning, M. L. Motility-driven glass and jamming transitions in biological tissues. Phys. Rev. X 6, 021011 (2016).

[34] Chaithanya, K. V. S., Rozman, J., Košmrlj, A. & Sknepnek, R. Cell-level modelling of homeostasis in confined epithelial monolayers. J. Elast. 157, 29 (2025).

[35] Lie, J. T., Birbal, R., Ham, L., van der Wees, J. & Melles, G. R. J. Donor tissue preparation for Descemet membrane endothelial keratoplasty. J. Cataract Refract. Surg. 34, 1578–1583 (2008).

[36] Peh, G. S. L., Toh, K.-P., Ang, H.-P., Seah, X.-Y., George, B. L. & Mehta, J. S. Optimization of human corneal endothelial cell culture: density dependency of successful cultures in vitro. BMC Res. Notes 6, 176 (2013).

[37] Telea, A. An image inpainting technique based on the fast marching method. J. Graph. Tools 9, 23–34 (2004).

[38] Bezdek, J. C., Ehrlich, R. & Full, W. FCM: the fuzzy c-means clustering algorithm. Comput. Geosci. 10, 191–203 (1984).

[39] Meyer, F. Topographic distance and watershed lines. Signal Process. 38, 113–125 (1994).

[40] Morton, V. & Torgerson, D. J. Effect of regression to the mean on decision making in health care. BMJ 326, 1083–1084 (2003).

[41] Holcomb, J. P. Regression with covariates and outcome calculated from a common set of variables measured with error: estimation using the SIMEX method. Stat. Med. 18, 2847–2862 (1999).

## Supplementary References

[S1] Tu, Y.-K. & Gilthorpe, M. S. Revisiting the relation between change and initial value: a review and evaluation. Stat. Med. 26, 443–457 (2007).

[S2] Kelly, C. & Price, T. D. Correcting for regression to the mean in behavior and ecology. Am. Nat. 166, 700–707 (2005).

[S3] Han, S. J., Oak, Y., Groisman, A. & Bhatt, T. Traction microscopy to identify force modulation in subresolution adhesions. Nat. Methods 12, 653–656 (2015).

[S4] Haarman, S., Kim, S. Y., Isogai, T., Dean, K. M. & Han, S. J. Particle retracking algorithm capable of quantifying large, local matrix deformation for traction force microscopy. PLoS One 17, e0268614 (2022).

